# Nutrient copper signaling promotes protein turnover by allosteric activation of ubiquitin E2D conjugases

**DOI:** 10.1101/2021.02.15.431211

**Authors:** C.M. Opazo, A. Lotan, Z. Xiao, B. Zhang, M.A. Greenough, C.M. Lim, H. Trytell, A. Ramírez, A.A. Ukuwela, C.H. Mawal, J. McKenna, D.N. Saunders, R. Burke, P.R. Gooley, A.I. Bush

## Abstract

Nutrient copper supply is critical for cell growth and differentiation, and its disturbance is associated with major pathologies including cancer and neurodegeneration. Although increasing copper bioavailability in late Precambrian facilitated emergence of novel cuproproteins, their intricate regulation by this essential trace element remains largely cryptic. We found that subtle rises in cellular copper strikingly increase polyubiquitination and accelerate protein degradation within 30 minutes in numerous mammalian cell lines. We track this surprising observation to allostery induced in the UBE2D ubiquitin conjugase clade through a conserved CXXXC sub-femtomolar-affinity Cu^+^ binding motif. Thus, physiologic fluctuation in cytoplasmic Cu^+^ is coupled to the prompt degradation of UBE2D protein targets, including p53. In *Drosophila* harboring a larval-lethal knockdown of the nearly identical fly orthologue UbcD1, complementation with human UBE2D2 restored near-normal development, but mutation of its CXXXC Cu^+^ binding motif profoundly disrupted organogenesis. Nutrient Cu^+^ emerges as a trophic allosteric modulator of UBE2D activity through a structural motif whose evolution coincides with animal multicellularity.

**One Sentence Summary:** Modulation of nutrient copper impacts protein turnover and animal morphogenesis through conserved allostery of ubiquitin E2D conjugases.

**Hilights:**

- Nutrient copper supply is critical for cell growth and differentiation
- The E2D clade of ubiquitin conjugases contains a sub-femtomolar-affinity Cu^+^ binding motif
- Allosteric activation by Cu^+^ markedly accelerates protein polyubiquitination
- This sensor couples physiologic fluctuations in cytoplasmic Cu^+^ with the degradation rate of E2D targets, including p53
- This metazoan signaling mechanism is critical for *drosophila* morphogenesis

**In Brief:** Conserved allostery of ubiquitin E2D conjugases links nutrient copper signaling to protein degradation and animal morphogenesis.

Graphical Abstract

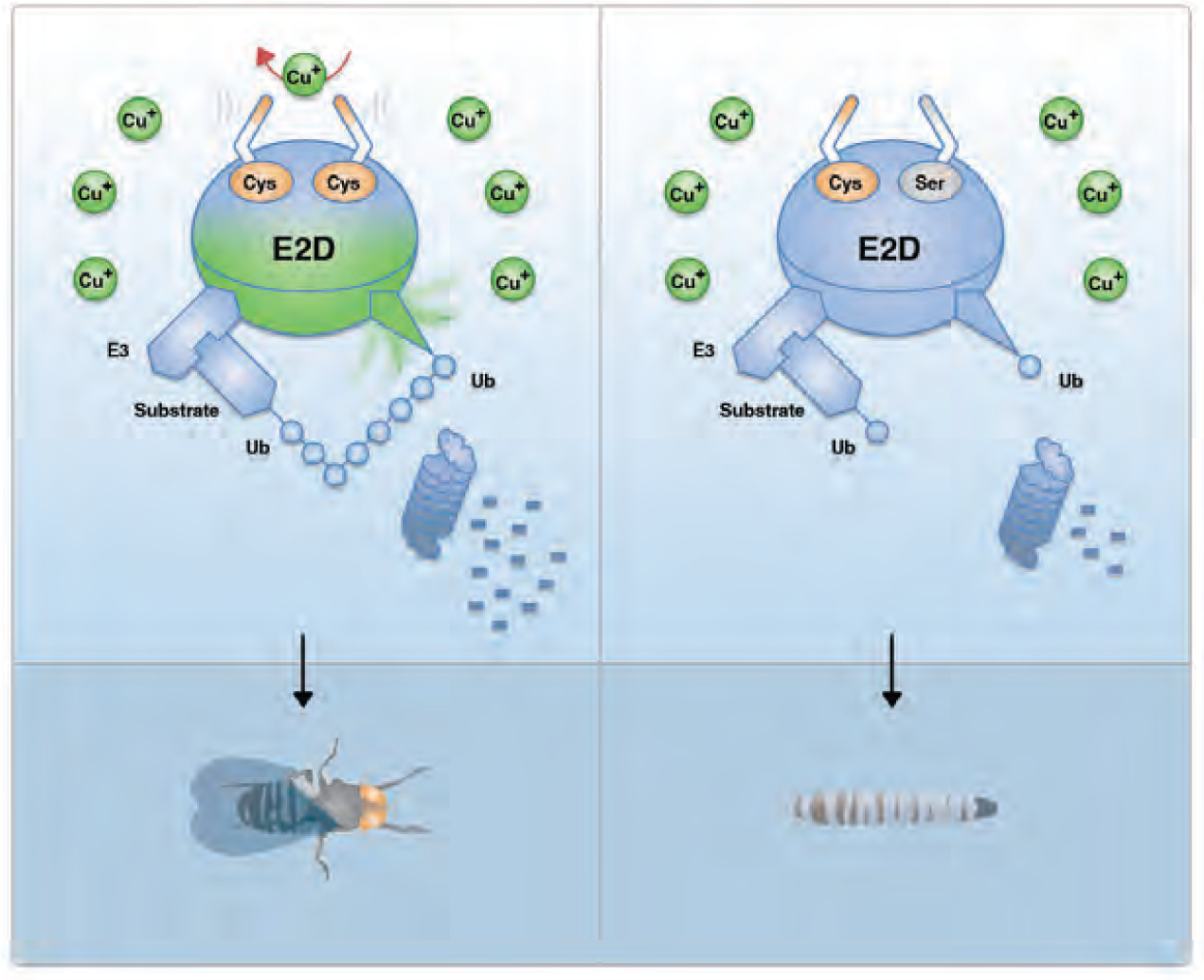

## INTRODUCTION

Copper is an indispensable nutrient in animals. While it has classically been appreciated for its essential structural and catalytic functions (*1*), a novel role for Cu^+^ has recently been proposed in cell signaling to modulate enzyme activity and cell physiology (*2*). Cytoplasmic Cu^+^ is buffered to the sub-femtomolar (fM) range by abundant glutathione (GSH) (*3*), which also maintains active site sulfhydryl residues of E2 enzymes in the reduced state, supporting the formation of protein-ubiquitin conjugates *de novo* (*4*). The ubiquitin-conjugating enzymes (E2s) are increasingly appreciated as critical players in the orchestration of ubiquitination, but the functional characterization of sequence diversity within the family, which in humans includes 40 human protein members, is rudimentary (*5, 6*). E2s can be regulated by non-covalent interactions and covalent post-translational modifications, yet the functional significance of this is still notional (*7*). A rendezvous between E2s and copper supply has not yet been reported. Here we show that cellular copper abundance strikingly transmits a signal to promote protein turnover transduced through E2D enzymes, through an evolutionarily conserved high-affinity Cu^+^-binding CXXXC motif. Cu^+^-induced allostery increases E2D conjugase activity, markedly accelerating the formation of polyubiquitination and protein turnover, which proves critical for *Drosophila* development.

## RESULTS

### Cu^+^ promotes protein ubiquitination and degradation

While investigating the impact of elevating cellular copper levels in mouse primary cortical neuronal cultures by supplementing media with low concentrations of CuCl_2_ (Figure 1A), we were surprised to witness a striking accumulation of polyubiquitinated proteins (Figure 1B). We found similar responses in different cell lines derived from hamsters (CHO), humans (HEK293T), mice (NSC34) and rats (N27), where polyubiquitination was dose-dependent (Figure S1A), in tandem with rising intracellular copper levels (Figure S1B). In line with previous reports, such low-dose copper treatment did not impair viability (Figure S2A) or increase reactive oxygen species (Figure S2B) (*8, 9*). Copper imported into the cell is known to rapidly interact with a large excess (millimolar-range) of the major cytosolic redox buffer, GSH, and is then trafficked to designated cuproproteins (*10*). Such lack of toxicity upon copper supplementation is consistent with the remarkable capacity of cytosolic GSH to hold the free cytoplasmic copper concentration below 1 fM, even in the face of extreme levels (100 µM) of copper supplementation (*3*). Supporting the conclusion that Cu^+^-enhanced polyubiquitination was not a stress-driven phenomenon, exposure of N2a cells to H_2_O_2_ oxidation could not recapitulate the blush of polyubiquitination induced by copper, and conversely, the antioxidant pyruvate did not attenuate this blush (Figure 1C). Although the proteasome inhibitor MG132 induced a similar blush of polyubiquitinated proteins (Figures 1B and S1A), copper supplementation did not inhibit proteasomal activity (Figure S2C), consistent with previous reports demonstrating proteasome inhibition only at much higher copper concentrations (*11, 12*), which we avoided.

**Figure 1.**
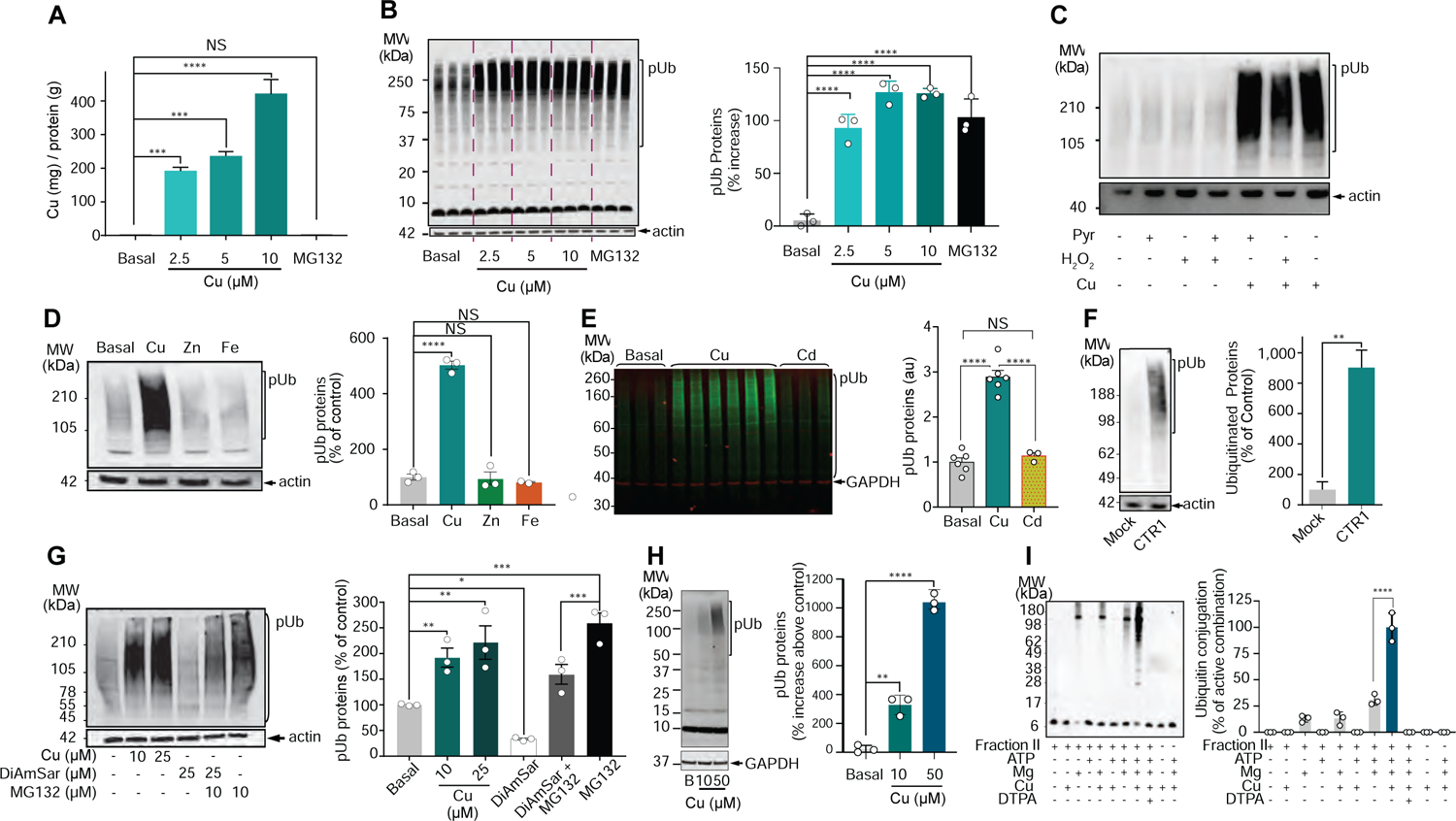
Copper promotes protein ubiquitination. (A) Cellular copper levels following supplementation. Primary cortical neurons were untreated (basal) or supplemented with CuCl_2_ (Cu; up to 10 µM) or a proteasome inhibitor (MG132; 10 μM) for 3 h in Locke’s media. Bar graph depicts means ± SEM (n = 3). \*\*\**P*<0.001, \*\*\*\**P*<0.0001, ANOVA followed by Dunnett’s test. (B) Ubiquitination response to copper supplementation in primary cortical neuronal cells. Ubiquitinated proteins (pUb) were detected by blot (P4D1 antibody). Actin was used as loading control and MG132 as positive control. Bar graph depicts means ± SEM (n = 3). \*\*\*\**P*<0.0001, ANOVA followed by Dunnett’s test. (C) N2a mouse neuroblastoma cells were exposed to the indicated reagents (Cu = 10 µM, H_2_O_2_ = 5 µM, pyruvate [Pyr] = 10 µM) for 3h in Locke’s media. pUb were detected by Western blot. (D) Primary cortical neuronal cells were incubated ± 10 µM CuCl_2_, ZnCl_2_ or FeCl_3_ for 3 h. pUb were detected by Western blot. Bar graph depicts means ± SEM (n = 3). \*\*\*\**P*<0.0001, ANOVA followed by Dunnett’s test. (E) HEK cells were incubated ± 10 µM CuCl_2_ or CdCl_2_ for 3 h in Locke’s media. pUb were detected by Western blot. Actin was used as loading control. Bar graph depicts means ± SEM (n = 3-6). \*\*\*\**P*<0.0001, ANOVA followed by Tukey’s test. (F) Levels of pUb in HEK-CTR1 (stably transfected) and HEK-mock cells. pUb were detected by Western blot. Bar graph depicts means ± SEM (n = 3). \*\**P*<0.01, student’s *t*-test. (G) pUb levels in N2a mouse neuroblastoma cells incubated ± CuCl_2_ (10 µM or 25 µM), copper chelator (DiAmSar, 25 µM), MG132 (10 µM), or DiAmSar + MG132 for 3 h. Bar graph depicts means ± SEM (n = 3). \**P*<0.05, \*\**P*<0.01, \*\*\**P*<0.001, ANOVA followed by Holm-Sidak’s test. (H) Copper promotes ubiquitination in freshly lysed HeLa cell supernatants. Supernatants were incubated in the absence (basal) or presence of CuCl_2_ (10 and 50 μM), in 200 µM GSH + 50 mM Tris-HCl (pH 7.5) for 3h. pUb were detected by Western blot. GAPDH was used as loading control. Bar graph depicts means ± SEM (n = 3). \*\**P*<0.01, \*\*\*\**P*<0.0001, ANOVA followed by Tukey’s test. (I) Copper activates polyubiquitination *in vitro*. Fraction II (the protein fraction of HeLa S3 lysate that binds to anion exchange resin), a physiologic source of E-enzymes, was incubated (37 °C, 5 h) with biotinylated ubiquitin (2.5 µM) in the presence or absence of ATP (2 mM), MgCl_2_ (5 mM), CuCl_2_ (50 µM) and DTPA (5 mM). Samples were submitted to electrophoresis under non-reducing conditions. Ubiquitin conjugates was detected by avidin-HRP binding using ECL detection system. Bar graph depicts means ± SEM from three replication experiments. \*\*\*\**P* <0.0001, student’s *t*-test. See also Figures S1 and S2.

Neither iron and zinc nor cadmium could induce a similar polyubiquitination blush (Figures 1D-E). Cuprous ions (Cu^+^) are rapidly oxidized by dioxygen (*13*), so cupric (Cu^2+^) salts are routinely used to supplement cell culture media. Some Cu^2+^ can enter the cell via DMT1 (*14*) but most Cu^2+^ is reduced at the membrane and transferred into the cytoplasm as Cu^+^ by CTR1 (*15*). Leveraging this, we stably transfected human CTR1 (*16*) into HEK293 cells, finding that this also induced accumulation of polyubiquitinated proteins (Figure 1F). Thus, a sustained rise in cytoplasmic levels of Cu^+^ increases copper-enhanced polyubiquitination. Conversely, treatment with the copper chelator DiAmSar (*17*) decreased steady-state levels of polyubiquitinated proteins in N2a cells (Figure 1G), confirming that under resting conditions, endogenous Cu^+^ promotes polyubiquitination.

To further characterize copper-enhanced polyubiquitination, we added Cu^+^ (kept reduced by supplementary GSH) to the post-centrifugation soluble fractions of freshly lysed HeLa cells. In this preparation, prominent formation of polyubiquitinated proteins (Figure 1H) was unlikely to reflect cellular responses to protein denaturation and oxidative stress. We then tested the ability of Cu^+^ to modulate the early steps of ubiquitination in a cell-free system containing E1, E2 and E3 enzymes, with biotinylated ubiquitin adduct formation as the readout. We studied Fraction II from HeLa S3 lysate, first used to map this pathway(*18*). Fraction II was incubated in 5 mM EDTA for 30 min and desalted to remove all exchangeable metal ions present. As expected, polyubiquitinated protein conjugates were detected upon re-addition of Mg^2+^ and ATP (Figure 1E). While in the absence of Mg^2+^/ATP, Cu^+^ (50 µM) did not induce ubiquitin conjugation, Cu^+^ strikingly augmented polyubiquitinated conjugation induced by Mg^2+^/ATP, while metal ion sequestration by DTPA abolished polyubiquitin conjugation (Figure 1I).

Consistent with its impact in generating polyubiquitinated proteins, copper promoted protein degradation. Following a ^35^S-cysteine/methionine pulse in primary cortical cultures, copper (10 µM) supplementation during a 30-min chase led to increased secretion of labelled low-molecular weight peptides, reflecting accelerated protein degradation (Figure 2A). Mirroring this result, pulse-chase autoradiography profiles in NSC34 (Figures 2B and S2D) and MEF (Figure S2E) cell lines showed that copper treatment accelerated the disappearance of labelled cellular proteins. Such acceleration of protein degradation mitigated the possibility that the polyubiquitination response was driven by Cu^+^-mediated inhibition of deubiquitination enzymes. Overall, Cu^+^ enhancement of protein polyubiquitination and degradation was replicated across numerous mammalian cell lines (Figure S2F), highlighting the biological relevance of this apparent signaling response.

**Figure 2.**
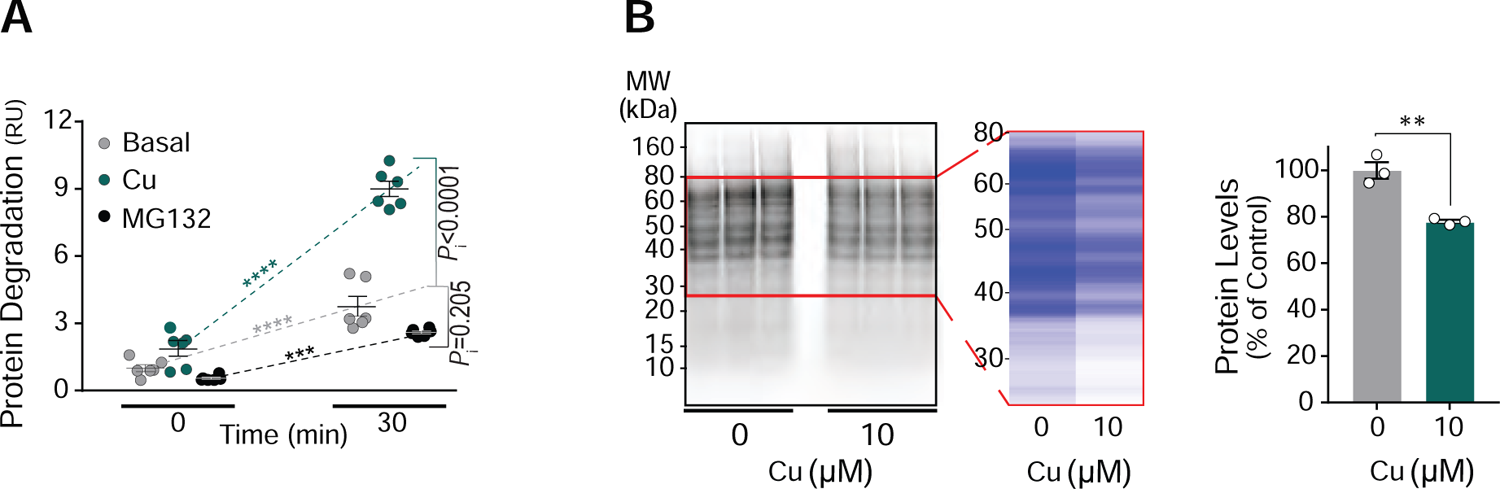
Copper promotes protein degradation. (A) Primary cortical neuronal cells were labelled with ^35^S-Cys/Met for 1 h, followed by chase in unlabeled media containing cycloheximide ± CuCl_2_ (10 µM) or MG132 (50 µM) for 30 min. The graph shows an index of protein degradation (radioactive counts from media remaining soluble after TCA precipitation, reflecting ratio of total peptide fragments to counts from total cell proteins). Data presented as means ± SEM (n = 6). \*\*\**P*<0.001, \*\*\*\**P*<0.0001, two-way-ANOVA followed by Sidak’s tests. *P*i, significance of time×medium interaction terms. (B) NSC34 cells were labelled with ^35^S-Cys/Met, followed by chase in unlabeled media containing cycloheximide ± CuCl_2_ (10 µM) for 30 min. Autoradiography protein profile upon gel electrophoresis of triplicate cell cultures is shown. The red frame encloses a region of interest, represented as an intensity heatmap for each experimental condition. Bar graph depicts means ± SEM (n = 3). \*\**P*<0.01, independent-samples *t*-test. See also Figure S2.

### p53 is a major target for Cu^+^-enhanced ubiquitination and degradation

To pursue the targets for Cu^+^-enhanced polyubiquitination, we characterized the ubiquitome (*19*) in NSC34 cells (Figure S3A and Table S1), which manifested robust increases in polyubiquitination induced by copper (Figure S1). TUBE-affinity capture of protein ubiquitin-modifications from cell lysates revealed that supplementation with 10 or 25 µM copper increased cellular pools of ubiquitin-associated proteins (Figure S3B), with 882 of these proteins reliably identified (Figures S3C-E and Table S1). Copper supplementation selectively increased the abundance of a subset (48/882) of these proteins, while signaling the suppression of a smaller subset (18/882, Figure 3A and Table S2). Both the ubiquitin-associated proteins (Figures 3B and S3F) and the respective biological pathways (e.g., regulation of apoptosis, proteolysis and cell cycle, Figures 3C and S3G; Table S3) that were enriched in NSC34 cells supplemented with copper displayed significant overlap across CuCl_2_ concentrations (10 and 25 μM), indicating that mild copper supplementation produces a coherent biological response. As polyubiquitin tagging is often followed by proteasomal degradation, the preponderance of proteasomal subunits and ubiquitination-regulating machinery (e.g., NEDD8) (*20*) among this subset of over-abundant proteins (Figure 3A) could serve as a negative feedback loop that would ultimately limit protein degradation in the face of increasing copper levels.

**Figure 3.**
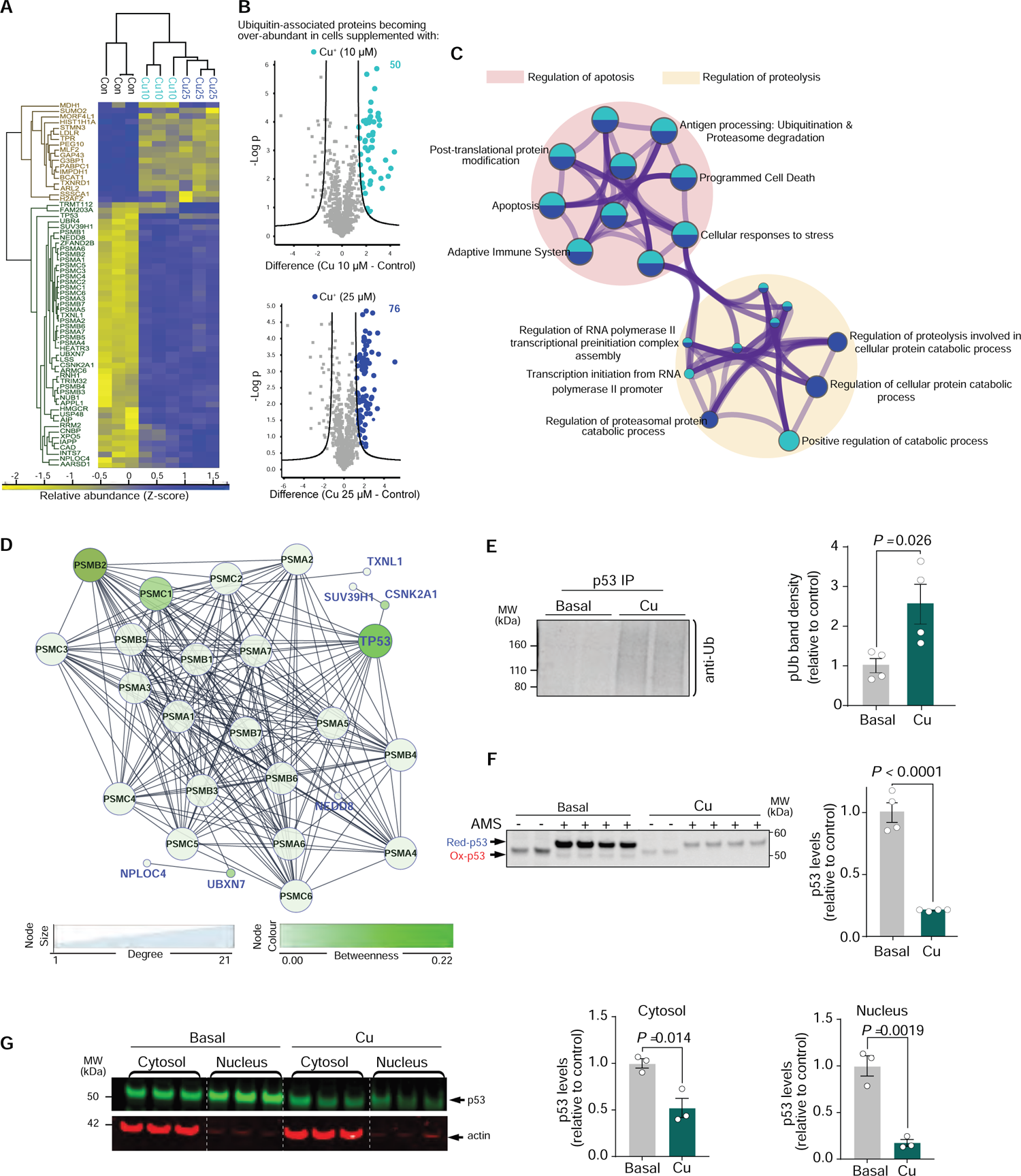
p53 is a major target for Cu^+^-enhanced ubiquitination and degradation (A) NSC34 cells were treated ± CuCl_2_ (10 µM, 25 µM) for 3 h, and harvested for ubiquitomics. Heatmap depicting the relative abundance (standardized per protein) of 66 ubiquitin-associated proteins whose abundance was altered by copper (q<0.01, ANOVA followed by permutation-based-FDR). 48 ubiquitin-associated proteins were over-abundant in Cu^+^-supplemented media, compared with 18 that were over-abundant in control media (significance based on cumulative binomial distribution). Dendrograms depict hierarchical clustering according to media conditions (columns; three replicates per condition) and protein abundance profile (rows). (B) Volcano plots depicting, for each of 882 Ub-associated proteins, the fold change (x-axis) between its abundance in lysates of NSC34 cells treated with CuCl_2_ at 10 µM (top panel) or 25 µM (bottom panel) and cells remaining in control media, and the significance of this change (y-axis, -log_10_P), based on a modified (S_0_=2) two-sample *t*-test. Cutoff curves indicate significant hits. Over-abundant proteins in CuCl_2_-supplemented media are colour-coded (light blue, 10 µM; dark blue, 25 µM) and their total number in each condition denoted. (C) Network of enriched ontology terms pied by gene counts in both conditions. To capture their inter-relationship, subsets of enriched terms from each of two distinct functional clusters have been selected and rendered as a network plot, where terms with a similarity > 0.3 are connected by edges. Node sizes are proportional to the number of hits associated with each term. Nodes are displayed as pies, where each sector (colour-coded as above) is proportional to the number of hits originated from its corresponding protein list. (D) Cu^+^-enhanced ubiquitome. A connected subnetwork of 27 proteins (nodes) that displayed over-abundance in Cu^+^-supplemented media was generated using STRING protein-protein interactions database. Size and colour correspond to node-degree and betweenness-centrality, respectively. (E) Copper promotes ubiquitination and degradation of p53. HEK293 cells were incubated ± CuCl_2_ (10 μM) for 3h, and ubiquitinated p53 was immunoprecipitated by anti-p53 antibody and blotted with anti-Ub antibody (DAKO). This blot illustrates the boost in polyubiquitinated p53 induced by copper (see Figure S4B for full blot). Bar graph depicts means ± SEM based on two replications. Significance of independent-samples *t*-test is denoted. (F) Western blot for p53 in the same experiment shows that copper promoted the degradation of p53. Bar graph depicts densitometry means ± SEM (n=4). AMS (4′-acetamido-4′-maleimidylstilbene-2,2′-disulfonic acid), which binds reduced cysteine residues in the extract, generated an electrophoretic shift, confirming that p53 cysteines were not oxidized by copper. (G) p53 Western blot of nuclear and cytosolic fractions of these cells reveals that copper induces both cytosolic and nuclear p53 loss. Actin was used as loading control. Bar graphs depict densitometry means ± SEM (n=3). Significance of independent-samples *t*-tests are denoted. See also Figures S3-S4 and Tables S1-S4.

To appreciate individual ubiquitin-associated proteins that displayed over-abundance in copper-supplemented cell lysates, we curated their corresponding protein-protein interactions (PPIs) from the STRING database (Figure S3A and Table S4). Based on network analysis of the Cu^+^-enhanced interactome (Figure 3D), p53 displayed high connectivity and the highest betweenness-centrality (Figure S4A), reflecting interactions with both proteasomal and non-proteasomal proteins. Building on the assumption that highly connected proteins with a central role in a network’s architecture are more likely to be essential to the network’s overall function(s) (*21, 22*), this analysis indicated that Cu^+^-enhanced ubiquitination of p53 could represent a physiological component of an apparent copper-signaling response.

Since our ubiquitomics analysis identified p53 as a central target for Cu^+^-enhanced ubiquitination in NSC34 cells, we tested whether diminished p53 levels upon copper supplementation could be replicated in additional cell lines. In HEK293 cells, copper supplementation (10 µM, 3h) conspicuously enhanced p53 polyubiquitination (Figures 3E and S4B) while promoting its degradation (Figure 3F). This p53 polyubiquitination was unlikely to reflect a stress response (e.g., to oxidation) because a contrasting increase in p53 levels was reported when cells had been intoxicated with 10-20-fold greater copper concentrations (*23–25*) or with a Cu-ionophore (*26*). Nevertheless, we still considered the possibility that the low levels of supplemented copper could be oxidizing cysteine residues within p53. To assess this, we used AMS, a reagent that binds non-oxidized cysteines, inducing a small electrophoretic shift (*27*). We found that AMS binding enhanced p53 immunoreactivity and confirmed that the p53 degradation induced by low-dose copper occurred in the absence of cysteine oxidation (Figure 3F). Prominent copper-induced degradation of p53 was evident in both cytosolic and nuclear fractions derived from cycloheximide-treated HEK293 cells (Figure 3G), confirming the absence of stress-associated nuclear translocation of p53 (*23*). A reciprocal accumulation of polyubiquitinated proteins and degradation of p53 was also observed in copper-supplemented mouse and human fibroblasts (Figure S4C), indicating that copper signaling might be a general mechanism to regulate p53 levels in mammalian cells.

### Cu^+^ enhances protein ubiquitination via E2D conjugases

To probe the mechanism(s) underlying the selective increase in ubiquitin tagging that was fueling protein degradation, we considered the ubiquitination cascade. As for E1, we would expect its activation (e.g., by Cu^+^) to generate a much larger list of polyubiquitination targets than we detected (Figure 3A). However, beyond E1, the breadth of Cu^+^-enhanced polyubiquitin tagging rendered an upstream target for Cu^+^ signaling (e.g., E2 activation) more likely than a downstream target (e.g., activation of many E3s). While E3s focus the targeting, E2s offer unique contribution to the target specificity of polyubiquitination (*5, 7, 28, 29*). p53 dominated the functional network of the Cu^+^-enhanced ubiquitome (Figure 3D) and was subsequently validated as a target for Cu^+^-enhanced polyubiquitination in several cell lines (Figures 3E and Figures S4B-C), positioning the E2s that preferentially target it as candidates for Cu^+^ enhancement. Among characterized E2 clades (*30*), the E2D (UBE2D; Ubch5) clade has a prominent role in promoting polyubiquitination of p53 (*31*). Examining the STRING PPI database (Figure S3H), 70 of the 882 ubiquitin-associated proteins were predicted to interact with at least one of the four paralogues (E2D1-4) of this clade (Figure 4A and Table S5). Cu^+^-enhanced ubiquitin-associated proteins were indeed highly enriched among E2D targets compared to non-E2D targets (34% vs 3%).

**Figure 4.**
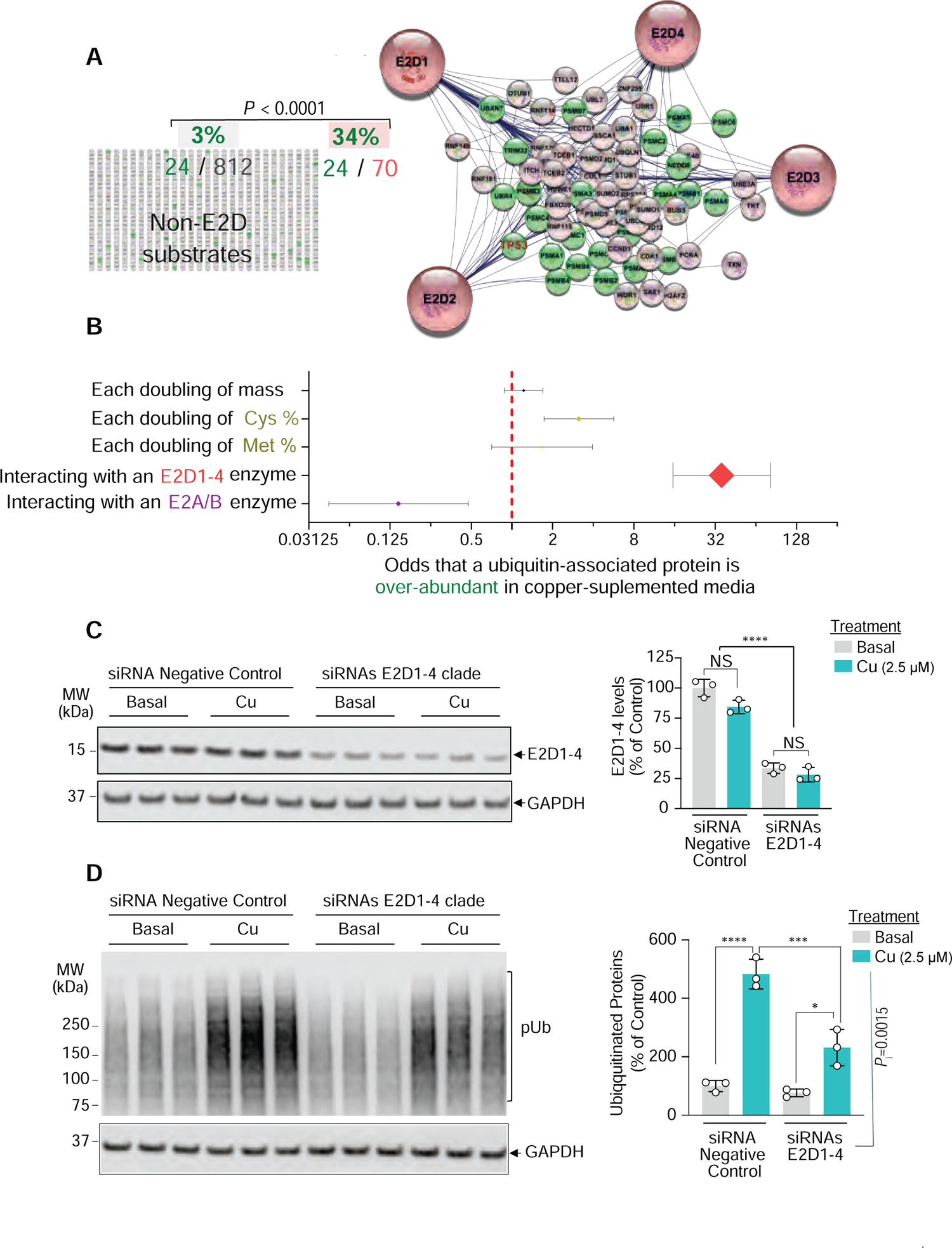
Cu^+^ enhances protein ubiquitination via E2D conjugases (A) Based on ubiquitomics data (Figure 3A), enrichment analysis comparing the proportion of proteins displaying Cu^+^-enhanced over-abundance (green nodes) among STRING-predicted E2D1-4 substrates (right) versus the corresponding proportion among non-E2D1-4 substrates (left side). Significance based on Fisher’s exact test. (B) Effects of protein mass, redox-sensitive amino acids, and being an E2D1-4 or E2A/B substrate, on Cu^+^-enhanced ubiquitination. Odds (*x*-axis, log_2_-scale) ± 95% CI of a ubiquitin-associated protein becoming over-abundant upon copper supplementation were estimated for each doubling of protein mass or cysteine/methionine content, or for being an E2D1-4 or E2A/B substrate. Diamond sizes reflect Wald-statistic of each predictor, based on a multiple logistic regression model. (C) Following 48h incubation with control or E2D1-4-targeting siRNA, HEK293 cells were exposed to non-supplemented (basal) or CuCl_2_-supplemented (Cu 2.5 µM) Locke’s media for 3 h. E2D1-4 levels were detected with an antibody that recognizes the full E2D clade. Bar graphs depict means ± SEM (n = 3). NS, non-significant, based on two-way-ANOVA followed by Bonferroni’s tests. \*\*\*\**P*<0.0001, main effect of siRNA. (D) Ubiquitinated proteins (pUb) in same experiment were detected by Western blot (P4D1 antibody). GAPDH was used as loading control. Bar graphs depict means ± SEM. \**P*<0.05, \*\*\**P*<0.001, \*\*\*\**P*<0.0001, two-way-ANOVA followed by Bonferroni’s tests. *P*i, significance of siRNA×Cu interaction term. See also Figures S3, S5 and Table S5.

Was copper sensitivity observed with targets of other E2 clades? We examined targets of the E2A/B (Rad6A/B) clade, as it is also implicated in p53 ubiquitination (*32, 33*). Copper supplementation yielded a minor enrichment of ubiquitin-associated proteins among the 47 identified E2A/B targets (Figure S5A), yet ∼70% of E2A/B targets could also be tagged by E2Ds (Table S5). Controlling for joint substrates, interaction with an E2D enzyme was the most prominent predictor of copper enhancement for a given ubiquitin-associated protein (Table S5). The odds of Cu^+^-enhanced ubiquitination for a protein handled by E2Ds were ∼35-fold greater than proteins handled by other E2s, even after controlling for protein mass and content of the redox-sensitive amino acids cysteine and methionine (Figures 4B and S5B). As E2D conjugases promote substrate (e.g., p53) *poly*ubiquitination which signals proteasomal degradation (*31*), enrichment of E2D targets among the Cu^+^-enhanced ubiquitome is consistent with the observed enhancement of protein degradation by copper (Figures 2A-B and S2D-E). Unlike E2Ds, UBE2A/B conjugases promote substrate (e.g., p53) *mono*ubiquitination to regulate activity and subcellular localization (*34, 35*). Thus, enrichment of E2D (as opposed to E2A/B) targets is consistent with p53 levels decreasing simultaneously in cytosol and nucleus upon copper supplementation (Figure 3G). Taken together, these results led us to hypothesize that Cu^+^-enhanced (poly)ubiquitination might be explained by augmented E2D activity.

To evaluate the contribution of E2Ds to copper-promoted ubiquitination in intact cells, we performed a functional experiment using a commercially available knockout (KO) cell line. E2D1-4 are paralogues (Figure S5C), and ablation of any one paralogue in Hap1 cells does not impair viability. We chose to examine the effect of copper on an E2D2 KO cell line, as E2D2 has been the most extensively characterized and is the main conjugation enzyme for p53 (*31*). Like all other cell lines tested, copper induced a polyubiquitination blush in the Hap1 cells, but this blush was attenuated in the E2D2 KO cells compared to parental controls (Figure S5D). Consistent with this, Cu^+^-induced protein degradation was also diminished in the E2D2 KO cells (Figure S5E).

These results impelled us to evaluate the overall contribution of the E2D clade to copper-promoted ubiquitination in cells. We reduced the expression of all four E2D paralogues by using small interfering RNA (siRNA)-mediated knockdown and examined three different human cell lines (HEK, Figures 4C-D; HepG2, Figures S5D-E and HeLa, Figures S5F-G). 48 hours post-transfection, siRNA efficiently suppressed (∼75%) E2D1-4 protein expression across cell lines (Figures 4C and S5F,H). Subsequently, copper (2.5-5 µM) was added to the cultures, which were then harvested after 3 hours of incubation. In control-siRNA-transfected cells copper treatment promoted polyubiquitination, yet E2D1-4 suppression markedly attenuated this effect was (Figures 4D and S5G,I). The viability of the cells at the time of harvesting was not affected by the various interventions (Figure S5J). These findings highlight a key role for the E2D clade in mediating the promotion of polyubiquitination by copper.

### Sub-femtomolar-affinity Cu^+^ binding allosterically activates E2D2

To simulate cytoplasmic conditions in a cell-free assay, we appreciated that although total intracellular copper is ∼10 µM, it is largely bound to abundant proteins such as metallothioneins and SOD1 and trafficked through affinity gradients (*10*). Cytoplasmic exchangeable copper is maintained predominantly in its reduced (Cu^+^) form, and its maximum free concentration is typically kept in the sub-fM range, by exchange with millimolar concentrations of GSH (*3, 36*). By mimicking the physiological excess of GSH, we established a cell-free functional assay suitable for assessing Cu^+^-interactions at ∼fM concentrations (Figure S6A). Under these conditions, Cu^+^ markedly promoted the activity of recombinant human E2D2, as measured by the formation of ubiquitin adducts resistant to reducing conditions (Figure 5A).

**Figure 5.**
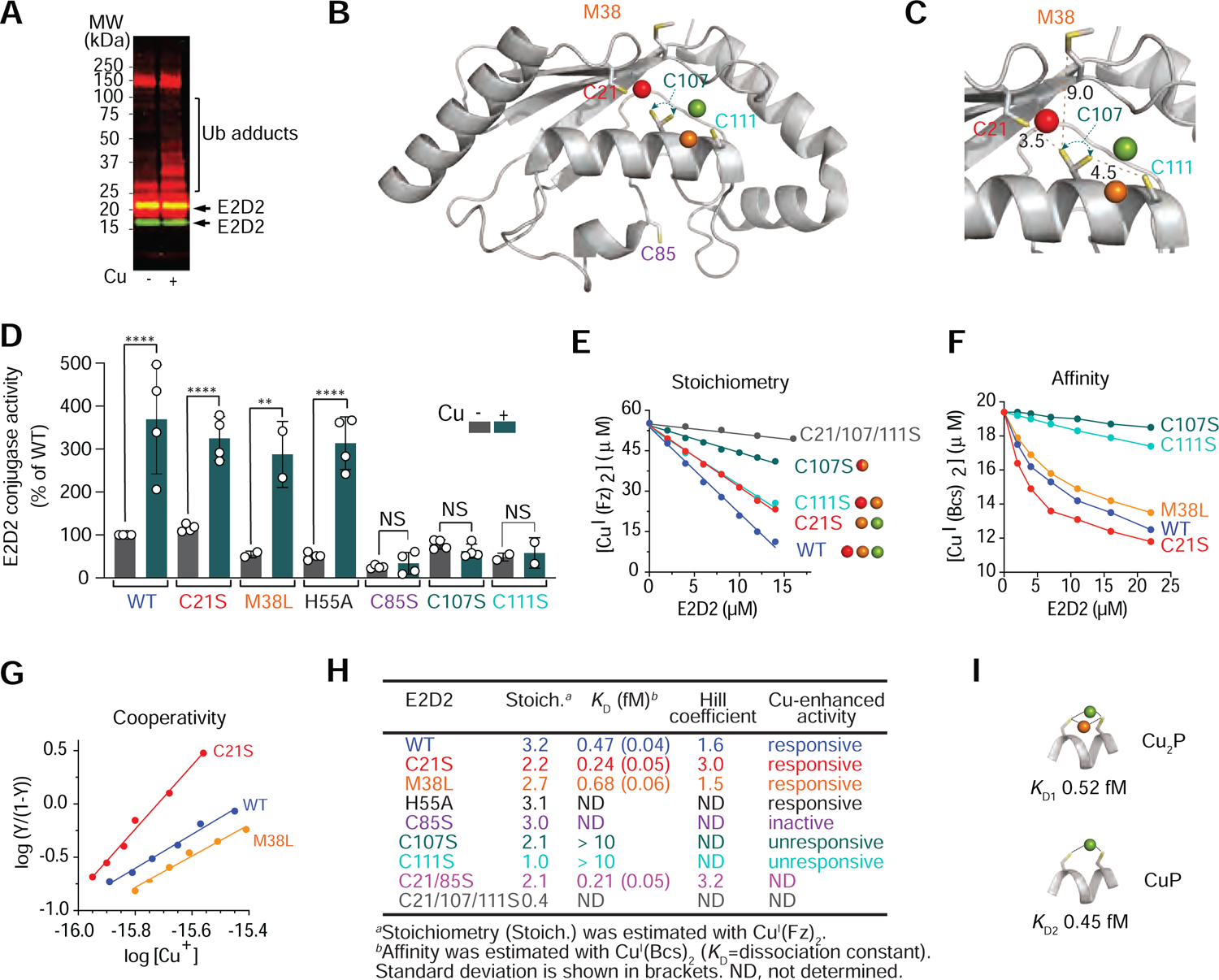
Sub-femtomolar-affinity Cu^+^ binding allosterically activates E2D2 (A-D) Assessing which putative Cu^+^ binding sites mediate enhanced E2D2 conjugase activity. (A) Biotin blot of cell-free conjugation activity of wildtype (WT) E2D2. The enzyme was incubated ± CuCl_2_ (5 µM) in the presence of GSH (200 µM), UBA1 (100 nM), MgCl_2_ (5 mM), ATP (5 mM) and biotinylated-Ub (2.5 µM) in Tris-HCl (50 mM, pH 7.5) at 37 °C. Reaction was stopped after 1 h by adding TCEP (50 mM). Ubiquitin adducts and E2D2 were detected by Streptavidin-680IR (red signal) and Western blot (green-yellow signal), respectively. Increased ubiquitin tagging (ubiquitin adducts ranging 25-100 kDa) indicated enhanced conjugase activity. (B) Crystal structure of E2D2 (pdb: 2ESK). Coloured spheres represent putative sub-pM affinity Cu^+^-binding sites. Sidechains of conserved Cys/Met residues are shown as sticks. The sidechain of C_107_ is shown as two rotamers, reflecting flexibility of the distance between key Cu^+^ ligands. (C) Zoom-in view of the high-affinity cooperative Cu^+^-binding region, highlighting distances (Å; yellow dashed lines) between the proposed ligands. (D) Bar graphs depict mean ± SEM ubiquitin conjugase activity of WT (reference= 100 % activity), or C21S, M38L, H55A, C85S, C_107_S and C111S E2D2 mutants (5 µM) in the absence (grey bars) or presence (green bars) of CuCl_2_ (5 µM), under the same conditions described above. \*\**P*<0.01, \*\*\*\**P*<0.0001, independent-samples *t*-tests. (E-H) Quantifying stoichiometry, affinity and cooperativity of high-affinity Cu^+^-binding to E2D2. (E) Quantifying Cu^+^-binding stoichiometry with the pM-affinity probe Cu^I^(Fz)_2_ under non-competitive conditions. Sphere colours correspond to sub-pM-affinity Cu^+^-binding sites proposed in panels B-C. (F) Quantifying Cu^+^-binding affinity of WT E2D2 and selected mutants with the sub-fM-affinity probe Cu^I^(Bcs)_2_ under competitive conditions. The C_107S_ and C111S mutants compete only weakly for Cu^+^-binding under these conditions. (G) Quantifying cooperativity for the sub-fM-affinity Cu^+^-binding detected by the Cu^I^(Bcs)_2_ probe. Slopes of response curves for WT and selected mutants represent Hill coefficients. (H) Table summarizing Cu^+^-binding properties and Cu^+^-enhanced activity of WT and mutant E2D2 proteins. (I) Two distinct Cu^+^-binding states (CuP and Cu2P) proposed by our NMR data (See also Figure 6 and Figure S6D), alongside their corresponding Cu^+^ *K*_D_ values. See also Figure S6 and Table S6.

Focusing on E2D2, we adopted a site-directed mutagenesis strategy to map the putative Cu^+^-binding site responsible for Cu^+^-enhanced polyubiquitination. Sequence inspection of the human E2D clade (Figure S5A), and structural inspection of E2D2 (UbcH5b, pdb: 2ESK, Figure 5B-C), identified two putative Cu^+^-binding regions: i) three cysteines (C21, C107 and C111) ± M38, and ii) two histidines (H32 and H55). Using the functional cell-free assay, all three non-catalytic-cysteine E2D2 mutants (C21S, C_107S_ and C111S) retained basal ubiquitin-conjugation activity (Figures 5D and S6B), consistent with a recent report focusing on a triple C21A/C107A/C111S mutant (*20*). However, while both C107 and C111 were critical for Cu^+^-enhancement of E2D2 conjugation activity, C21, M38 and H55 were not (Figures 5D and S6B).

In solutions where free Cu^+^ was kept at picomolar (pM) concentrations by buffering with excess ferrozine (Fz) ligand (*37*), E2D2 bound three Cu^+^ ions (Figures 5E,H and S6C). This binding stoichiometry remained unchanged for H55L and C85S mutants (Figure 5H), indicating that the putative binding site of H32/H55 was too weak to bind Cu^+^ under these conditions, and that the active site C85, as expected, was not involved in Cu^+^ binding. Both C21S and C111S mutants could bind two Cu^+^ ions only, whereas the C_107S_ mutant could bind just one, and a triple mutant (all three Cys → Ser) retained less than 0.4 Cu^+^ on average (Figure 5E,H). The saturated Cu^+^ stoichiometry of the M38L mutant was only slightly less than that of wild-type (WT, Figure 5H). Consequently, the Cu^+^-binding sites with sub-pM affinities are confined to the perimeter defined by C21, C107, C111 and M38 only (Figure 5C).

Next, Cu^+^ affinities of WT E2D2 and the C21S, C21/85S, C_107S_, C111S and M38L mutants were quantified with the chromophoric probe Cu^I^(Bcs)_2_ under anaerobic conditions. This probe, which buffers free Cu^+^ in the sub-fM range (*37*), detected that WT E2D2 competed for binding of two Cu^+^ ions cooperatively with an average *K*_D_ ∼0.5 fM and a Hill coefficient of ∼1.6 at pH 7.4 (Figures 5F-H). Mutation of M38L resulted in only a slight loss of affinity. C21S (or C21/85S) mutation increased both binding affinity and cooperativity, presumably due to the loss of the third weak site associated with C21. In contrast, mutation of either C_107S_ or C111S markedly attenuated Cu^+^-binding affinity. Thus, the C_107_XXXC_111_ region, a consensus motif for Cu^+^-coordination (*38*), is the sub-fM Cu^+^ binding site of E2D2 (Figure 5I).

### Cu^+^ binding at the C_107_XXXC_111_ motif of E2D2 triggers conformational changes that transduce to the active site vicinity

To assess the conformational changes induced by high-affinity Cu^+^-binding, we examined two protein variants, E2D2^C21/85S^ and E2D2^C111S^ using 2D NMR spectroscopy (Figure 6 and Table S6). C21S mutation increased the protein solubility in the presence of Cu^+^ at the protein concentrations required for the NMR experiments, but the mutant retained both high-affinity Cu^+^-binding (Figure 5F) and Cu^+^-enhanced conjugase activity (Figure 5D). Compared to E2D2^C21S^, in the double-mutant E2D2^C21/85S^ Cu^+^-binding affinity and stoichiometry remained unchanged (Figure 6H), disqualifying Cu^+^ interactions with the active site C85. In contrast, C111S mutation ablated both high-affinity Cu^+^-binding (Figure 5F) and Cu^+^-promoted enzyme activity (Figure 5D). Experiments were conducted in a cytosolic-mimicking KPi buffer supplemented with mM GSH, which limits free Cu^+^ to sub-fM level through cooperative assembly of a Cu^I^ (GS) cluster.

**Figure 6.**
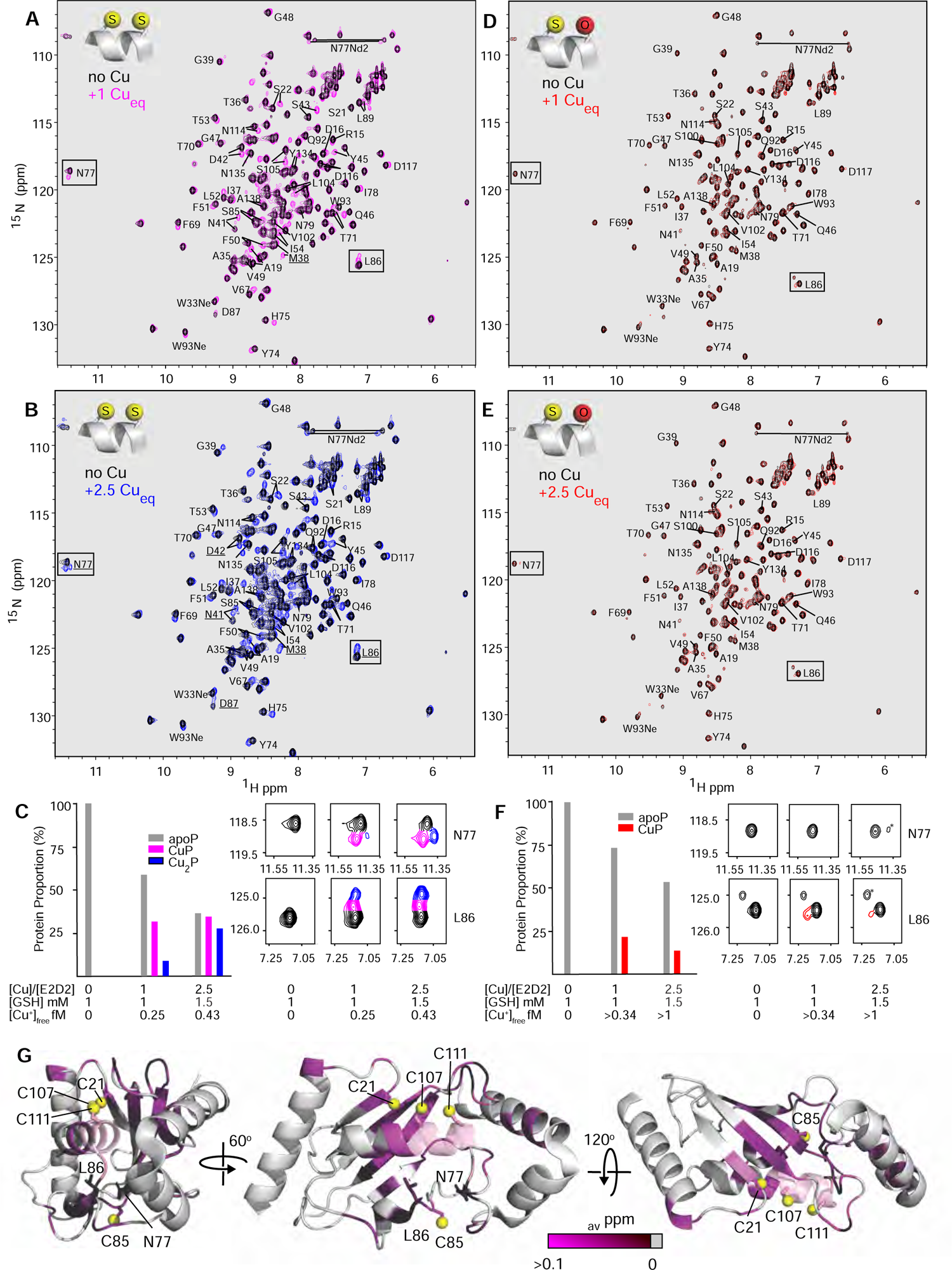
Cu^+^ binding at the C107XXXC111 motif of E2D2 triggers conformational changes that transduce to the active site vicinity (A-B) Overlay of 2D ^1^H-^15^N HSQC spectra of 0.2 mM E2D2_C21/85S_ without Cu^+^ (black) and with 0.2 mM Cu^+^ (magenta, A) or 0.5 mM Cu^+^ (blue, B) in 100 mM KPi buffer (pH 7.4), 2.0 mM NH2OH, 1.0 mM (A) or 1.5 mM (B) GSH and 5% D_2_O. Resonances that show Cu^+^-induced chemical shifts and could be assigned to CuP species in the presence of 0.2 mM Cu^+^ are indicated in both spectra. These new resonances increased in intensity and/or gave rise to a resolved third peak on the addition of 0.5 mM Cu^+^. Several residues showing these three distinct peaks assigned to apoP, CuP and Cu_2_P species are underlined and include M38, N41 and D42 near the Cu^+^-binding site and N77, L86 and D87 near the active site. The resonances for I106 to D112 were excluded as reassignment was impossible, although these resonances did show significant intensity losses in the presence of Cu^+^ (Figure S6F). Notably, the NH_2_ sidechain of N77 and the NH indoles of W33 and W93 also show Cu^+^-induced chemical shift changes. (C) Changes to chemical shift and peak intensity for two well-resolved resonances, assigned to N77 and L86 (spectral expansions in right-hand panels) located near the active site and distant from the Cu^+^-binding site. The left-hand panels show the average peak-height changes for the three resonances assigned to the apoP, CuP and Cu2P species (see Table S6 for details). (D-E) Overlay of ^1^H-^15^N HSQC spectra of 0.2 mM E2D2_C111S_ without Cu^+^ (black) and with 0.2 mM Cu^+^ (red, D) or 0.5 mM Cu^+^ (red, E) in 100 mM KPi buffer (pH 7.4), 2 mM NH2OH, 1 mM GSH and 5% D_2_O. The resonances for C21, D42, C85, I88, S111, D112, I119, A124 and Y127 of E2D2_C111S_ were excluded as they could not be unambiguously identified. For E2D2_C111S_ there was little Cu^+^-induced variation in chemical shift. However, upon copper addition E2D2_C111S_ became less stable (especially at 2.5:1 [Cu]/[E2D2]), and its precipitation over the experiment course resulted in the loss of resonance intensity. (F) Histogram of peak intensity changes for L86 and spectral expansions for N77 and L86, equivalent to panel C. A weak Cu^+^-induced peak was observed for L86 at 1:1 [Cu]/[E2D2] that weakened at 2.5:1 but retained similar proportion relative to that of apoP. The loss of intensity is consistent with ∼10% and ∼40% precipitation for 1:1 and 2.5:1 ([Cu]/[E2D2]), respectively (Table S6). The peaks marked (*) are contaminants that were observed in the absence of Cu^+^. (G) Cu^+^-induced chemical shift changes in panel A mapped onto the structure of E2D2 (pdb: 2ESK), demonstrating how conformational changes induced by Cu^+^ transduce to the active site vicinity. The largest chemical shift differences (Δδ_av_ > 0.1 ppm) are in magenta with smaller shifts shown as progressively darkened. Residues I106 to D112, whose resonances displayed significant intensity changes but could not be reassigned, are shown in transparent pink. Values for chemical shift changes are given in Figure S6E. The sulfur atoms of the Cu^+^-binding ligands C21, C107 and C111 and the active site C85 are shown as yellow spheres. The sidechains of N77 and L86 are also indicated. See also Figure S6 and Table S6.

Similar to WT enzyme spectra (*39*), Cu^+^-free samples of both E2D2 mutants exhibited well-dispersed ^1^H-^15^N HSQC spectra (Figure 6A,D), confirming minimal impact of the point mutations (C21S, C85S, C111S) on 3D structure. Addition of Cu^+^ to the control mutant (E2D2^C21/85S^), with free Cu^+^ buffered by GSH to sub-fM levels, induced conspicuous ^15^NH chemical shifts of many residues including the well-resolved resonances of N77 and L86 in the active-site region (Figure 6A-C). E2D2^C21/85S^ maintained solubility as the Cu^+^:protein ratio was increased from 1:1 to 2.5:1. In contrast, addition of Cu^+^ to E2D2^C111S^ (1:1) under comparable conditions caused negligible change of the overall ^1^H-^15^N HSQC spectrum (Figure 6D,F), except for a few resonances that appeared to have derived some minor new features (e.g., L86 with ∼22% protein proportion). Increasing the Cu^+^: E2D2^C111S^ ratio to 2.5:1 surpassed the buffering capacity of GSH (Table S6) and led to visible protein precipitation with detectable loss of overall spectral intensity (Figure 6F). Nonetheless, the total spectrum of the remaining soluble E2D2^C111S^ was largely unchanged (Figure 6E-F). These experiments demonstrated that: (i) Possessing an intact C_107_XXXC_111_ motif, E2D2^C21/85S^ competed with GSH for Cu^+^ effectively, but consistent with its attenuated binding affinity (Figure 5F), E2D2^C111S^ competed only weakly under the same conditions; (ii) high-affinity Cu^+^-binding at the C_107_XXXC_111_ motif led to considerable overall tertiary conformational change that might be the basis for Cu^+^’s allosteric effect; (iii) the weak Cu^+^-binding to E2D2^C111S^, which likely involves C21 as a coordinating residue, led to protein denaturation, which, we hypothesize, might be intended to abort E2D activity in the face of excess copper.

The well-resolved NH resonances of N77 and L86 residues, proximal to the active site C85 but distant from the high-affinity Cu^+^-binding motif C_107_XXXC_111_ (Figure 6G), were used for protein speciation analysis. Two discernible allosteric changes induced by Cu^+^-binding at this motif were quantified by the peak height of each species as a fraction of the sum of the peak heights of all protein species (Figures 6C and S6D; Table S6): (i) upon addition of 0.5-equivalent of Cu^+^, 24% of the N77 and L86 resonance peak heights in the E2D2_C21/85S_ apo-protein (apoP) shifted to another discrete state we defined as “CuP”, as well as to a second, less abundant (∼2%) state we defined as “Cu_2_P”; (ii) titrating in more Cu^+^ converted increasing fractions of apoP to CuP and Cu2P, with the latter notable in the higher Cu^+^ concentrations, consistent with the sequential binding of two Cu^+^ ions to the same C_107_XXXC_111_ motif (Figure S6D). Detailed protein speciation analysis and estimation of the associated free Cu^+^ concentration under each sample condition allowed derivation of two Cu^+^ dissociation constants, *K*_D1_ = 0.52 ± 0.02 fM and *K*_D2_ = 0.45 ± 0.02 fM at pH 7.4, which described this sequential Cu^+^-binding process (Figure 5I and Table S6). Notably, both *K*_D_ values were sub-fM, differed only marginally, and matched the average *K*_D_ = 0.21 ± 0.05 fM determined with Cu(Bcs)_2_ at same pH (Figure 5H). This rationalizes our interpretation that both CuP and Cu_2_P species are in equilibrium with apoP as additional Cu^+^ is loaded (Figure S6D), and reprises the 2:1 Cu^+^-binding stoichiometry of the high-affinity di-Cys metal-binding motif in the N-terminal domains of *A. thaliana* HMA2/4 proteins (*40*).

Resonances of CuP species that could be assigned and demonstrated resolvable chemical shift changes from apoP were scattered throughout the total spectra (Figure 6A). These were mapped onto the primary protein sequence (Figure S6E) and the reported 3D structure (Figure 6G). The large chemical shift changes triggered by high-affinity Cu^+^-binding at C_107_XXXC_111_ radiated to the proximity of the E2D2 active site (C85), possibly through a pathway across the four antiparallel β-strands (Figure 6G) which comprise a regulatory backside-binding site (*41*). The well-resolved peptide NH of N77 and L86, and the sidechain NH_2_ of N77, showed clear chemical shift changes on titration with Cu^+^ (Figure 6A-C). Considering that N77 forms a critical groove that tethers ubiquitin tightly to E2, so that it is primed for nucleophilic attack (*42, 43*), the NMR shift of N77 and nearby residues highlights the allosteric relevance of Cu^+^-binding at C_107_XXXC_111_. Together with the structure-activity relationships (Figure 5D-H), our findings indicate that the high-affinity Cu^+^-binding coordinated by C107 and C111 mediates the allosteric activation of E2D2. Notably, the sequential Cu^+^-binding of apoP → CuP → Cu_2_P triggers distinct chemical shift changes for several residues near the Cu^+^-binding site including M38, N41 and D42 and also transmits allosterically to distant residues near the C85 active site, including N77, L86 and D87 (Figure 6A-B). Therefore, Cu^+^-loading of the binding site can induce augmented levels of remote structural modification at the active site vicinity and, thus, tunable levels of allosteric regulation.

In the presence of Cu^+^, the NH resonances of the XC_107_XXXC_111_X motif (I106 to D112) could not be reassigned in the triple resonance experiments. No Cα/Cβ connectivities were detected that could be assigned to this region for either the apoP or CuP species, even though the NH resonances of S108, L109, L110 and C111 were well resolved. The lack of Cα/Cβ connectivities implies line-broadening that we attributed to a facile Cu^+^-exchange process (with GSH) mediated by two sub-fM *K*_D_ interactions that shuttled the protein rapidly between three states (apoP, CuP, Cu_2_P, Figure S6D). However, peak intensities of residues 108-111 were significantly attenuated (30 to 40% of the original intensity; Figure S6F), similar to other resonances affected by Cu^+^, and consistent with Cu^+^-inducing chemical shifts. As these residues constitute E2D2’s α-2 helix, against which the I44 hydrophobic patch of ubiquitin packs, structural modification of this helix upon Cu^+^-binding could enhance enzyme activity by stabilizing ubiquitin in the closed conformation (*41*).

Transferring ubiquitin from E2s to their target(s) usually involves a complex with an E3 ligase. To assess whether the proposed sub-fM affinity Cu^+^-binding site could be accessible in this context, we inspected the structural model of a characterized E2-E3 dimeric complex (E2D2-Ub-MDM2)_2_ (Figure S6G(i)) (*43*). The structure revealed that the C107XXXC111 motif was located on the opposite side to the active C85 site (Figure S6G(ii)). The C111 thiol is exposed on the protein surface and is solvent-accessible, yet the C107 thiol appears to be buried by ubiquitin (Figure S6G(iii)). However, a closer inspection of the E2D2-ubiquitin crystal structure (PDB: 3a33) (*44*), stabilized by mutating E2D2-C85S (enabling a stable peptide bridge with ubiquitin’s G76, Figure S6G(iv), left), revealed that in the absence of MDM2, the ubiquitin unit flipped to the other side of E2D2, partially exposing C107 (Figure S6G(iv), right; the C107 thiol remains partially buried). Therefore, docking of an E3 ligase should minimally affect solvent accessibility of the C107XXXC111 motif, especially given high flexibility of the ubiquitin orientation relative to E2D2. Considering both structural models, we focused our remaining experiments on C111.

### C111 of E2D is a conserved physiological Cu^+^ sensor critical for *Drosophila* morphogenesis

We initially tested if endogenous cellular Cu^+^ could be mobilized to promote E2D2 activity or whether enhanced activity occurred only during Cu^+^ excess. We prepared Hap1 E2D2-KO cell lysates and added back recombinant WT (E2D2^WT^) or C111S mutant (E2D2^C111S^) enzymes (Figure S7A). As expected, E2D2^WT^ added to these lysates induced a prominent polyubiquitination blush. This was dose-dependently attenuated by selective, strong Cu^+^ chelation (bathocuproine:tetrathiomolybdate, 1:1), demonstrating that endogenous Cu^+^ indeed amplified the polyubiquitination. In contrast, application of recombinant Cu^+^-insensitive E2D2^C111S^ only modestly enhanced polyubiquitination in the lysate, and attenuation of the blush upon Cu^+^ chelation was muted, indicating that E2D2’s Cu^+^-sensitivity is physiologically relevant.

Among human E2s, the Cu^+^-binding residues C107 and C111 constitute a unique signature of the E2D clade (Figure S7B). Despite evolutionary conservation (*30*), in most eukaryotic lineages polar hydrophilic amino acids reside in position 107 or 111 (or both, Table S7), indicating that the ancient E2D orthologue in the last eukaryotic common ancestor was not equipped with a high-affinity Cu^+^-binding domain (Figure S7C). Indeed, the Cu^+^-binding motif C107XXXC111 is unique to multicellular animals (Table S7). While horizontal gene transfer and convergent evolution could not be ruled out, appearance of the C107XXXC111 motif across major non-bilaterian and bilaterian lineages indicates that it was likely to exist at the root of the animal kingdom (Figure 7A). In distant unicellular holozoans (e.g., filastereans and ichthyosporeans) and in choanoflagellates, regarded as the closest single-celled relatives of metazoans (*45*), threonine occupies the 111 position, indicating that the Cu^+^-binding motif was unlikely to exist in the last holozoan common ancestor (Figure 7A). Thus, emergence of this motif could probably be dated to the Neoproterozoic, sometime between the last common unicellular and multicellular metazoan ancestors (Figure 7A).

**Figure 7.**
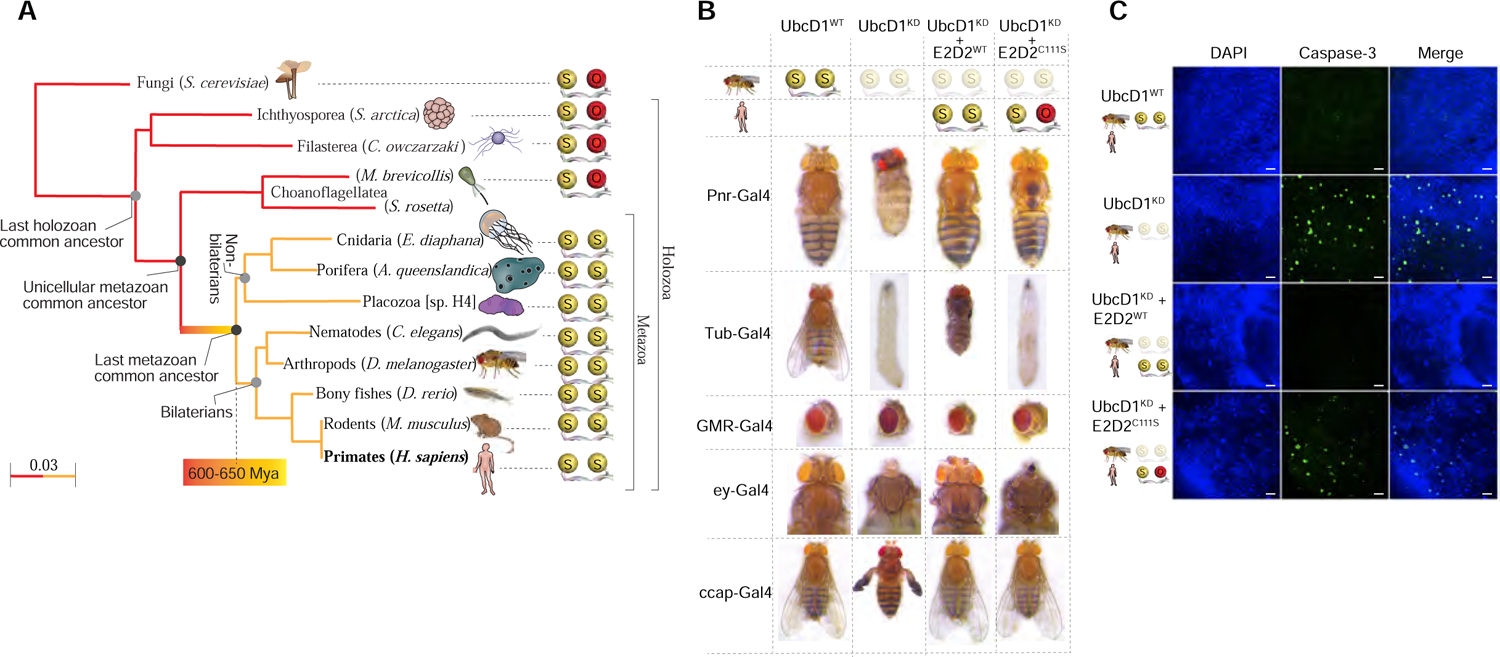
C111 of E2D is a conserved physiological Cu^+^ sensor critical for *Drosophila* **morphogenesis** (A) Evolutionary tree of E2D2 orthologues among holozoa. Representative species from major animal lineages and from three independent, closely related, unicellular holozoan lineages were used for generating a Fast Minimum Evolution tree based on Grishin protein distance (left-lower corner scale), with human E2D2 as reference. Side chain atoms of the Cu^+^-binding ligands at position 107 (left) and 111 (right) are shown as spheres, coloured yellow (sulfur at C107) and red/yellow (oxygen/sulfur at T/C111, respectively). Appearing only after separation of unicellular and multicellular metazoan lineages, yet evident across multiple animal lineages, emergence of a fully developed Cu^+^ ligand configuration could be approximated. (B) Phenotypes caused by targeted *UbcD1* knockdown. Control phenotypes (Gal4 transgene only) are shown on the left column. Second column shows the effect of *UbcD1* knockdown in the adult thorax (Pnr-Gal4; cleft and hyperpigmentation), the entire animal (Tub-Gal4; larval lethality), the eye (GMR-Gal4, rough eye), the developing head / eye (Ey-Gal4; complete loss of head structures) and crustacean cardioactive peptide (CCAP)-expressing neurons (CCAP-Gal4; failure of wing expansion). Third column shows near-complete rescue by co-expression of human E2D2-WT. Fourth column shows reduced / absent rescue by co-expression of human (Cu^+^-insensitive) E2D2-C111S. (C) Brain sections showing induction of active Caspase 3 in *Ey-Gal4* expressing cells upon knockdown of *UbcD1* and following expression of human E2D2-WT or E2D2-C111S. Sections were stained with DAPI (blue, nuclear staining) and Caspase-3 activity was determined (green signal). Scale bars represent 10 µm. See also Figure S7 and Table S7.

Novel signaling pathways that appeared at the metazoan root often modulate cell fate and morphogenesis by co-opting pre-existing cell proliferation mechanisms (*46*), and given C107XXXC111 motif conservation throughout bilaterian evolution (Figure 7A), we explored its developmental relevance in the fruit fly. Short of the four-paralogue repertoire that evolved in vertebrates, the *D. melanogaster* orthologue UbcD1 (aka Effete) shows a remarkable homology with human E2Ds, especially with E2D2 (Figure S7D and Table S7). Using the GAL4/UAS bipartite *in vivo* gene manipulation system with three independent UAS-RNAi lines (Figure 7B), we demonstrated that *UbcD1* knockdown: 1) When ubiquitous, caused early larval death, indicating that *UbcD1* is an essential gene; 2) during early head / eye development resulted in complete loss of the head structures; 3) in the central dorsal stripe resulted in a thoracic cleft plus increased pigmentation in affected cells; 4) in the entire central nervous system caused early pupal death; 5) in the CCAP neuropeptide-producing cells caused a failure of normal adult wing expansion; and 6) in the developing eye caused loss and disorganization of the normal ommatidial array. These phenotypes could be explained by UbcD1’s known roles in promoting apoptosis (*47, 48*), neuronal dendrite pruning (*49, 50*) and Hedgehog pathway signaling (*51*). Remarkably, similar phenotypes have also been associated with copper dysregulation (*52–55*). Moreover, one of UbcD1’s known E3 ubiquitin ligase partners, Slmb, has been identified as a novel regulator of copper homeostasis (*51, 56*).

To ascertain the importance of E2D2’s Cu^+^-sensing C111 residue *in vivo*, we generated transgenic *Drosophila* strains allowing the targeted expression of human E2D2, either WT (*h*E2D2^WT^) or C111S-mutant (*h*E2D2^C111S^). With all the GAL4 driver lines tested earlier, *h*E2D2^WT^ expression provided an almost complete rescue of the detrimental phenotypes induced by *UbcD1* knockdown (Figure 7B), reconfirming the expected functional conservation across species (Figure S7D). However, despite being transcribed at the same levels as *h*E2D2^WT^ (since transgene insertion occurred at the same genomic location), expression of *h*E2D2^C111S^ offered only minimal rescue (Figure 7B). The hypomorphs were characterized by caspase-3 activation in neuronal tissue (Figure 7C), indicating that cell death was mediated by apoptosis. Linking the allosteric Cu^+^-sensing C107XXXC111 motif of E2Ds with *Drosophila* development and head formation, these functional data are concordant with pathways that were over-represented in the mammalian cell-based Cu^+^-enhanced ubiquitome (Figure 3C and Table S3), highlighting that Cu^+^ signaling via C107XXXC111 is *functionally* (and not only *structurally*) conserved across metazoan evolution.

## DISCUSSION

Copper has only recently been recognized as a potential signaling ion in brain and other tissues (*2*). Direct partial *inhibition* of PDE3B activity by copper binding to C768, located away from the enzyme active site, emerged as the first example of a modulatory Cu^+^ binding site (*57*). This was reflected in 3T3-L1 cells, where extracellular copper supplementation (50 µM) potentiated lipolysis. Here we show what, to the best of our knowledge, is the first example of copper signaling inducing an allosteric conformational change that *enhances* enzyme activity. We have delineated an unexpected confluence between ubiquitin conjugation and copper, interwoven by a unique sub-fM-affinity Cu^+^-binding site (C107XXXC111) on the E2D clade. This tunable switch markedly promotes target polyubiquitination and can thus regulate the degradation rate of many proteins, contributing to development and, notably, head formation in *Drosophila*. While we demonstrated that E2D2 activation is a major component of the copper-induced polyubiquitination blush, and that activation of the E2D clade is likely to mediate much of the protein turnover induced by copper supplementation in cell culture, we have not excluded lesser contributions from other components of the ubiquitin-proteasomal system, including potential interplay between copper and E3 ligases (*58*) or deubiquitinating enzymes (*59*).

The high-affinity Cu^+^ allosteric binding site on E2D evolved (Figure 7A) contemporaneously with increasing copper bioavailability in Neoproterozoic oceans. Ancient geochemical shifts fashioned metal–protein partnerships (*60*) and indeed, the appearance of this site might reflect a concurrent transition to copper-based nitrogen assimilation (*61*). In this setting it may have been advantageous for a nascent metazoan kingdom to evolve a Cu^+^-signaling mechanism that couples protein turnover with surrounding nitrogen availability. Coinciding with evolution of other cuproenzymes (*62*), incorporation of the C107XXXC111 motif into E2s would link Cu^+^ levels with the turnover of p53, which had served a protective role in unicellular holozoans (*63*). With the advent of animal multicellularity, this signaling pathway could tune p53 to control cell fate and organism size (*46*), eventually evolving to influence tissue morphogenesis (Figure 7B-C).

The discovery of this novel p53-regulating signaling pathway has broad biomedical implications. p53 has been shown to interact with SCO2 (*64*), which carries the CXXXC motif and has regulatory roles in cellular copper homeostasis (*65*). In the opposite direction, and consistent with data showing that copper chelators increase p53 (*66*), we found that Cu^+^ promoted p53 degradation by allosterically activating E2D2. p53 depletion is a critical initiator of malignancy, and cancer tissue exhibits a mysterious high demand for copper (*1, 2, 67, 68*). Our findings could link increased cancer cell copper uptake with p53 depletion, accounting for the emerging benefits of copper modulation in cancer treatment (*69*). Conversely, major neurodegenerative disorders are complicated by low tissue copper concentrations (*70–72*) and ubiquitinopathy alongside elevated p53 (*73*). Based on our findings, decreasing total brain copper levels with advancing age (*74*) or neurodegeneration could elevate p53 and impair proteostasis. The cause of age- and disease-dependent brain copper supply failure is uncertain, but is unlikely to be corrected nutritionally (*75*), as Cu^+^ does not passively cross the blood-brain-barrier. The present data indicate that correction of defective proteostasis could underlie the enigmatic potential of copper chaperones to rescue neurodegeneration (*76*).

## ACKNOWLEDGEMENTS

The Florey Institute of Neuroscience and Mental Health acknowledges the strong support from the Victorian Government and in particular the funding from the Operational Infrastructure Support Grant. This work was supported by the Australian Research Council (AIB, ZX and CO), the National Health and Medical Research Council (AIB, DNS), and the CRC for Mental Health (AIB); grant ACT-04 from the Chilean Government (CO); as well as MNDRIA, Guest Family (DNS). We thank A. Ciechanover (Technion-Israel Institute of Technology, Israel), G. Schwarz (University of Cologne, Germany), B. J. Monahan and D. Komander (Walter and Eliza Hall Institute of Medical Research, Australia), J. Camakaris, B. Dean, G. Pavey, R. A. Cherny, S. Luza, I. Volitakis, V. Perreau, A. Southon, A. Lothian, B. Turner and K. Dent (The Florey Institute of Neuroscience and Mental Health, Australia), A.G. Wedd and Shenggen Yao (Bio21 Institute, The University of Melbourne, Australia), S. Lutsenko (Johns Hopkins University), Chris Chang (Berkeley University), L.G. Aguayo, C. López, L. Guzmán, G. Moraga-Cid, M. Reyes (University of Concepción, Chile), C. Peters (Max Planck Institute of Neurobiology, Germany), C. Aylwin, (Oregon Health and Sciences University, USA), M. A. Cater (Deakin University, Australia), M. Avila (Universidad de las Américas, Chile), G. De Ferrari (Universidad Andrés Bello, Chile), B. Roberts (Emory University, USA), N. Faux (IBM Research-Australia), M. Cortés and E.L. Robb (Mental Health Research Institute of Victoria, Australia). We are grateful for Israel Science Fund grant 2343/18 (AL) and the AUSiMED Fellowship Grant made possible by the generosity of the Lowy Foundation (AL).

## AUTHOR CONTRIBUTIONS

H.T. M.A.G. and C.M.O, performed the Western blot analyses. A.A.U. and Z.X. prepared the plasmids, expressed and purified the E2D2 proteins, performed the mutation analysis on E2D2 and the copper binding studies. J.M. collected MS ubiquitomics data. D.N.S. and A.L. analysed the ubiquitome data. M.A.G. performed the AMS binding assay, proteasome activity assay and co-immunoprecipitation experiments. C.M.O, C.M.L. and H.T. performed the *in vitro* ubiquitin conjugation assays. C.M.O., C.M.L. and C.H.M. performed the protein pulse-chase experiments. Z.X. designed the structural analyses. A.R. and C.M.O performed the studies with siRNAs. A.L. performed the sequence, evolution and statistical analyses. R.B., C.M.O., Z.X. and A.I.B. discussed the *Drosophila* studies. R.B. designed, evaluated and interpreted the *Drosophila* data. B.Z. performed the *Drosophila* studies. P.G. and Z.X. designed and performed the NMR studies and P.R.G., Z.X and A.I.B. discussed the NMR data; A.L., C.M.O, Z.X., R.B., D.N.S, P.R.G and A.I.B. wrote the manuscript. All authors read and approved the final manuscript.

## DECLARATION OF INTERESTS

Prof. Bush is a shareholder in Alterity Therapeutics Ltd, Cogstate Ltd, Brighton Biotech LLC, Grunbiotics Pty Ltd, Eucalyptus Pty Ltd, and Mesoblast Ltd. He is a paid consultant for, and has a profit share interest in, Collaborative Medicinal Development Pty Ltd.

## METHODS

### Primary cultures and cell lines

#### Mouse cortical primary cultures

Cortical neuronal cultures were prepared as described previously (*77*) in accordance with ethics committee approval of the University of Melbourne. Briefly, cortices were removed, dissected free of meninges, and dissociated in 0.025% (w/v) trypsin. Dissociated cells were plated in poly-l-lysine coated sterile 6 well culture plates in minimal essential medium (MEM) supplemented with 10% fetal calf serum. Cultures were maintained at 37°C in 5% CO_2_ for 2 h before the plating medium was replaced with Neurobasal growth medium containing B27 supplements (Life Technologies Inc., USA). The cells were used for experiments after 6 days in culture. The medium was replaced with Locke’s media (in mM: NaCl, 154; KCl, 5.6; CaCl_2_, 2.3; MgCl_2_, 1; NaHCO_3_, 3.6; glucose, 5; Hepes, 5; pH 7.4) (*77*), into which experimental compounds were added.

#### Fibroblasts

The wild-type mouse embryonic fibroblasts were a generous gift from Bart De Strooper and have been previously described (*78*). The SV-40 transformed normal human fibroblast cell line GM2069 (Coriell Institute, Camden, NJ) has been previously described (*79*). **Parental and E2D2 knockout cells lines**. HAP1 parental and E2D2 knockout cell lines were obtained from a commercial source (Horizon, UK). HAP1 cells are derived male chronic myelogenous Leukemia (CML) cell line KBM-7 (*80*). E2D2 was knocked out with CRISPR Cas9 gene editing technology. Both cells were cultured in Iscove’s Modified Dulbecco’s Medium (IMDM) with 10% FCS and 1% Pen/Strep.

#### Cell lines

Chinese hamster ovary (CHO) cells, Human Embryonic Kidney (HEK293T) cells, HeLa, HepG2, NSC34 motoneuron-like cells, N2a mouse neuroblastoma cells, mouse embryonic fibroblast and human fibroblasts were maintained in Eagle’s Dulbecco’s Modified Eagle’s Medium (DMEM; Life Technologies Inc., USA) supplemented with 10% (w/v) fetal calf serum (Bovogen, France), penicillin and streptomycin. N27 cells were maintained in RPMI (Life Technologies Inc., USA) supplemented with 10% (w/v) fetal calf serum (Bovogen, France). Parental HAP1 and E2D2 KO cells (Horizon Discovery, UK) were maintained in Iscove’s Modified Dulbecco’s Medium (IMDM, Life Technologies Inc., USA) supplemented with 10% fetal calf serum (Bovogen, France). Cultures were incubated at 37°C in 5% CO_2_. During experiments, cells were treated in Locke’s media (154 mM NaCl, 5.6 mM KCl, 3.6 mM NaHCO_3_, 2.3 mM CaCl_2_, 5.6 mM D-glucose, 5 mM HEPES, pH 7.4) (*77*) with various concentrations of CuCl_2_, NH_4_Fe(SO_4_)_2_, ZnCl_2_, Diamsar or MG132 (Calbiochem). Human Embryonic Kidney 293T (HEK) cells stably transfected with the human CTR1 construct within the pcDNA3.1 vector and the vector-only control HEK cells were maintained in DMEM low glucose with 2mM Glutamine, 10% (w/v) fetal calf serum (Life Technologies Inc., USA) and Geneticin (400µg/ml) (Life Technologies Inc., USA). E2D2 knockout cell lines were obtained from a commercial source (Horizon, UK). E2D2 knockout cells are derived male chronic myelogenous Leukemia (CML) cell line KBM-7 (*80*). E2D2 was knocked out with CRISPR Cas9 gene editing technology. Cells were cultured in Iscove’s Modified Dulbecco’s Medium (IMDM) with 10% FCS and 1% Pen/Strep.

#### Western blot

Control and copper-supplemented cells lines were washed once with PBS and harvested in 50 mM Tris HCl pH 7.5 containing 1 % (v/v) NP-40, NaCl 150 mM, 10 mM N-ethylmaleimide (NEM), protease inhibitors cocktail without EDTA (Roche, USA) and phosphatase inhibitors cocktail (Roche, USA), using cell scrapers. The homogenized samples were submitted to 10 cycles of 10 s at maximal intensity at 4⁰C in a sonicator bath (Bioruptor, Diagenode, USA). Then the homogenates were centrifuged at 10,000 g for 10 min at 4⁰C and the supernatants (NP40-soluble fraction) were collected, and protein concentrations were determined using the BCA Protein Assay Kit (Pierce, USA) and supernatants were aliquoted for Inductively coupled plasma mass spectrometry (ICP-MS) and Western blot analysis. These samples were stored at −80⁰C until use. 10µg of protein was submitted to SDS-PAGE electrophoresis using NuPAGE Novex or Bolt 4-10% Bis-Tris gels (Thermo Fisher Scientific, USA). Proteins were transferred to nitrocellulose membranes using the iBlot Dry Blotting System (Thermo Fisher Scientific, USA) and ubiquitinated proteins were detected with a mouse monoclonal antibody (diluted 1:500; P4D1; Santa Cruz Biotechnology, USA), a rabbit polyclonal antibody (diluted 1:500; Z0458; DAKO, Denmark), a mouse monoclonal antibody (diluted 1:500; FK1, Enzo Life Sciences, USA) or a mouse monoclonal antibody (diluted 1:500; FK2, Enzo Life Sciences, USA). Mouse monoclonal or rabbit polyclonal anti-actin and anti-GAPDH antibodies (diluted 1:10,000; Sigma-Aldrich, USA) were used to detect ß-actin and GAPDH, respectively, which were used as loading controls. The signal was detected by using secondary antibodies coupled to infrared fluorescent dyes (diluted 1:10,000; IR680 and IR800, Li-Cor, Lincoln, USA) or by using secondary antibodies coupled to horseradish peroxidase (HRP) (diluted 1:10,000; DAKO, Denmark). The infrared integrated intensity was acquired in a LiCor imager (Li-Cor, Lincoln, USA) using the Odyssey 4.0 analytical software and the HRP activity was visualized after incubation in ECL Western Blotting Detection Reagent (GE Healthcare Life Sciences) and visualized using a Fuji film imaging system. Densitometric evaluation was performed using Multi Gauge Version 3.0 (FujiFilm).

#### ICP-MS analysis

50 µL of NP-40 soluble fraction was lyophilized for 12 h and then digested with nitric acid under heating conditions. After that, hydrogen peroxide was added to each sample and they were heated again. Measurements were made using an Agilent 7700 series ICP-MS instrument under routine multi-element operating conditions, using a Helium reaction gas cell. The samples were analysed without identification. The levels of copper were corrected by proteins (g). **MTT cell viability assay**. Control and copper-supplemented mouse cortical primary neurons (6 DIV) were washed once with PBS and then incubated with the dye 3, [4,5-dimethylthiazol-2-yl-] diphenyltetrazolium bromide (MTT, 0.5 mg/mL). After 1 h of incubation, the media was removed, and the cells solubilized with 100% DMSO. Cell viability was assessed using an EnSpire Multimode Plate Reader (PerkinElmer Inc., USA) measuring the formazan absorbance at 560 nm. **Trypan blue exclusion**. Control and copper-supplemented cells were washed once with PBS and trypsinized for 5 min with 1x sterile trypsin solution (Thermo Fisher Scientific, USA). Cell suspensions were mixed with equal volume of 0.4% Trypan blue solution (Thermo Fisher Scientific, USA). Ten μL of this suspension was loaded on a hemocytometer. Cell viability was quantified in a Countess automated cell counter (Thermo Fisher Scientific, USA).

#### DCF oxidation

Intracellular levels of reactive oxygen species (ROS) were measured using the cell-permeant redox-sensitive probe 2’,7’-dichlorodihydrofluorescein diacetate (H2DCFDA) (Molecular Probes, Thermo Fisher Scientific, USA). HEK 293 cells were grown in a 96-well plate for fluorescent-based assays until ∼80% confluence. Then, the cells were incubated with H2DCFDA (10 µM) in Locke’s media for 60 min at 37 °C, followed by a washing step and a subsequent treatment with H_2_O_2_ (up to 250 µM) or CuCl_2_ (up to 10 µM) in Locke’s media at 37°C. The fluorescent signal was recorded in an EnSpire Multimode Plate Reader (PerkinElmer Inc., USA) (Ex/Em: 495/525 nm) and the initial rate of DCF oxidation calculated in the linear dynamic range during the first 5 min of exposure to the treatments (*81*).

### Cellular Protein Degradation Assay

Experiments were performed in 12 well plates (primary cortical cells, HAP1 parental and E2D2 knockout cell lines at 70-80% confluence), which were washed three times with 1 mL of Locke’s media. Then, cells were incubated with 1 mL of Met/Cys free media supplemented with 10 % FCS (dialyzed vs PBS) at 37°C for 15 min. Media were replaced with fresh media supplemented with Easytag Express S35 Protein Labeling Mix (PerkinElmer, 100-200 µCi/mL) for 1 h. Cells were washed and incubated in Met/Cys free media supplemented with 10 % FCS (dialyzed) or Locke’s media (500 μL per well) supplemented with cycloheximide (1 μg/μL), -/+ Cu (10 or 25 µM as chloride or histidinate) or MG132 (50 µM) (total volume of 500 μL). Media and cells were precipitated by cold-TCA for 48 h at 4 °C. Then, samples were centrifuged at 10,000 g for 10 min at 4 °C. Soluble and pellet fractions were collected. Pellets were resuspended in 2% SDS solution and sonicated indirectly for 30 min in a bath sonicator. Radioactive counts from the media that remain soluble after TCA precipitation (peptide fragments) and pellets from the cells (total non-degraded cell proteins) were measured in a TopCount radioactivity detector (Perkin Elmer). A “protein degradation index” was calculated as the ratio between the counts detected in the soluble media and the counts detected in the pellet cell fraction (*82*).

### Autoradiography

^35^S-positive bands within NP40 cell extracts were detected in a Typhoon system (GE Healthcare Life Sciences, UK). The autoradiography protein profile observed in the non-supplemented or Cu-supplemented cells was represented as a heatmap of blot intensity (ranging white to blue) for 90 to 120 equal bins averaged across each experimental condition.

### Cellular proteasome activity

Control and copper-supplemented cell lines were washed once with PBS and harvested in 10 mM HEPES (pH= 7.6), using cell scrapers. Then, the cells were homogenized with 20 passages through a 25 G needle and centrifuged at 10,000 g for 10 min at 4⁰C. The supernatants were separated, and the protein concentration determined as described above. Optimal concentrations of the samples were determined for each enzymatic assay, and the samples adjusted with 10 mM HEPES (pH = 7.6) to final working concentrations: chymotrypsin-like activity 0.2 µg/µl, trypsin-like activity 2 µg/µl, and caspase-like activity 0.4 µg/µl. Proteasome activities (chymotrypsin-like, trypsin-like and caspase-like) were assayed using the Proteasome-Glo kit (Promega, USA) in 384-well plates for luminescent assays. Twenty-five µl of each specific protease reagent was added to 25 µl of each sample. A blank (10 mM HEPES; pH=7.6) as well as a standard curve of purified 26S Proteasome (0.5, 1 and 10 g/L; Enzo Life Sciences Inc., USA) were included in each assay. Each enzymatic activity was measured after 20 minutes of incubation when the signal was in the dynamic range (Promega, USA): the luminescent signal of each well was detected and quantified using a FlexStation 3 reader (Molecular Devices, USA). Data are expressed as arbitrary units related to 26S standard curve activity. A combined proteasome activity measure was calculated using a mixed model reflecting a mean of the three enzyme-like activities within each sample.

### siRNA transfections

HEK293, HeLa and HepG2 cells were seeded at 3×10^5^ cells per well in 6-well plates and incubated at 37 °C and 5% CO_2_. Following 24 h of culture, cells were transfected with siRNA. Cells were treated either with 15 nM AllStars negative control siRNA (Qiagen, USA), or a combination of E2D-specific siRNA (5 nM of each siRNA): E2D1 (5’-UCCACUGGCAAGCCACUAUUA-3’), E2D2/3 (5’-AACAGUAAUGGCAGCAUUUGU-3’) and E2D4 siRNAs (5’ – CCGAAUGACAGUCCUUACCAA-3’) all from Qiagen, USA(*83*).

siRNA complexes were formed using HiPerfect transfection reagent (Qiagen, USA). The volume of siRNA for a final concentration of 15 nM per well was mixed with 6 μL of HiPerfect in 100 μL of Dulbecco’s modified Eagle’s medium (DMEM, Gibco, USA), incubated for 10 min and added to cells in media without antibiotics, as recommended by the manufacturer. Following 24 h culture media was changed, and a second siRNA transfection was conducted. After 24 h cells were supplemented with CuCl_2_ (2.5 and 5 μM) for 3 h in Locke’s buffer. Cell viability was monitored by PrestoBlue assay (Thermo Fisher Scientific, USA). After treatment cells were prepared for Western blot and ICP-MS analysis.

### PrestoBlue viability assay

To evaluate the effect of siRNA transfections -/+ copper treatments on cell viability, cells were washed and incubated with PrestoBlue viability reagent diluted in Locke’s buffer (1:10) for 30 min after the treatment. The fluorescent signal was determined using the Flexstation® 3 Plate Reader (Molecular Devices) for fluorescence reading (Excitation λ: 560 nm and Emission λ: 590 nm).

### Immunoprecipitation

HEK293 cells were grown to 80-90% confluence in 175 cm^2^ flasks before incubating in DMEM +/- 10 M CuCl_2_ for 2 h at 37 °C. Cell monolayers were washed three times in ice-cold Phosphate Buffered Saline (PBS), scraped into 10 mL PBS and centrifuged at 800 x g, 4 °C for 10 min. Cell pellets were lysed by resuspension and rotation at 4 °C for 30 minutes in immunoprecipitation lysis buffer containing 1% CHAPSO, 150 mM NaCl, 2 mM DTT, 1x EDTA-free protease inhibitor cocktail (Sigma, Australia), 50 mM Tris-HCl, pH7.4. Cell suspensions were centrifuged at 14,000 x g, 4 °C for 10 min. Supernatants were transferred to fresh tubes and pre-cleared using washed protein-G-Sepharose beads. Protein concentration was measured using a micro BCA protein assay kit and samples were diluted to 1 mg/mL total protein. Immunoprecipitation was performed by incubating samples in 2 g/mL anti-rabbit (FL-393, sc-6243) or anti-mouse (DO-1, sc-126) P53 antibodies (Santa Cruz Biotechnology) for 2 h at 4 °C, gently rotating. For immunocapture 50 µL of washed Protein-G-Sepharose beads were added to samples and incubated overnight at 4 °C, gently rotating. Brief centrifugation (30 sec, 14,000 x g, 4 °C) was used to collect beads prior to washing three times for five minutes each wash in PBS on a rotating wheel at 4 °C. To elute proteins, the washed beads were incubated in 50 μL of 2 x NuPAGE loading buffer supplemented with 3 % SDS and 20 mM DTT. Samples were vortexed to mix and incubated at 90 °C for 5 min prior to centrifugation for 30 s at 14,000 x g to pellet beads. The eluates were removed with crimped gel-loading tips to avoid aspirating beads. Samples were then separated by SDS-PAGE (as previously described) and anti-rabbit polyubiquitin antibodies (DAKO) were used to detect immunoprecipitated proteins. All reagents were purchased from Thermo, Australia, unless otherwise stated.

### p53 degradation

HEK293 cells were seeded at 300,000 cells/ mL and grown in full growth media (DMEM +10% FCS + Sodium Pyruvate) for 3 days prior to treatment. Cells were incubated in Locke’s media supplemented with cycloheximide (10 µM) -/+ CuCl_2_ (5 μM) for 3h at 37°C. The cells were harvested in cold PBS and pelleted at 4°C at 2,000 g for 5 min. The pellets were then lysed in a cytoplasmic extraction (CE) buffer (10 mM HEPES, pH 7.9, 10 mM KCl, 0.1 mM EDTA, 0.3 % NP-40 and 1 x protease and phosphatase inhibitors, Roche, USA) by passing through a 23G needle and syringe 30 times and then left to stand on ice for 5 min. After centrifuging for 5 min at 4, 000 g at 4°C, the supernatant was saved, and the remaining pellet was washed with CE buffer without NP40. The washed pellet was further lysed with Nuclear Extraction (NE) buffer (20 mM HEPES pH 7.9, 0.4 M NaCl, 1 mM EDTA, 25% Glycerol and 1x protease and phosphatase inhibitors) for 10 min on ice. The supernatant was collected after centrifugation at 4°C at 10,000 g for 5min. The collected fractions were measured for their protein content using the Pierce BCA Protein kit and 20 μg of protein from each sample were run on a NuPage 4-12% Bis-Tris Protein gel in 1x Bolt MES running buffer. The protein gel was then transferred to a nitrocellulose membrane using the iBlot2 and iBlot2 transfer stacks using the following parameters – 15 V for 5mins, 20 V for 6 min. and 25V for 2 min. The membrane was blocked with 5 % skim milk + 1 % BSA in TBS-T and incubated with primary antibody diluted in TBS-T. Fluorescent secondary antibody was used at a 1:10,000 dilution in TBS-T and the membrane was imaged using the Odyssey classic imager and Image studio for processing.

### Analysis of p53 redox state in cell culture

The oxidation state of p53 in control and copper-supplemented cells was assessed via alkylation of reduced cysteine residues with 4-acetamido-4’-maleimidylstilbene-2,2’-disulfonic acid (AMS) as we previously described (*84*). Total cell extracts were prepared and pre-incubated on ice for 1 h with 10% trichloroacetic acid to protonate all thiols and precipitate total cellular protein. The precipitates were centrifuged at 16,000 g for 15 min (4°C) and the pellets were dissolved in 0.1% SDS/ 0.67 M Tris-HCl pH 8 +/- 15 mM 4-acetamido-4’-maleimidylstilbene-2,2’-disulfonic acid (Thermo Fisher Scientific, USA). The samples were incubated at 37 °C for 2 h, fractionated on 4-12% NuPage Bis-Tris gels (Thermo Fisher Scientific, USA), transferred to nitrocellulose membranes and probed with an anti-p53 polyclonal antibody (Santa Cruz Biotechnologies, USA).

### Ubiquitomics (see also Figure S3)

#### Sample preparation

Cell lysates were harvested following incubation with CuCl_2_ (0, 10, or 25 µM) in ice-cold lysis buffer (50 mM Tris-HCl (pH 7.5), 150 mM NaCl, 0.5% NP-40, 1 mM DTT, protease inhibitors (EDTA-free protease inhibitor, Roche, USA), DUB inhibitor (10 mM N-ethylmaleimide) and phosphatase inhibitor (PhosphoStop, Roche, USA) using a POLYTRON hand homogenizer(*19*). Lysed cells were incubated for 10 mins on ice and then centrifuged at full speed for 10 min at 4 °C. The cleared supernatant was transferred to a fresh microcentrifuge tubes, discarding the cellular debris pellet. Total protein concentration was determined by BCA Protein Assay Kit (Pierce, USA), according to the manufacturer’s instructions. Lysates were kept at −80°C for long-term storage.

#### Ubiquitin Affinity Purification

Ubiquitin-associated proteins were affinity purified from cell lysates using specialized affinity matrix (VIVAbind Ubiquitin Matrix, VB2950, VIVA Bioscience), according to the manufacturer’s protocol with additional bead washing. Briefly, 500 μg of total lysate was incubated with 30 μL VIVAbind Ubiquitin Matrix agarose bead slurry for 2 hours at 4 °C with end-to-end rotation. Beads were washed six times (50 mM Tris-HCl (pH 7.5), 150 mM NaCl, 1 mM dithiothreitol) to remove detergent residue from contaminating downstream mass spectrometry analysis.

### Trypsin digest and peptide desalting

Proteins were digested in Buffer 1 (2 M urea, 50 mM Tris-HCl (pH 7.5), trypsin (5 μg/mL)) for 30 mins at 27 °C with vigorous shaking. Digested beads were briefly centrifuged, and the supernatant collected into fresh LoBind microcentrifuge tubes (Eppendorf). Beads were washed twice in Buffer 2 (2 M urea, 50 mM Tris-HCl (pH 7.5), 1 mM dithiothreitol) and the supernatants pooled with the trypsin digest and left to digest overnight at room temperature. The following day, the digested peptides were alkylated with 20 μL iodoacetamide (5 mg/mL) (Life Science, Sigma-Aldrich, GmbH) and incubated for 30 min in the dark. Alkylated peptides were acidified with 1 μL of trifluoroacetic acid (TFA) (Fluka BioChemika, Sigma-Aldrich, GmbH) to inhibit trypsin and acidify peptides for desalting using C-18 StageTips. The C18 StageTips were constructed in-house using 6 layers 3M Empore solid phase extraction C18 (Octadecyl) membrane (Agilent Technologies) inserted into a 200 μL LoBind micropipette tip. StageTips were activated with methanol (Vetec Fine Chemicals), equilibrated with 0.1% (v/v) TFA, 60% acetonitrile (MeCN) (LC/MS grade, Fisher Scientific), followed by two washes with 0.1% (v/v) TFA. Peptides were eluted using a two-step elution with 0.1% (v/v) TFA, 60% (v/v) MeCN and then dried using a vacuum concentrator. Dried peptides were stored at – 80 °C.

### NanoLC-MS/MS (mass spectroscopy)

Dried peptides were resuspended in 5% (v/v) formic acid, 2% (v/v) MeCN and loaded onto a 20-cm column with a 75 μm inner diameter, packed in-house with 1.9 μm C18 ReproSil particles (Dr. Maisch GmbH). Peptides were then introduced by nanoelectrospray into a LTQ Orbitrap Velos Pro coupled to an Easy-nLC HPLC (Thermo Fisher) over a linear gradient of acetonitrile at 250 nl/min over 140 min. Tandem mass spectrometry data were collected for the top 10 most abundant ions per scan. The mass/charge (*m/z*) ratios selected for MS/MS were dynamically excluded for 30 s.

### Raw data processing of MS data

Raw data files were processed using the MaxQuant computational platform for mass spectrometry-based shotgun proteomics v1.2.7.4 (*85*). Database searching was performed against the UniProt Mouse Database (2015) using the Andromeda search engine integrated into MaxQuant. A false discovery rate of 1% was tolerated for protein, peptide, and sites, and one missed cleavage was allowed. MaxQuant parameters used in the analysis are provided in Table S1.

### Filtering of MS data

MaxQuant-processed data were filtered using the statistical workflow *Pegasus* - a modified workflow within the R software environment based on Perseus - to remove contaminants, reverse hits, proteins only identified by site, and proteins with < 2 unique peptides as previously described (*86*). All further analysis was performed using the filtered data.

### Annotation of MS data

Due to the great structural and sequence similarity among mammalian orthologous proteins, it is quite common to perform interaction experiments across species, and such cross-species experiments represent a large fraction of the available evidence in literature (*87*). As our simulations suggested that the STRING database contains more information on the human protein-protein interaction network, we used human orthologues for protein-protein interaction analysis. We uploaded the column corresponding to the heading ‘Gene’ from UniProt Mouse Database into STRING’s search engine and retrieved a list of corresponding Ensembl protein IDs (ENSP) in humans, on which STRING database relies, together with their amino acid sequences. This list was then cross-matched back to the original identifiers, and known E2 enzymes were removed, resulting in the final list of Ub-associated proteins that were explored in all subsequent analyses.

### Processing label-free quantification (LFQ) data

We used Perseus (v. 1.6), a software package for shotgun proteomics which performs bioinformatic analyses of the output of MaxQuant (*85*). LFQ data was log-transformed to achieve normality. Before proteomics data can be used for the actual data analysis, they need to be subjected to missing-value imputation. A frequently used assumption in proteomics experiments is that low expression proteins give rise to missing values, therefore a Gaussian distribution with a median shifted from the measured data distribution median towards low expression should result in accurate imputation of such values. Default imputation settings (http://www.coxdocs.org/doku.php?id=perseus:user:use_cases:interactions), shrink distributions to a factor of “0.3” and shift it down by “1.8” standard deviations) were applied. Given the anticipated effects of copper on the abundance of Ub-associated proteins, replacement of missing values was applied to each expression column separately. For quality control, we generated histograms to verify that the distributions of the imputed values did not form a separate normal distribution, that they started around the same point in the replicates and were narrower than the distributions of the measured values. We also inspected multi-scatter plots to assure that while correlations between each possible pair of columns were high, correlation within the groups was slightly higher than between the groups of replicates. Following Z-score transformation across rows, heatmaps were used to visualise Euclidean distance-based hierarchical clustering according to media conditions and protein abundance profile.

### Ontology enrichment

Analysis was performed using Metascape (metascape.org), an online bioinformatics pipeline for multiple gene lists, which supports effective data-driven gene prioritization decisions (*88*). The analysis workflow included, (i) Based on two modified (S_0_=2) independent samples *t*-tests (permutation-based false-discovery rate with a cut-off q-value of 0.05), ubiquitin-associated proteins that displayed copper-mediated differential abundance were categorized into three distinct lists: a) those whose abundance was higher in copper-supplemented (CuCl_2_ 10 µM) vs. control conditions; b) those whose abundance was higher in copper-supplemented (CuCl_2_ 25 µM) vs. control conditions; and c) those whose abundance was higher in control vs. copper-supplemented (CuCl_2_ at either 10 µM or 25 µM) conditions; (ii) Extraction of annotations for the gene list using GO Biological Processes, Reactome Gene Sets, CORUM and KEGG Pathways (default options); and (iii) Functional enrichment analysis of the gene list. Minimal overlap of three genes, enrichment factor of 1.5 and P-value of 0.01 were used for filtering. Remaining significant terms were then hierarchically clustered into a tree based on kappa-statistical similarities among their gene memberships. A kappa score of 0.3 was applied as the threshold to cast the tree into term clusters. A subset of up to 10 representative terms from each of two clusters enriched among proteins whose abundance was higher in copper-supplemented conditions were then selected and converted into a network layout. Networks were visualized with Cytoscape using ‘force-directed’ layout with edges bundled for clarity.

### Quantifying relative abundance

To determine which Ub-modified proteins display a relative abundance profile that is significantly altered by copper, a modified one-way ANOVA based on a robust method for detection of significantly changing biomolecules in large omics datasets was employed (*89*) (a constant of 2, as per recommended settings [http://www.coxdocs.org/doku.php?id=perseus:user:use_cases:interactions], was added to the within-group variance, followed by permutation-based false-discovery rate with a cut-off q-value of 0.01, as per recommended settings). Finally, Ub-associated proteins that displayed Cu-mediated differential abundance were categorized into those whose abundance increased or decreased in Cu-supplemented media based on the direction of the difference between sample means in the 6 Cu-supplemented vs. 3 control columns. The number of Cu-mediated under- and over-abundant Ub-associated proteins was contrasted using binomial distribution.

### Interactome of Cu^+^-enhanced Ub-associated proteins

Proteins that displayed overabundance in the ubiquitin-associated proteome (ubiquitome) from cells grown in Cu-supplemented media were used as input. Using the STRING 10.5 Database for protein-protein interactions (PPIs) (*90*), for defining an interaction as positive we selected the high confidence cut-off (combined score of 0.7 or above). STRING calculates the combined score by integration of several types of associations: experimentally-determined-interactions (such as co-purification and co-crystallization experiments), coexpression (proteins whose genes are observed to be correlated in expression, across many experiments), phylogeny, homology, neighborhood and database-annotated (using known metabolic pathways, protein complexes, signal transduction pathways, etc from curated databases). We excluded unsupervised textmining (automated text-mining of the scientific literature) from the combined score calculation to reduce bias that could potentially reflect disproportional citations of some proteins. Next, from within Cytoscape 3.6.0 framework (*90*) we used the NetworkAnalyzer (*91, 92*) to identify the major connected component within the network.

### Topological analysis

Focusing on the retrieved subnetwork, a topological network analysis was performed; for each node (protein) we quantified betweenness centrality (*91*), which reflects the amount of control that this node exerts over the interactions of other nodes in the network (*93*).

Betweenness centrality *C_b_(n)* of a node *n* is computed as follows^82^: *C_b_*(*n*) = ∑*_s≠n≠t_* (*σ_st_* (*n*) / *σ_st_*), where *s* and *t* are nodes in the network different from *n*, *σ_st_* denotes the number of shortest paths from *s* to *t*, and *σ_st_* (*n*) is the number of shortest paths from *s* to *t* that *n* lies on. The betweenness value for each node *n* is normalized by dividing by the number of node pairs excluding *n*: (*N*-1)(*N*-2)*/2*, where *N* is the total number of nodes in the connected component that *n* belongs to. Thus, the betweenness centrality of each node is a number between 0 and 1.

### Identifying Ub-associated proteins that interact with E2Ds or E2A/B

Using the same STRING Database (*90*), in this analysis we selected the medium confidence cut-off (combined score of 0.4 or above, excluding unsupervised textmining) for defining an interaction, as interactions were measured at a group level, without focusing on attributes of any particular substrate. With 882 potential substrates and either four E2D or two E2A/B enzymes as input, the network generated by STRING was loaded into Cytoscape. We identified which Ub-associated proteins interacted with at least one member of the E2D clade (E2D1/2/3/4) and/or with at least one member of the E2A/B clade.

### Enrichment of Cu-enhanced substrates within each clade

Employing the hypergeometric test often used enrichment analysis (represented by a Fisher’s exact test), the proportions of Ub-associated proteins displaying Cu-enhanced overabundance among proteins interacting with a E2D enzyme and proteins interacting with a E2A/B enzyme were contrasted with the corresponding proportions calculated for proteins that did not display Cu-enhancement.

### Assessing the effect of specific protein characteristics

Content (fraction out of total amino acids) of the redox-sensitive amino acids cysteine and methionine was extracted from each protein’s amino acid sequence. Mean log-transformed values of protein mass and cysteine or methionine content for proteins that did versus those that did not display overabundance in cells supplemented with copper were compared with a non-paired *t* test.

### Controlling for covariates

To assess the effect of interacting with a specific E2 clade while controlling for mass, cysteine/methionine content and (co)interaction with the other clade, a multiple logistic regression model was employed in SPSS, with log_2_-mass, log_2_-cysteine/methionine content and being a E2D or E2A/B substrate as independent predictors and enhancement in Cu-supplemented media as the dichotomous outcome. Individual coefficients were examined in terms of their effect (as represented by Wald-statistic) and its confidence intervals.

### *In-vitro* ubiquitin conjugation

#### *In vitro* ubiquitin conjugation with Fraction II

*In vitro* polyubiquitination was measured using the protocol and components from the Ubiquitylation Kit (Biomol; Enzo Life Sciences). Some modifications were included in this *in vitro* system as described by Sato *et al*., 2008 (*94*). Biotinylated ubiquitin (2.5 µM) was incubated in the presence or absence of Fraction II, human recombinant UBA1 (100 nM), various E2 enzymes (∼2.5 µM), 2 mM ATP, 5 mM MgCl_2_, 50 µM CuCl_2_, 50 µM NH_4_Fe(SO_4_)_2_, 50 µM ZnCl_2_ and 5 mM DTPA in Tris-HCl 100 mM (pH 7.5) at 37°C for 5 h. For the experiments using Fraction II as the source of E-enzymes, Fraction II (the protein fraction of HeLa S3 lysate that binds to anion exchange resin, BostonBiochem) was firstly incubated in 5mM EDTA for 30 min and desalted using Zeba Desalt Spin Columns (Thermo Scientific, USA). Other components included: 1 mM DTT, 10 mM creatine phosphate (Sigma), 0.6 U creatine phosphokinase (Sigma), and 0.6 U inorganic pyrophosphatase (New England Biolabs). Samples were submitted to electrophoresis under non-reducing conditions. Proteins were transferred to nitrocellulose membranes using the iBlot Dry Blotting System (Life Technologies Inc., USA) and the formation of ubiquitin conjugates was detected by binding of avidin-HRP (Zymed, Life Technologies Inc.) using ECL detection system (GE Healthcare Life Sciences). The signal for polyubiquitinated proteins was recorded by using MultiGauge V3.0 (Fujifilm).

### Construction of expression plasmids

The DNA sequence encoding the full-length E2D2 protein was amplified by PCR and cloned into the expression vector pET20b between the restriction sites NdeI and BamHI to obtain the expression plasmid for the native protein. Site-directed mutagenesis was performed by PCR using the WT E2D2 gene as a template to construct the equivalent expression plasmids for protein variants E2D2-C21S, -M38L, -H55A, -C85S, -C_107S_, -C111S, - C21/85S and C21/107/111S. DNA sequencing confirmed the correct gene insert for each expression plasmid constructed.

### Protein expression, purification, and quantification

The expression plasmids were transformed into *E. coli* BL21(DE3) CodonPlus cells (Novagen) for protein expression. Each one liter of 2YT medium containing ampicillin (100 mg/ L) and chloramphenicol (34 mg/ L) was inoculated with an overnight culture (5 mL) of the expression cells. After the cells were grown aerobically with vigorous shaking at 37 °C to an OD_600_ ∼1.0, expression of the targeted protein was induced overnight at room temperature with IPTG at a final concentration of 0.5 mM. The cells were harvested by centrifugation and clarified cell extracts prepared in 20 mM Tris-Cl buffer, pH 8.0, 1.0 mM EDTA. The clarified cell extract was adjusted to pH ∼7.0 with acetic acid and the targeted protein was purified by a NaCl gradient elution (0 - 400 mM) from a SP Sepharose fast flow cation-exchange column equilibrated in 20 mM MOPS buffer, pH 7.0. The eluted fractions containing each targeted protein were pooled and concentrated. The protein was fully reduced by incubation with 5 mM DTT and further purified with a Superdex-75 FPLC gel filtration column in 100 mM KPi buffer, pH 7.4, 150 mM NaCl. The purity and identity of each purified protein were confirmed by SDS-PAGE and by electrospray ionization mass spectrometry (ESI-MS). For characterization and functional assay, each purified protein was fully reduced again by an overnight incubation with DTT (5.0 mM) inside an anaerobic glove-box and the excess DTT was completely removed via a Bio-Gel P-6 DG gel-desalting column (Bio-Rad) in thoroughly de-oxygenated MOPS buffer (20 mM, pH 7.3, 150 mM NaCl). The protein concentrations were estimated from the protein primary sequences (ε= 25,440 M^-1^ cm^-1^ at 280 nm for the fully reduced forms) and the reduced Cys thiol contents assayed in buffer containing 4 M urea with the Ellman’s reagent 5,5’-dithiobis-(2-nitrobenzoic acid). Both values matched each other for all final purified proteins, confirming the fully reduced states of all Cys thiols. Selected protein samples enriched with ^13^C and ^15^N isotopes were produced similarly from cells grown in a M9 minimal medium supplemented with ^13^C-glucose and/or ^15^N-NH4Cl for study by NMR spectroscopy.

### *In vitro* ubiquitination conjugation assays

i) Wild type (WT) or C21S, M38L, H55A, C85S, C_107S_ and C111S E2D2 mutants (5 µM) were incubated minus/plus CuCl_2_ (5 µM) in the presence of GSH (200 µM), UBA1 (100 nM), MgCl_2_ (5 mM), ATP (5 mM) and biotinylated-Ub (2.5 µM) in Tris-HCl (50 mM) for 1 h at 37 °C. The reaction was stopped by the addition of TCEP (50 mM). Conjugation activity of each these enzymes was detected by the formation of ubiquitin adducts bound to Streptoavidin-680IR. The infrared integrated intensity was acquired in a LiCor imager (Li-Cor, Lincoln, USA) using the Odyssey 4.0 analytical software. ii) Wild type (WT) and C111S E2D2 mutants (1 µM) were incubated in the presence of cell homogenates from E2D2 KO cells (10 µg) and GSH (200 μM), minus/plus bathocuproine:tetrathiomolybdate (up to 10 µM) for 1 h at 37 °C. iii) HeLa supernatants were incubated ± CuCl_2_ (10 or 50 µM) in the presence of GSH (200 µM) in Tris-HCl (50 mM) for 3 h at 37 °C. The ubiquitination activity was detected by using a mouse monoclonal anti-ubiquitin antibody (diluted 1:500; P4D1, Santa Cruz Biotechnologies, USA) and anti-mouse 800IR antibody. The infrared integrated intensity was acquired in a LiCor imager (Li-Cor, Lincoln, USA) using the Odyssey 4.0 analytical software.

### Analytical chemistry - quantification of Cu^+^ binding

The experiments were conducted via ligand competition for Cu^+^ according to reported protocols (*37, 95*). Briefly, chromophoric Cu^+^ complexes Cu^I^L2 (L = Fz, Bcs) were employed as the detection probes. Differences in their formation constants (log β_2_ = 15.1 and 19.8 for L = Fz and Bcs, respectively) (*37, 95*) mean that they buffer free Cu^+^ at different concentration ranges and thus may probe Cu^+^-binding sites with different affinities. Transfer of Cu^+^ from these probes to a protein target can be monitored from the change in solution absorbance characteristic of these complexes. The reported extinction coefficients were used for the concentration estimation (ε= 4,320 cm^-1^ M^-1^ at 470 nm for Cu^I^(Fz)_2_ and 13,000 cm^-1^ M^-1^ at 483 nm for Cu^I^(Bcs)_2_) (*37, 95*).

The complex Cu^I^(Fz)_2_ buffers free Cu^+^ at pM concentration range and was used to estimate the Cu^+^-binding stoichiometries for those protein sites that are able to remove Cu^+^ from the probe non-competitively. Titration of WT E2D2 protein into a probe solution of Cu^I^(Fz)_2_ (compositions in total concentrations: [Cu]_tot_ = 55 µM; [Fz]_tot_ = 300 µM; [NH_2_OH]_tot_ = 200 µM; [ascorbate]_tot_ = 100 µM) in 50 mM MOPS buffer, pH 7.4 led to a rapid transfer of Cu^+^ from Cu^I^(Fz)2 to the protein with precipitation. Samples were centrifuged for 10 min to completely remove the protein precipitates. The absorbance of the clarified solutions was found to decrease linearly with the amount of the added protein, suggesting that the protein removed Cu^+^ from the probe complex Cu^I^(Fz)2 non-competitively. Consequently, the Cu^+^ binding stoichiometries of the protein for those sites with sub-pM affinity were determined from the slope of a linear plot of the probe concentrations in the solution versus the added protein concentrations.

Protein variants H55A, M38L and C85S behaved similarly. Titration of protein variants C21S, C_107S_, C111S, C21/85S and C21/107/111S into the same Cu^I^(Fz)_2_ solution led to a solution with less or no precipitation. After the centrifugation, the solution absorbance after each titration also decreased linearly and consequently, the Cu^+^-binding stoichiometries for these four proteins were determined also from the slope of a linear plot of the probe concentrations versus the added protein concentrations.

The complex Cu^I^(Bcs)_2_ buffers free Cu^+^ at sub-fM concentration range and was used to estimate the Cu^+^-binding affinities for those protein sites that compete for Cu^+^ with this probe effectively. Each protein, except the C107 variant, was proven to bind 2-3 equivalents of Cu^+^ with sub-pM affinity but only two with affinity at fM range. Consequently, the Cu^+^ affinity (expressed as the dissociation constant, *K*_D_) was estimated from a linear plot of the Hill equation 1 based on a two Cu binding model (*96*):

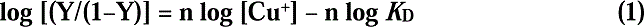

Where n is the Hill coefficient and Y is the fractional Cu^+^ occupancy of the total Cu^+^-binding sites (which is two for E2D2 with an intact CXXXC motif) under each experimental condition. The Y value was derived from the amount of Cu^+^ removed from the probe complex Cu^I^(Bcs)_2_ by the protein. The free [Cu^+^] was estimated from the dissociation equation of the remaining Cu^I^(Bcs)_2_ with its concentrations detected directly in various assay solutions.

The experiments were conducted by preparing a serious of solutions in KPi buffer (100 mM, pH 7.4) containing the same total probe components (20 µM Cu, 60 µM Bcs, 100 µM NH_2_OH, 100 µM GSH) with increasing amounts of protein (0 – 20 µM). The solution absorbance at 483 nm characteristic of Cu^I^(Bcs)_2_ was followed for each solution for 40-90 min until a steady value was read but without precipitation in each solution. Inclusion of GSH in the above assay facilitated the Cu^+^-transfer from the probe Cu^I^(Bcs)_2_ to the proteins but GSH at 100 µM was demonstrated to have no detectable competition for Cu^+^ under the above assay condition.

GSH binds Cu^+^ cooperatively as a Cu^I^ (GS) cluster and buffers free Cu^+^ (concentration expressed as pCu = – log [Cu^+^]) approximately according to equation 2 under the condition of pH 5.5-7.5 (*3*):

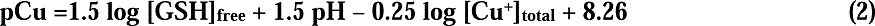

where [GSH]_free_ and [Cu(I)]_total_ refer to the concentrations of Cu^+^-unbound free GSH and total Cu^+^ bound by GSH, respectively. Accordingly, GSH is estimated to buffer free Cu^+^ at ∼fM concentration range with [GSH] = 100 µM but imposes free Cu^+^ to sub-fM range at [GSH] > 100 µM. Indeed, protein speciation analysis by NMR spectroscopy detected an effective competition for Cu^+^ between GSH (at > 100 µM) and protein variant E2D2_C21/85S_. This allows an independent evaluation of high-affinity Cu^+^-binding to E2D2_C21/85S_ using GSH as a Cu^+^-buffer and Cu^+^-affinity standard according to equation 2. The results are summarized in Figure 5H.

To ensure reliable protein speciation analysis and determination of Cu^+^-binding stoichiometries and affinities, the concentrations of all key reagents (proteins, GSH, Cu, Fz and Bcs) were calibrated and confirmed for each experiment. Protein structures were visualised with PyMOL. Protein sequence alignment was performed with Clustal-W.

### NMR spectroscopy and spectral analysis and assignments

NMR experiments were performed on ^13^C- and/or ^15^N-enriched 0.2 mM protein samples prepared in a N2-glovebox in thoroughly deoxygenated 100 mM KPi buffer (pH 7.4), 2 mM NH_2_OH, 1.0-1.5 mM GSH and 5% D_2_O. Cu (0.0-0.5 mM) was added as CuSO_4_ in pre-mixing with NH_2_OH/GSH to reduce Cu^2+^ to Cu^+^, stabilized as Cu^I^ (GS)_6_, prior to being added to each fully reduced protein. This procedure prevented protein oxidation and minimized Cu^+^-induced protein precipitation. Since the WT E2D2 protein with high Cu^+^ content tended to precipitate at a time scale of hours required for the NMR experiments, two representative protein variants E2D2_C21/85S_ and E2D2_C111S_ were selected for the NMR investigation. The C21S mutation has little impact on either the high-affinity Cu^+^-binding to E2D2 or on its Cu^+^-enhanced ubiquitin conjugation activity but the C111S mutation abolishes both the high-affinity Cu^+^-binding and the Cu^+^-enhancement of ubiquitin conjugation activity.

Spectra were recorded at 25 °C on a Bruker 700 AVIIIHD spectrometer equipped with a TCI cryoprobe. Three-dimensional (3D) HN(CO)CACB and HNCACB data were recorded to assign the ^1^H, ^15^N, ^13^Cα and ^13^Cβ resonances of ^13^C- and ^15^N-labeled E2D2_C21/85S_ in the absence and presence of Cu^+^. Spectra were recorded using 10% non-uniform sampling (NUS) and Poisson gap sampler(*97*), processed in NMRPipe (*98*) using SMILE(*99*) and analyzed using SPARKY NMRFAM (*100*). Chemical shift assignment of ^15^N-labeled E2D2_C111S_ was based on a comparison of the peak shift positions of the proteins. Due to large shift differences, peak overlap or ambiguity, the resonances of D42, I88, S111, D112, I119, A124 and Y127 could not be assigned in E2D2_C111S_. Chemical shift and intensity perturbations were monitored for ^15^N-labeled E2D2_C21/85S_ in the absence and presence of 0.5, 1.0, 1.5, 2.0 and 2.5 equivalents of Cu^+^; and for ^15^N-labeled E2D2_C111S_ in the absence and presence of 1.0 and 2.5-equivalents of Cu^+^. Weighted average chemical shift differences (Δδ_av_) of amide resonances, shown in Figure 6G and Figure S6E, were obtained from, where ΔδH and ΔδN are the ^1^HN and ^15^N chemical shift differences respectively between the presence and absence of Cu^+^. Intensity changes (I/I0) shown in Figure S6F for the apo-state peaks were determined using peak heights in the presence (I) and absence (I0) of Cu^+^. The fractions of allosteric states shown in Figure 6C were determined from the peak height of each discernible ^15^NH resonance state for N77 and L86 as a fraction of the sum of the peak heights of the apo state with the CuP and Cu2P states. The fractions of apoP and “CuP” for E2D2_C111S_ shown in Figure 6F were determined from the corresponding peak heights of the solution ^15^NH resonances for only L86 (because we could not detect a Cu2P species or a N77 response to Cu^+^ addition), expressed as a fraction of the peak height of the apoP species in the absence of Cu^+^. Since Cu^+^ addition led to partial E2D2_C111S_ precipitation, the apoP and CuP peak heights in the presence of Cu^+^ summed to less than the starting apoP height (2.5_eq_ < 1_eq_).

The main chain resonances (^1^H, ^15^N, ^13^Cα and ^13^Cβ) were assigned for the Cu-free form of E2D2_C21/85S_ using triple resonance experiments and taking advantage of the published wild-type assignments(*39*). Titration of Cu^I^ (GS) reagent into ^15^N-labelled E2D2 solution in KPi buffer (100 mM, pH 7.4) led to the appearance of new resonances in the ^1^H-^15^N HSQC spectra assignable to the CuP and Cu_2_P species (Figure 6A-B), as highlighted for the well-resolved N77 and L86 (Figure 6C). Sampling conditions were controlled to allow maximal resonance assignments for the CuP species, as shown in Figure 6A. Titration with the Cu^I^ (GS) reagent generated new peaks. While well-resolved resonances were readily assigned (e.g., N77 and L86), those in crowded regions were sufficiently ambiguous to require reassignment. Reassignment was possible for many “split” peaks, correlations between the ^1^H-^15^N to the ^13^Cα and ^13^Cβ were generally weak suggesting conformational heterogeneity. We note that in the absence of Cu^+^, ^13^Cα/β correlations for the apo-state HN resonances of I106 to D112, that includes the Cu^+^-binding residues C107 and C111, were strong and easily assigned. In the presence of Cu^+^, we could not assign any correlations for the original peak that corresponds to the apo-state or identify new peaks of the CuP state. However, it is likely that these resonances are split as the peak that corresponds to the apo-state is reduced to 30 to 40%, similar to resonances that show clear resolvable splitting in the presence of Cu^+^ (Figure S6F).

Control experiments with the E2D2_C111S_ protein variant detected minimal chemical shift changes induced by Cu^+^ for any resonances (Figure 6D-E), even when the Cu^+^ concentrations were high enough to induce detectable protein precipitation (Figure 6F). Protein structure figures were generated with the program PyMOL.

### Evolutionary analysis

Sequence alignment of human E2s was performed with Clustal-W. Alignment of E2D2 orthologues was performed with Basic Local Alignment Search Tool (BLAST) (*102*) using the blastp algorithm with BLOSUM62 (default) scoring. Fast Minimum Evolution (*103*) trees based on Grishin (*104*) protein distance from reference sequence were generated using BLAST (*102*). For studying E2D2 orthologues among holozoa, representative species from major multicellular animal lineages, including non-bilaterians (placozoa, porifera, cnidaria) and bilaterians (nematodes, arthropods, vertebrates), and from three independent, closely related, unicellular holozoan lineages (choanoflagellate, filasterean and ichthyosporea) were assessed using human E2D2 (NP_003330.1) as reference. Timing of the emergence of the CXXXC motif was estimated based on its presence in major animal lineages yet absence in unicellular holozoans. For studying E2D orthologues among eukaryotes, representative species from major eukaryotic lineages (excavate, Amoebozoa, Sar, holomycota, Archaeplastida and holozoans) were assessed using human E2D2 (NP_003330.1) as reference. For studying the relationship between vertebrate E2D paralogues and their fly orthologue, E2D1-3 from representative species across five vertebrate classes were assessed using *D. melanogaster* UbcD1 (CAA44453.1) as reference.

### Drosophila Studies

#### *In vivo* gene manipulation

*UAS-RNAi* lines targeting *UbcD1* / *effete* were sourced from the Vienna *Drosophila* Resource Centre (ID numbers 110767 and 26011) and the Bloomington *Drosophila* Stock Center (ID number 35431). GAL4 lines used in this study were: *Pannier-GAL4* (BL3039, BL=Bloomington *Drosophila* Stock Center, Bloomington, IN, USA), *Tubulin-GAL4* (BL5138), *Eyeless-GAL4* (BL5535), *GMR-GAL4* (BL8121), *CCAP-GAL4* (gift from B. White, NIH, USA) and *Elav-GAL4* (BL458). *UAS-E2D2-WT* and *-E2D2-C111S* transgenes were generated by subcloning full-length E2D2-WT and E2D2-C111S cDNAs into the pUAST-attB vector. These constructs were injected into the *PhiC31 attP 51C* strain (provided by K. Basler, University of Zurich, Switzerland). Microinjections utilized an Eppendorf Femtojet apparatus with Femtotips II pre-pulled glass needles (Eppendorf). All *Drosophila melanogaster* strains and crosses were maintained on standard medium at 25 °C unless stated otherwise. All images shown are representative of >20 individuals of the same genotype.

### Apoptosis - immunostaining against active Caspase-3

Fly brains were dissected from newborn adult flies or pharate pupae, and directly fixed for 30 min in 4% paraformaldehyde (PFA) at 22 °C. After three × five-minute washes in phosphate-buffered saline (PBS), samples were rinsed with 0.5% Triton X-100 containing PBS (PBT) three times (20 min per time), followed by blocking with 5% goat serum in PBT at 22 °C for 1 h. They were incubated with polyclonal rabbit anti-Caspase 3 that detects the active form of caspase-3 (cleaved form), but not the inactive (full-length) form of caspase-3 (1:200; Cell Signaling Technology, Danvers, MA, USA), overnight at 4 °C. Anti-rabbit 488 (1:500; Invitrogen, Carlsbad, CA, USA) secondary antibody was used to detect primary antibodies. After DAPI staining (1 μg/ml, Bio-Rad, Hercules, CA, USA), adult fly brains were mounted in PermaFluor (Thermo Scientific, Fremont, CA, USA) for confocal microscopy (CellVoyager CV1000, Yokogawa, Tokyo, Japan).

### Statistics

For biochemical and functional experiments, independent samples *t*-tests, one-way and two-way ANOVAs were performed, depending on the context of biological questions addressed, followed by relevant post-hoc tests. When variances were not equal, results of robust tests were also assessed.

### Data Availability

For ubiquitomics and pathway analyses, raw data has been provided in the Supplemental Information (Tables S1-S5). For evolutionary analyses, raw data has been provided in the Supplemental Information (Table S7).

## SUPPLEMENTAL INFORMATION

### Supplementary Figures

Figure S1. Copper promotes protein ubiquitination across different cells, related to Figure 1.

Figure S2. Copper promotes protein degradation but does not lead to changes in cell survival, oxidative stress, or proteasome activity, related to Figures 1-2.

Figure S3. Bioinformatic workflow for MS ubiquitomics in NSC34 cells treated with copper, related to Figures 3-4.

Figure S4. p53 is a major target for Cu^+^-enhanced ubiquitination and degradation, related to Figure 3.

Figure S5. Cu^+^ enhances protein ubiquitination and degradation via E2D conjugases, related to Figure 4.

Figure S6. Sub-femtomolar-affinity Cu^+^ binding at the C107XXXC111 motif triggers conformational changes that allosterically activate E2D2, related to Figures 5 and 6.

Figure S7. Potential Cu^+^ ligands across human E2s and evolutionary trees of E2D orthologues in eukaryotes and in vertebrates, related to Figure 7.

### Supplementary Tables

Table S1. Ubiquitomics – MaxQuant parameters, MaxQuant output, filtering and annotation (Excel Table), related to Figures 3 and S3.

Table S2. Ubiquitomics – Quantifying relative abundance using Perseus (Excel Table), related to Figures 3 and S3.

Table S3. Ubiquitomics – Enriched ontology terms (Excel Table), related to Figures 3 and S3.

Table S4. Ubiquitomics – The interactome of Cu^+^-enhanced Ub-associated proteins (Excel Table), related to Figures 3 and S3-S4.

Table S5. Ubiquitomics – Enrichment of Cu^+^-enhanced proteins among predicted substrates of selected E2 clades (Excel Table), related to Figures 4 and S3-S4.

Table S6. Determination of Cu^+^-binding affinity by protein speciation analysis of NMR shifts, related to Figures 6 and S6.

Table S7. Evolution – Evolutionary tree of E2D orthologues among eukaryotes and among holozoans and of E2D paralogues among vertebrates (Excel Table), related to Figures 7 and S7.

## Notes

### Summary of Updates

Title, Summary, Introduction, Discussion, Figures and Supplementary Information were revised and updated.

