## Supplemental Information for "Nutrient copper signaling promotes protein turnover by allosteric activation of ubiquitin E2D conjugases"

#### Supplementary Figures

**Figure S1**. Copper promotes protein ubiquitination across different cells, related to Figure 1.

**Figure S2**. Copper promotes protein degradation but does not lead to changes in cell survival, oxidative stress, or proteasome activity, related to Figures 1-2.

**Figure S3.** Bioinformatic workflow for MS ubiquitomics in NSC34 cells treated with copper, related to Figures 3-4.

**Figure S4**. p53 is a major target for $Cu^{+}$-enhanced ubiquitination and degradation, related to Figure 3.

**Figure S5**. $Cu^{+}$ enhances protein ubiquitination and degradation via E2D conjugases, related to Figure 4.

**Figure S6**. Sub-femtomolar-affinity $Cu^{+}$ binding at the $C_{107}XXXC_{111}$ motif triggers conformational changes that allosterically activate E2D2, related to Figures 5 and 6.

**Figure S7**. Potential $Cu^{+}$ ligands across human E2s and evolutionary trees of E2D orthologues in eukaryotes and in vertebrates, related to Figure 7.

#### Supplementary Tables

**Table S1**. Ubiquitomics – MaxQuant parameters, MaxQuant output, filtering and annotation (Excel Table), related to Figures 3 and S3.

**Table S2**. Ubiquitomics – Quantifying relative abundance using Perseus (Excel Table), related to Figures 3 and S3.

**Table S3**. Ubiquitomics – Enriched ontology terms (Excel Table), related to Figures 3 and S3.

**Table S4**. Ubiquitomics – The interactome of $Cu^{+}$-enhanced Ubiquitin-associated proteins (Excel Table), related to Figures 3 and S3-S4.

**Table S5**. Ubiquitomics – Enrichment of $Cu^{+}$-enhanced proteins among predicted substrates of selected E2 clades (Excel Table), related to Figures 4 and S3-S4.

**Table S6**. NMR - Determination of $Cu^{+}$-binding affinity by protein speciation analysis of NMR shifts, related to Figures 6 and S6.

**Table S7**. Evolution – Evolutionary tree of E2D orthologues among eukaryotes and among holozoans and of E2D paralogues among vertebrates (Excel Table), related to Figures 7 and S7.

Figure S1

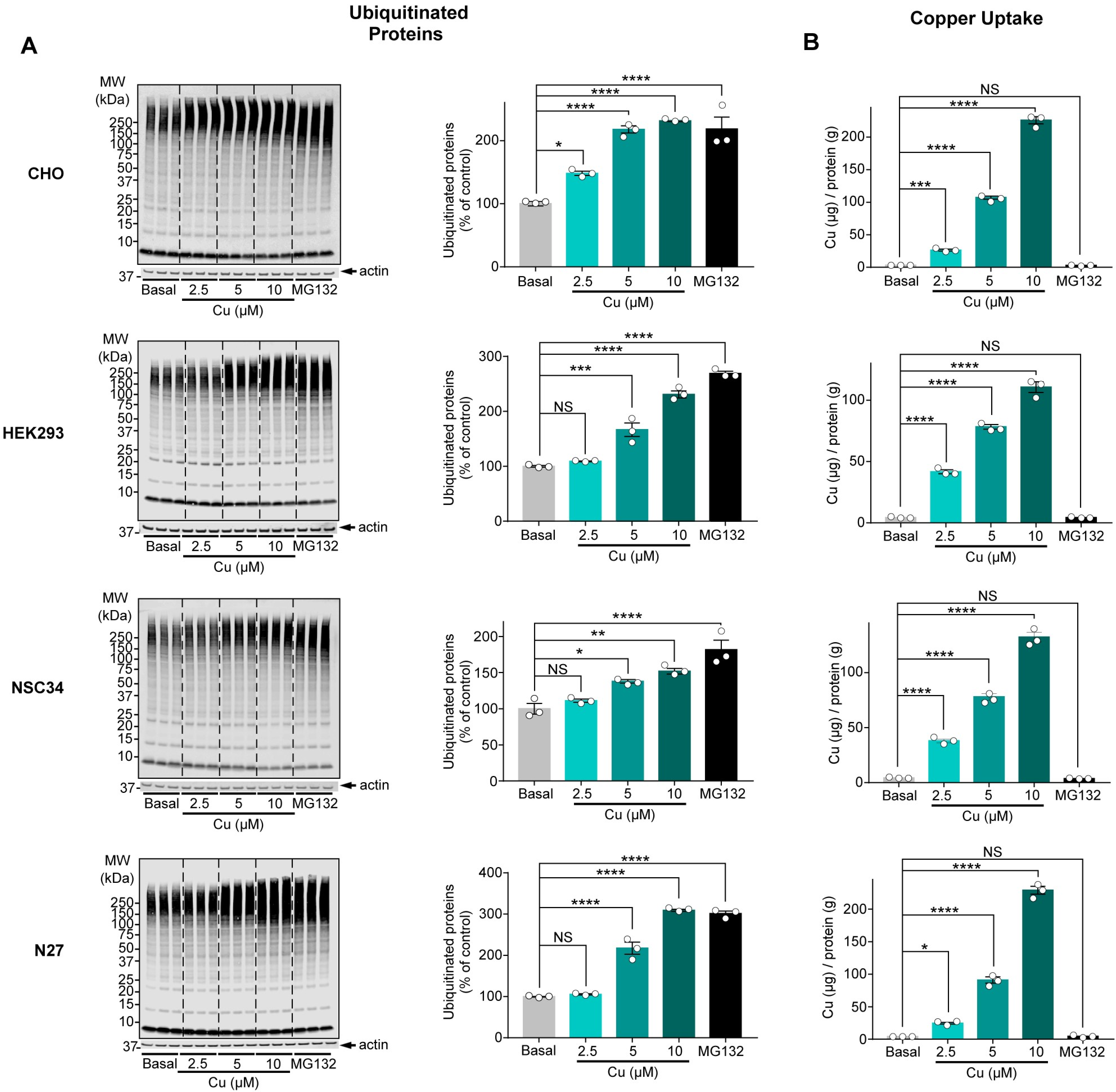

A
B
Ubiquitinated Proteins
Copper Uptake
CHO
HEK293
NSC34
N27
MW (kDa)
250
150
100
75
50
37
25
20
15
10
actin
Basal
2.5
5
10
MG132
Cu (μM)
Ubiquitinated proteins (% of control)
Cu (μg) / protein (g)
NS
*
**
***
****

**Figure S1. Copper promotes protein ubiquitination across different cells, related to Figure 1.**

(A) Ubiquitination response to copper supplementation in various cell lines: Chinese hamster ovary (CHO) cells, HEK293 cells, NSC34 cells and N27 cells. Cells were untreated (basal) or supplemented with $CuCl_2$ (Cu; up to 10 μM) for 3 h in Locke's media. Ubiquitinated proteins were detected by blot (P4D1 antibody). Actin was used as loading control.

(B) Cellular copper levels following supplementation.

Bar graphs in A-B depict means ± SEM (n = 3). NS=non-significant, **P*<0.05, ***P*<0.01, ****P*<0.001, *****P*<0.0001, ANOVA followed by Dunnett's test.

### Figure S2

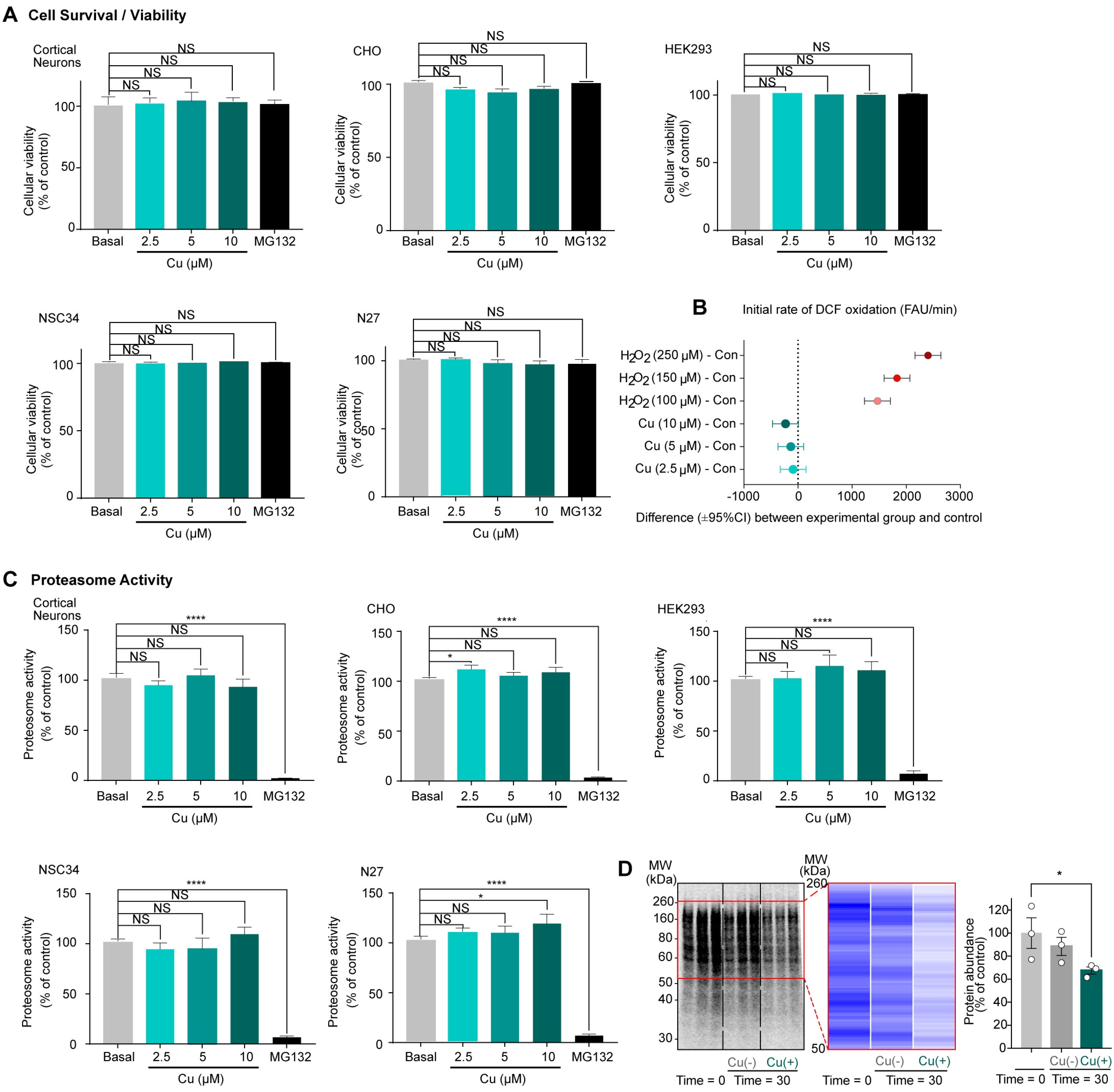


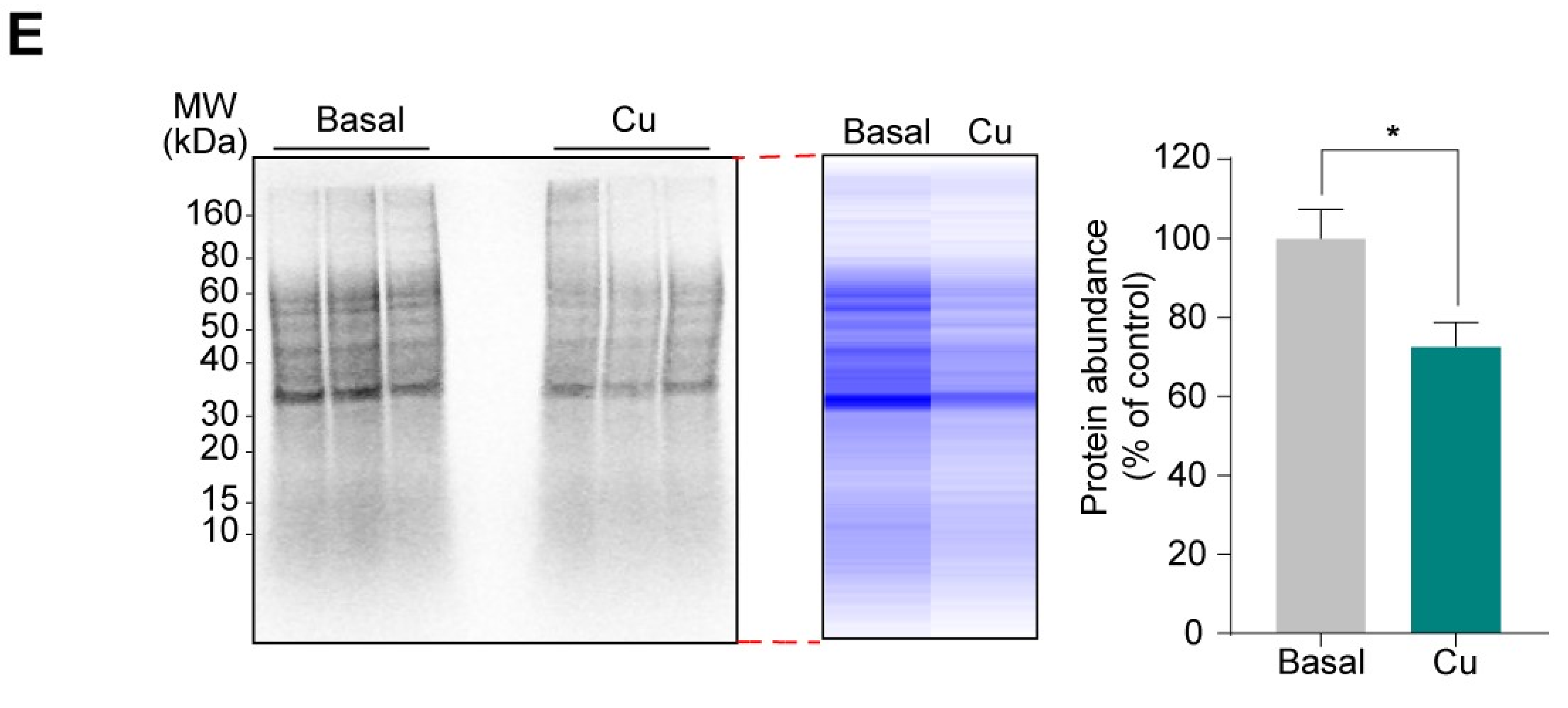


F

| Cell type | Cu-enhanced ubiquitination | Cu-promoted protein degradation |
|---|---|---|
| Cortical neurons | ✓ | ✓ |
| NSC34 | ✓ | ✓ |
| MEF | ✓ | ✓ |
| HEK293T | ✓ | NA |
| CHO | ✓ | NA |
| N27 | ✓ | NA |
| N2a | ✓ | NA |
| HepG2 | ✓ | NA |
| HeLa | ✓ | NA |
| Hap1 | ✓ | ✓ |
| HEK293 | ✓ | ✓ |
| Human fibroblasts | ✓ | ✓ |
| Mouse fibroblasts | ✓ | ✓ |

**Figure S2. Copper promotes protein degradation but does not lead to changes in cell survival, oxidative stress, or proteasome activity, related to Figures 1-2.**

(A) Cells were untreated (basal) or supplemented with $CuCl_2$ (Cu; up to 10 μM) for 3 h in Locke's media. Cell survival was measured using MTT assay (primary mouse cortical cultures) or Trypan Blue exclusion (CHO, HEK293, NSC34 and N27). Bar graphs depict means ± SEM (n = 3). NS=non-significant, ANOVA followed by Dunnett's test.

(B) Initial rate of DCF oxidation (change in fluorescence arbitrary units [FAU] / min) in HEK293 cells exposed to copper (Cu; up to 10 μM) or to hydrogen peroxide ($H_2O_2$; up to 250 μM). The graph shows the mean difference (±95% CI, n = 4) between each experimental condition and control, based on ANOVA followed by Dunnett's test.

(C) Proteasome activity in the same cells studied in panel A. MG132 (10 μM) was used as a positive control of proteasome activity inhibition. A combined proteasome activity reflecting its three major catalytic components (caspase-, trypsin- and chymotrypsin-like) is presented (see Methods). Bar graphs depict means ± SEM (n=3). NS=non-significant, $^{*}P<0.05$, $^{****}P<0.0001$, ANOVA followed by Dunnett's test.

(D) In this two-time-point independent replication of the experiment presented in Figure 2B, NSC34 cells were pulse-labelled with $^{35}$S-Met in Cys/Met-free media, followed by a chase incubation in unlabelled media containing the protein-synthesis inhibitor cycloheximide (1 mg/mL) ± $CuCl_2$ (10 μM) for 30 min. $^{35}$S-positive bands within NP40 cell extracts were detected in a Typhoon system. The gel shows the autoradiography protein profile observed in the non-supplemented (Cu-) or copper-supplemented (Cu+) cells. Quantitative analysis of the $^{35}$S-positive bands detected at nominal time 0 and after 30 min in the same cells. The red frame encloses the region of interest, transformed as an intensity heatmap for 95 equal bins averaged across each experimental condition. Bar graphs depict means ± SEM (n=3). *$P$<0.05, ANOVA followed by test for linear trend.

(E) MEF cells were pulse-labelled with $^{35}S$ in Cys/Met-free media and protein degradation was analysed after 30 min, as above. Bar graphs depict means ± SEM (n=3). *$P$<0.05, *t*-test for independent samples.

(F) Table summarizes the cells tested for $Cu^{+}$-enhanced ubiquitination and $Cu^{+}$-promoted degradation. NA=not assessed.

### Figure S3

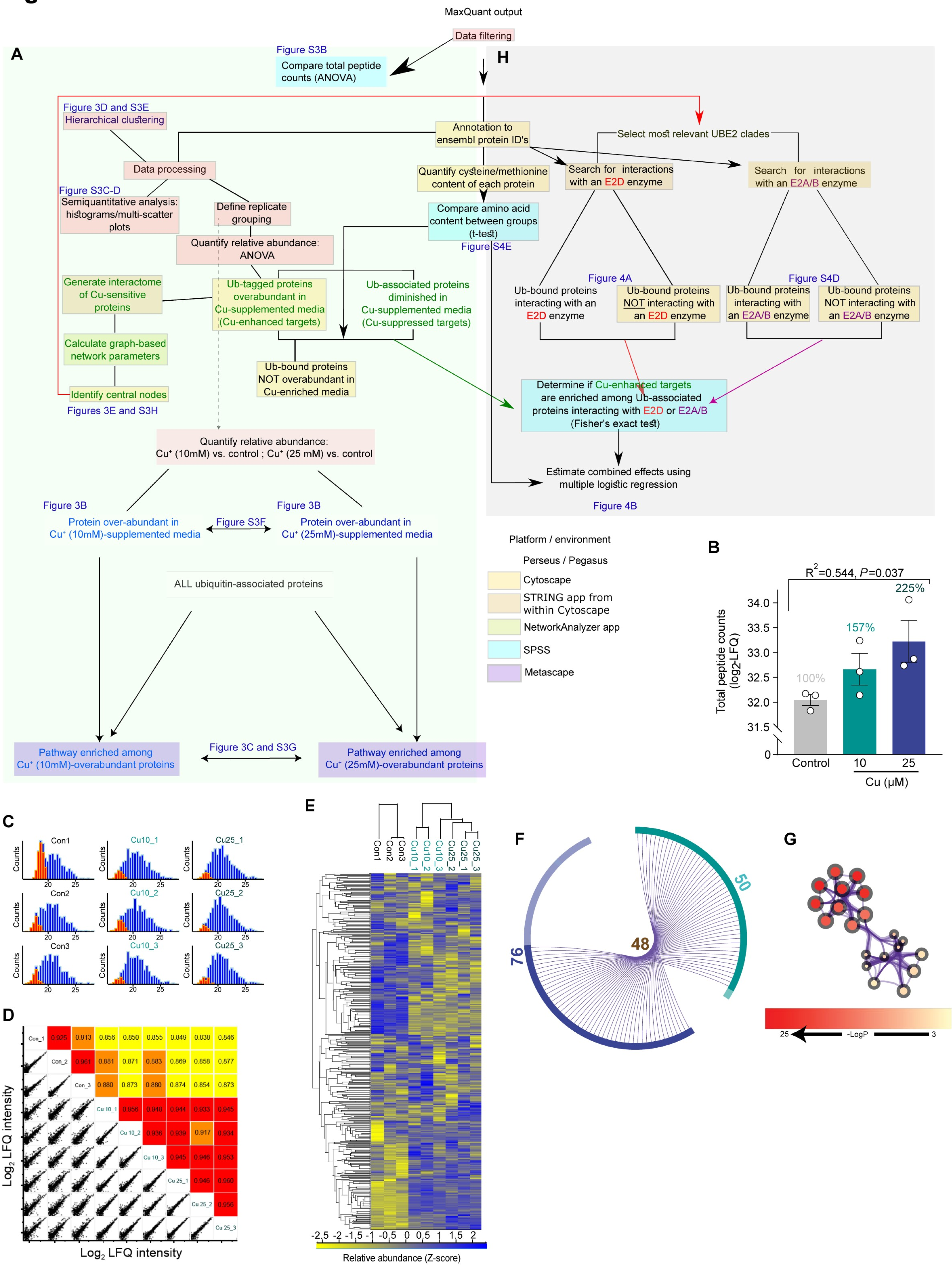

MaxQuant output
Data filtering
A
Figure S3B
Compare total peptide counts (ANOVA)
H
Figure 3D and S3E
Hierarchical clustering
Annotation to ensembl protein ID's
Select most relevant UBE2 clades
Data processing
Quantify cysteine/methionine content of each protein
Search for interactions with an E2D enzyme
Search for interactions with an E2A/B enzyme
Figure S3C-D
Semiquantitative analysis: histograms/multi-scatter plots
Define replicate grouping
Compare amino acid content between groups (t-test)
Figure S4E
Quantify relative abundance: ANOVA
Generate interactome of Cu-sensitive proteins
Ub-tagged proteins overabundant in Cu-supplemented media (Cu-enhanced targets)
Ub-associated proteins diminished in Cu-supplemented media (Cu-suppressed targets)
Figure 4A
Ub-bound proteins interacting with an E2D enzyme
Ub-bound proteins NOT interacting with an E2D enzyme
Figure S4D
Ub-bound proteins interacting with an E2A/B enzyme
Ub-bound proteins NOT interacting with an E2A/B enzyme
Calculate graph-based network parameters
Ub-bound proteins NOT overabundant in Cu-enriched media
Identify central nodes
Figures 3E and S3H
Determine if Cu-enhanced targets are enriched among Ub-associated proteins interacting with E2D or E2A/B (Fisher's exact test)
Quantify relative abundance: Cu⁺ (10mM) vs. control ; Cu⁺ (25 mM) vs. control
Estimate combined effects using multiple logistic regression
Figure 4B
Figure 3B
Protein over-abundant in Cu⁺ (10mM)-supplemented media
Figure S3F
Figure 3B
Protein over-abundant in Cu⁺ (25mM)-supplemented media
Platform / environment
Perseus / Pegasus
Cytoscape
STRING app from within Cytoscape
NetworkAnalyzer app
SPSS
Metascape
ALL ubiquitin-associated proteins
Pathway enriched among Cu⁺ (10mM)-overabundant proteins
Figure 3C and S3G
Pathway enriched among Cu⁺ (25mM)-overabundant proteins
B
R²=0.544, P=0.037
Total peptide counts (log₂-LFQ)
34.0
33.5
33.0
32.5
32.0
31.5
0
100%
157%
225%
Control
10
25
Cu (μM)
C
Con1
Con2
Con3
Cu10_1
Cu10_2
Cu10_3
Cu25_1
Cu25_2
Cu25_3
Counts
20
25
D
Con_1 0.925 0.913 0.856 0.850 0.855 0.849 0.838 0.846
Con_2 0.961 0.881 0.871 0.883 0.869 0.858 0.877
Con_3 0.880 0.873 0.880 0.874 0.854 0.873
Cu 10_1 0.956 0.946 0.944 0.933 0.945
Cu 10_2 0.936 0.939 0.917 0.934
Cu 10_3 0.945 0.946 0.953
Cu 25_1 0.946 0.960
Cu 25_2 0.956
Cu 25_3
Log₂ LFQ intensity
Log₂ LFQ intensity
E
Con1
Con2
Con3
Cu10_1
Cu10_2
Cu10_3
Cu25_2
Cu25_1
Cu25_3
-2,5 -2 -1,5 -1 -0,5 0 0,5 1 1,5 2
Relative abundance (Z-score)
F
50
48
76
G
25
-LogP
3

**Figure S3. Bioinformatic workflow for MS ubiquitomics in NSC34 cells treated with copper, related to Figures 3-4.**

(A) Workflow for assessing the effects of copper treatment on total peptide count, quality control, quantifying relative abundance of ubiquitin-associated proteins, identifying $Cu^{+}$-enhanced ubiquitin-associated proteins and central nodes in the $Cu^{+}$-enhanced ubiquitome and pathway enrichment among $Cu^{+}$-overabundant proteins.

(B) Bar graph depicting the effects of copper supplementation on the total ubiquitin-associated peptides captured and quantified by mass spectrometry. Bars represent mean ±SEM with three replicates per experimental condition. Significance based on ANOVA linear term.

(C) Label free quantification (LFQ) histograms. Protein intensities were calculated for each replicate and experimental condition. The distribution of log-transformed imputed values (orange) was created using default imputation parameters (a down-shift of 1.8 and distribution width of 0.3 relative to the distribution of the log-transformed observed values [blue]), to simulate the assumption of low abundant proteins giving rise to missing values. Con, control; Cu10, $CuCl_2$ 10 μM; Cu25, $CuCl_2$ 25 μM.

(D) Multi-scatter plots (below diagonal) and Pearson's correlations (above diagonal, colour-coded according to correlation coefficient) between each pair of replicates within and between experimental conditions.

(E) Heatmap depicting the relative abundance (standardised for each protein) of all 882 ubiquitin-associated proteins reliably identified by the ubiquitomics screen. Of these, 66 proteins with a relative abundance significantly altered by copper are presented in Figure 3A. Dendrograms depict hierarchical clustering according to copper challenge and protein abundance profile (rows).

(F) Circos plots displaying overlap between proteins that were over-abundant in each of two $CuCl_2$-supplemented conditions (as visualised in Figure 3B). The circle represents protein lists, where hits are arranged along the arc (10 μM $CuCl_2$-supplementation, cyan; 25 μM $CuCl_2$-supplementation, blue). Proteins that hit both lists are coloured in solid colours, while proteins unique to a list are shown in transparent colours. Purple curves link identical proteins. Adjacent to arcs are the numbers of proteins in each list, with the number of overlapping proteins denoted inside plot.

(G) Same network as Figure 3C, with node colours representing significance ($-\log_{10}P$) of each term's enrichment, i.e., its (over)representation among hits compared to the background list (all ubiquitin-associated proteins detected in mass-spectrometry).

### Figure S4

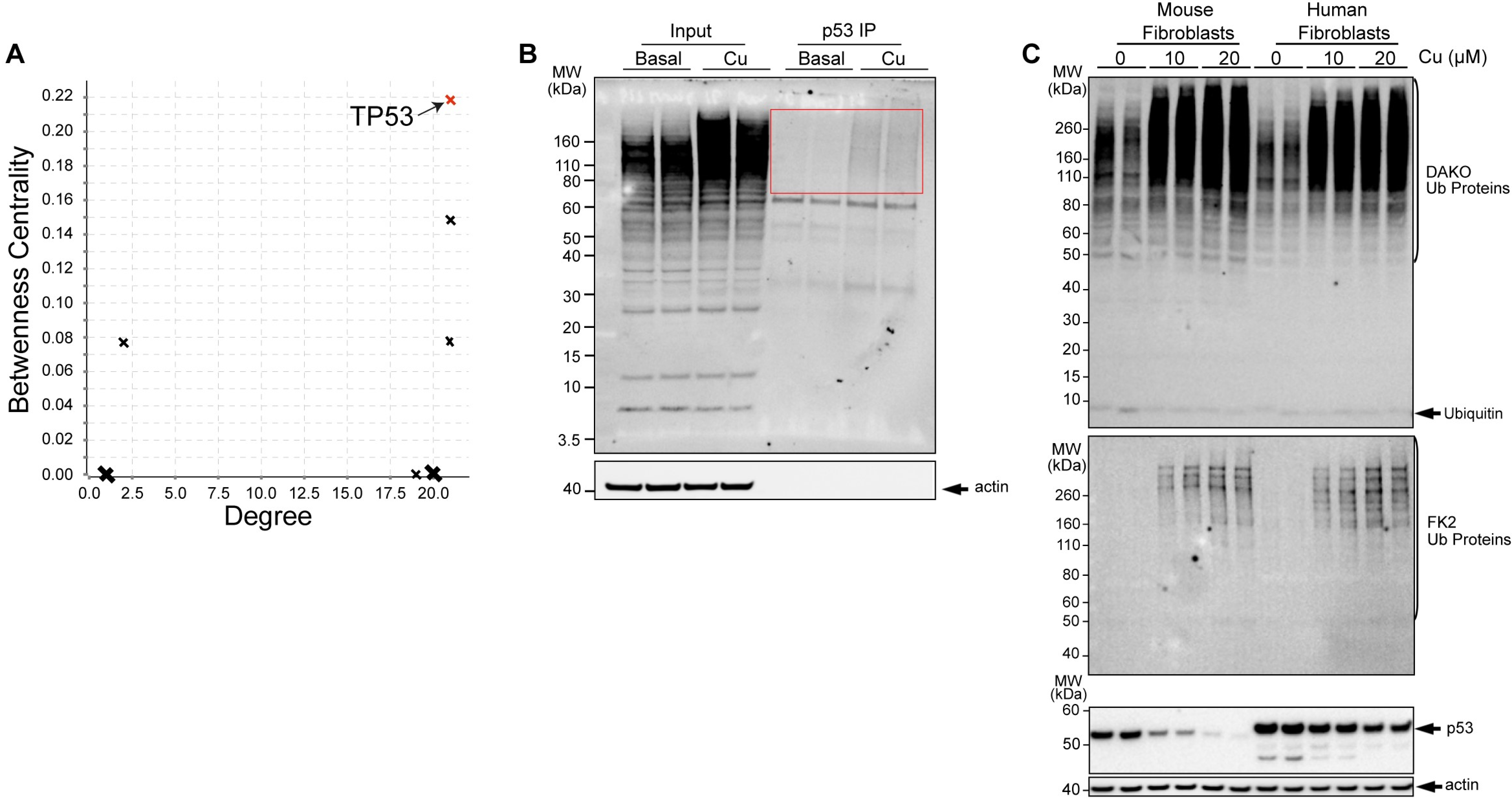
A
TP53
Betweenness Centrality
0.22
0.20
0.18
0.16
0.14
0.12
0.10
0.08
0.06
0.04
0.02
0.00
0.0 2.5 5.0 7.5 10.0 12.5 15.0 17.5 20.0
Degree
B
Input
p53 IP
Basal
Cu
Basal
Cu
MW (kDa)
160
110
80
60
50
40
30
20
15
10
3.5
40
actin
C
Mouse Fibroblasts
Human Fibroblasts
0 10 20 0 10 20 Cu (μM)
MW (kDa)
260
160
110
80
60
50
40
30
20
15
10
DAKO Ub Proteins
Ubiquitin
FK2 Ub Proteins
p53
actin

**Figure S4. p53 is a major target for $Cu^{+}$-enhanced ubiquitination and degradation, related to Figure 3.**

(A) Plot depicting node-degree (*x*-axis) versus betweenness centrality (*y*-axis) of 27 ubiquitin-associated proteins whose relative abundance was significantly enhanced by copper supplementation and which formed a connected subnetwork based on protein-protein interaction data curated from the STRING database ($Cu^{+}$-enhanced interactome, Figure 3D). The node representing p53 (in red) displayed an unusually high value of betweenness centrality, reflecting its interactions with both proteasomal and non-proteasomal proteins.

(B) Copper promotes ubiquitination of p53. Ubiquitin-tagged p53 was immunoprecipitated (anti-p53) from HEK293 cells incubated ± $CuCl_2$ (10 μM) for 3 h and detected by Western blot (FK2 antibody). The area of the blot within the red box is presented in Figure 3E.

(C) Copper promotes polyubiquitination and p53 degradation in mouse and human fibroblasts. Cells were treated ± $CuCl_2$ (10 μM) for 3 h. The $Cu^{+}$-enhanced p53 degradation (lower blot) inversely correlated with the formation of ubiquitinated proteins detected by DAKO (upper blot) and FK2 (middle blot) antibodies. The blots show data from duplicate cultures.

### Figure S5

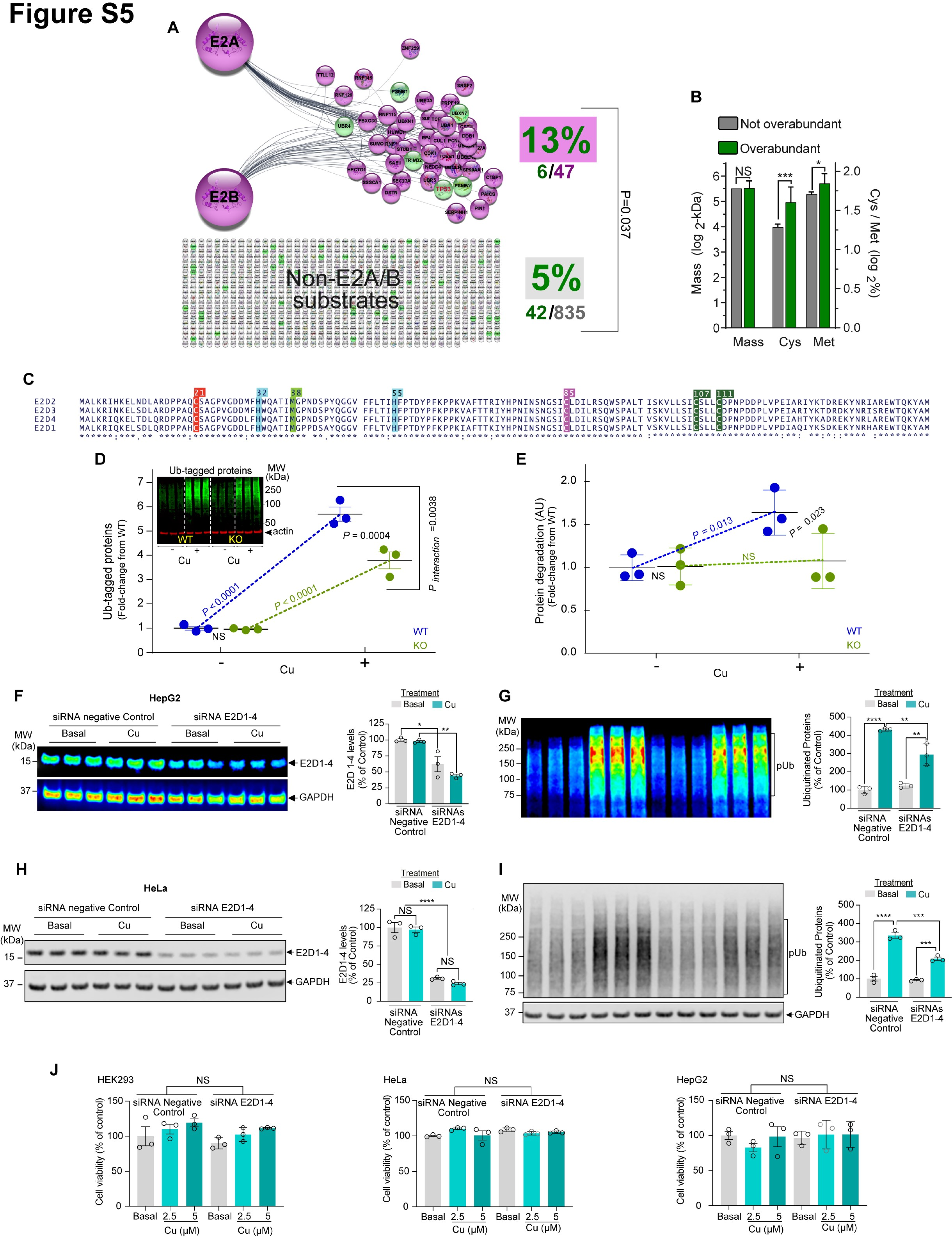
A
E2A
E2B
13%
6/47
Non-E2A/B substrates
5%
42/835
P=0.037
B
Not overabundant
Overabundant
Mass (log 2-kDa)
Cys / Met (log 2%)
NS
***
*
Mass
Cys
Met
C
E2D2
E2D3
E2D4
E2D1
D
Ub-tagged proteins
MW (kDa)
250
100
50
actin
WT
KO
Cu
Ub-tagged proteins (Fold-change from WT)
P<0.0001
P = 0.0004
P interaction =0.0038
NS
WT
KO
Cu
E
Protein degradation (AU) (Fold-change from WT)
P = 0.013
P = 0.023
NS
WT
KO
Cu
F
HepG2
siRNA negative Control
siRNA E2D1-4
Basal
Cu
MW (kDa)
E2D1-4
GAPDH
Treatment
Basal
Cu
E2D 1-4 levels (% of Control)
siRNA Negative Control
siRNAs E2D1-4
G
MW (kDa)
pUb
Ubiquitinated Proteins (% of Control)
H
HeLa
siRNA negative Control
siRNA E2D1-4
Basal
Cu
E2D1-4
GAPDH
E2D1-4 levels (% of Control)
NS
****
I
pUb
GAPDH
Ubiquitinated Proteins (% of Control)
J
HEK293
HeLa
HepG2
siRNA Negative Control
siRNA E2D1-4
Cell viability (% of control)
Basal
Cu (μM)
NS

**Figure S5. $Cu^+$ enhances protein ubiquitination and degradation via E2D conjugases, related to Figure 4.**

(A) Clade over-representation analysis comparing the proportion of ubiquitin-associated proteins displaying $Cu^+$-enhanced overabundance (green nodes) among STRING-predicted E2A or E2B substrates (purple nodes) with the corresponding proportion among non-E2A/B substrates (grey nodes). Copper caused the enhanced ubiquitination of 5% of all 835 proteins captured, but of the 47 E2A/B substrates captured, copper caused the enhanced ubiquitination of 13%, yielding a marginally significant enrichment (Fisher's exact test).

(B) Bar graph depicting protein mass (left *y* axis, $log_2$-kDa) and cysteine or methionine content (right *y* axis, $log_2$-% of total amino acids) for proteins that did (n=48), or did not (n=834), display overabundance in copper-supplemented media. Bars represent mean ± 95% CI. *$p<0.05$ and ***$p<0.001$, non-paired *t*-tests.

(C) Sequence alignment of E2D1-4 enzymes. The C85 is the enzyme active site. Other highlighted residues are potential $Cu^+$ ligands. Residues shared by all four paralogues are denoted with an asterisk.

(D) $Cu^+$-enhanced ubiquitination is blunted in E2D2 KO cells. Hap1 Parental (WT) and E2D2 knockout cells (KO) were treated ± Cu (10 μM) for 3 h. Insert - blot of ubiquitinated proteins. Graph - ubiquitinated protein levels for each condition (mean ± SEM), n=3. *P* values are from two-way ANOVA followed by pairwise comparisons, showing that protein ubiquitination in E2D2 KO cells is both decreased in magnitude and less responsive to copper than WT.

(E) $Cu^+$-enhanced protein degradation is largely driven by E2D2. The same cells were pre-incubated with $^{35}$S-Cys/Met for 1 h, then washed and treated ± Cu (25 μM) for 15 min. The graph shows an index of protein degradation (peptide degradation fragment counts normalized by cell lysate counts). *P* values are from two-way-ANOVA followed by pairwise comparisons. NS_non-significant.

(F) HepG2 cells were treated with control siRNA or siRNA against E2D1-4 for 48h. Then cells were untreated (basal) or supplemented with $CuCl_2$ (Cu; 5 μM) for 3 h in Locke's media. The levels of E2D1-4 were detected with an antibody that recognizes the full E2D clade.

(G) Ubiquitinated proteins (pUb) were detected by blot (P4D1 antibody). GAPDH was used as loading control.

(H) HeLa cells were treated with siRNAs under the same conditions described in "D". Then cells were untreated (basal) or supplemented with $CuCl_2$ (Cu; 2.5 μM) for 3 h in Locke's media. The levels of E2D1-4 were detected with an antibody that recognizes the full E2D clade.

(I) Ubiquitinated proteins (pUb) were detected by blot (P4D1 antibody). GAPDH was used as loading control.

Bar graphs in F-I depict means ± SEM (n = 3). * $P<0.05$, ** $P<0.01$, *** $P<0.001$, **** $P<0.0001$, two-way-ANOVA followed by Bonferroni's tests.

(J) Cell viability was measured using Presto blue assay in HeLa, HepG2 and HEK cells treated under the same conditions described in Fig. 3, B and C, and fig. S7, C and D. Data are means ± SEM (n=3). NS=non-significant, ANOVA followed by Dunnett's test.

### Figure S6

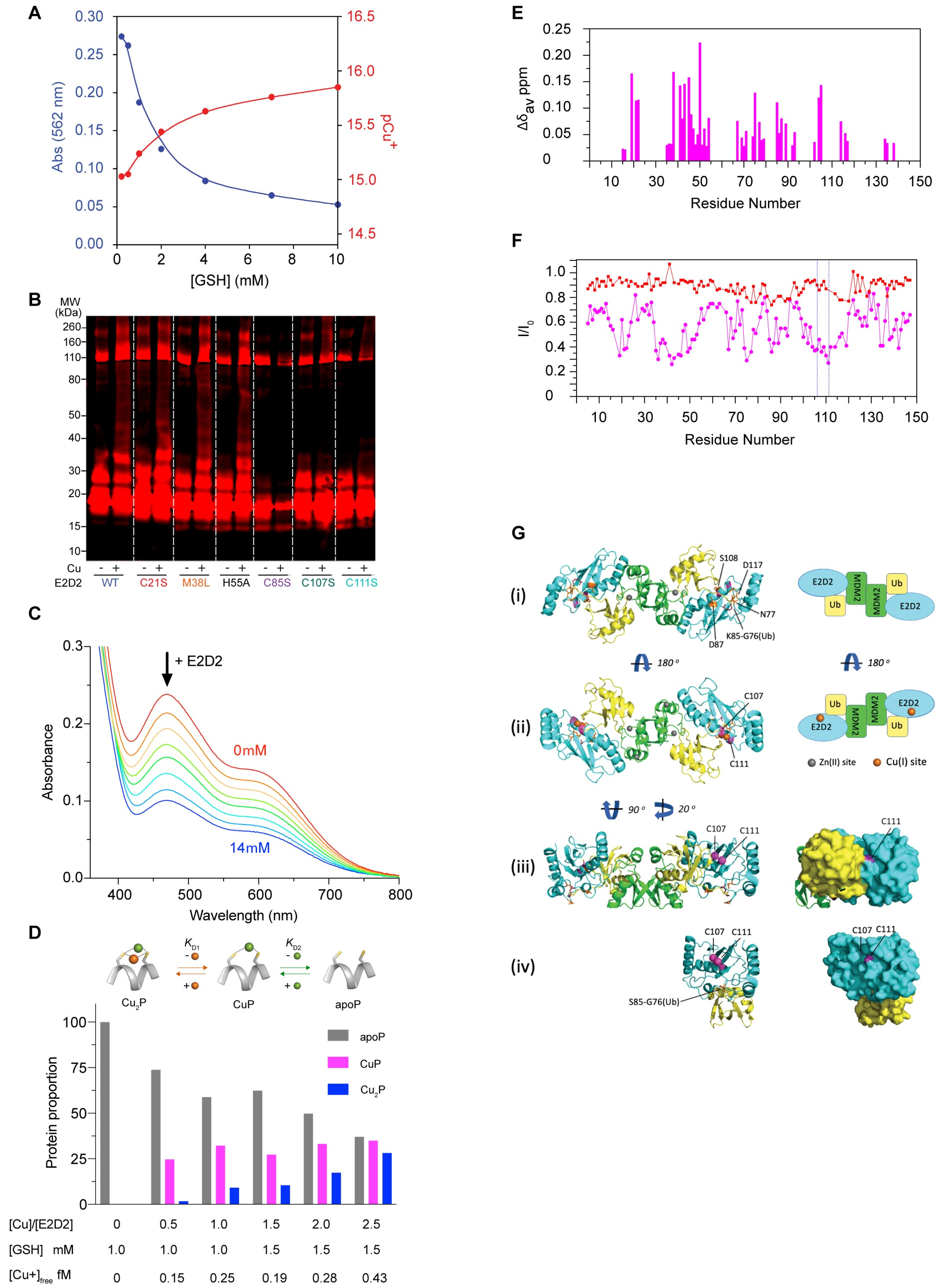

A
Abs (562 nm)
pCu+
[GSH] (mM)
B
MW (kDa)
260
160
110
80
50
40
30
20
15
10
Cu
E2D2
WT
C21S
M38L
H55A
C85S
C107S
C111S
C
+ E2D2
0mM
14mM
Absorbance
Wavelength (nm)
D
$K_{D1}$
$K_{D2}$
$Cu_2P$
CuP
apoP
Protein proportion
[Cu]/[E2D2]
[GSH] mM
$[Cu+]_{free}$ fM
E
$\Delta\delta_{av}$ ppm
Residue Number
F
$I/I_0$
Residue Number
G
(i)
(ii)
(iii)
(iv)
S108
D117
N77
K85-G76(Ub)
D87
C107
C111
S85-G76(Ub)
E2D2
MDM2
Ub
180 º
90 º
20 º
Zn(II) site
Cu(I) site

**Figure S6. Sub-femtomolar-affinity $Cu^+$ binding at the $C_{107}XXXC_{111}$ motif triggers conformational changes that allosterically activate E2D2, related to Figures 5 and 6.**

(A) Free [$Cu^+$] concentrations (as $pCu^+ = -\log [Cu^+]$) buffered by GSH (0.2 – 10 mM). The free [$Cu^+$] in a system containing $Cu^+$ (total 35 μM) and GSH (0.2 – 20 mM) in Na-MOPS buffer (50 mM, pH 7.3) was detected by $Cu^+$ indicator Bca (total 500 μM) and calculated by the solution absorbance at 562 nm characteristic of $Cu^I(Bca)_2$ ($\varepsilon$ = 7,900 $cm^{-1}$ $M^{-1}$; log $\beta_2$ = 17.3).

(B) Biotin blot of cell-free conjugation activities. WT E2D2, or C21S, M38L, H55A, C85S, C107S and C111S mutants (5 μM) were incubated ± $CuCl_2$ (5 μM) for 1 h in the presence of GSH (200 μM), UBA1 (100 nM), $MgCl_2$ (5 mM), ATP (5 mM) and biotinylated-ubiquitin (2.5 μM) in Tris-HCl (50 mM, pH 7.5). Increased ubiquitin-tagging (25-100 kDa) indicates greater conjugation activity.

(C) Changes in solution visible spectra of $Cu^I(Fz)_2$ (55 μM; prepared from 55 μM $CuSO_4$, 300 μM Fz, 200 μM $NH_2OH$ and 100 μM ascorbate) in MOPS buffer (50 mM, pH 7.4) upon titration with E2D2 protein (C21S mutant; 0-14 μM).

(D) Full variation data of apoP, CuP and $Cu_2P$ $E2D2_{C21/85S}$ proportions upon increments of free $Cu^+$ (see Table S6 for details). The data for Cu:protein ratios of 0:1, 1:1 and 2.5:1 are shown in Figure 6C to facilitate visual comparison with the same ratios in Figure 6F. Inset, proposed scheme of facile $Cu^+$-exchange at the $C_{107}XXXC_{111}$ motif that shuttles E2D between three states (apoP, CuP, $Cu_2P$), described by two similar $K_{D1}$ and $K_{D2}$ (Figure 5I).

(E) Weighted average chemical shift differences ($\Delta\delta_{av}$) of $^{15}NH$ resonances between apoP and $Cu^+$-bound $E2D2_{C21/85S}$ (CuP) forms for resonances that could be assigned to the CuP state.

(F) Intensity changes for the resonances of the peaks assigned to apoP after adding $Cu^+$. Intensity differences were normalized for peak heights recorded in the absence of $Cu^+$ ($I_0$), after addition of

0.20 mM $Cu^+$ to 0.20 mM $E2D2_{C85/21S}$ (magenta), or 0.2 mM $Cu^+$ to 0.20 mM $E2D2_{C111S}$ (red). The region I106 to D112, flanked between the two blue dashed lines, encompasses the $C_{107}XXXC_{111}$ motif: the resonance intensities in this region showed prominent reduction in the presence of $Cu^+$ but could not be assigned for either apoP form or CuP form (despite most resonances of the apoP form in the absence of $Cu^+$ being assigned readily), consistent with rapid $Cu^+$ binding and exchange processes, as depicted by the inset Scheme in D. There was little change for the same region of the control $E2D2_{C111S}$ sample under the same condition.

(G) Conserved C111 of E2D2 is solvent accessible. (i) Left panel: structure model of dimeric E2D2-Ub-E3(MDM2) complex essential for p53 ubiquitination and degradation. The model was generated by superimposing the E2D2-Ub-MDM2 portion of the structure onto the MDMX structure of the determined crystal structure of E2D2-Ub-MDM2-MDMX (PDB: 5mnj). The latter complex was stabilized by mutation of E2D2-Cys85 to E2D2-Lys85 that formed a stable Lys85-Gly76(Ub) isopeptide bond to the Ub substrate. The residues surrounding Cys85 whose mutations have dramatic effect on discharge of substrate Ub from the E2D2-Ub-MDM2 complex are highlighted in sticks and labelled; Right panel: the schematic drawing of the dimeric complex on left. (ii) Left panel: a 180° rotation view of the above structure highlighting the identified fM affinity $Cu^+$-binding site (showed in orange sphere) consisting of two Cys thiols in the $C_{107}XXXC_{111}$ motif (in magenta); Right panel: the schematic drawing of the dimeric complex on left. The four Zn binding sites in the complex that are essential for the catalysis are shown in grey spheres. (iii) Left panel: an optimal view of the putative Cu(I) site consisting of the two sulfur atoms (shown in magenta sphere) in $C_{107}XXXC_{111}$ motif; Right panel: protein surface of the right half structure on left. Note Cys111 thiol is solvent accessible but Cys107 is completely buried by Ub (and by E2D2 itself as well, see below). (iv) Left panel: X-ray structure of E2D2-Ub complex with the structure of the E2D2 unit superimposing the E2D2 structure shown in (iii) (PDB: 3a33). The complex was stabilized by mutation of E2D2-Cys85 to E2D2-Ser85 that formed a stable peptide link Ser85-Gly76(Ub) to the ubiquitin substrate; Left panel: protein surface of the structure. Note Cys111 is solvent accessible but Cys107 is still partially buried by E2D2 itself.

### Figure S7

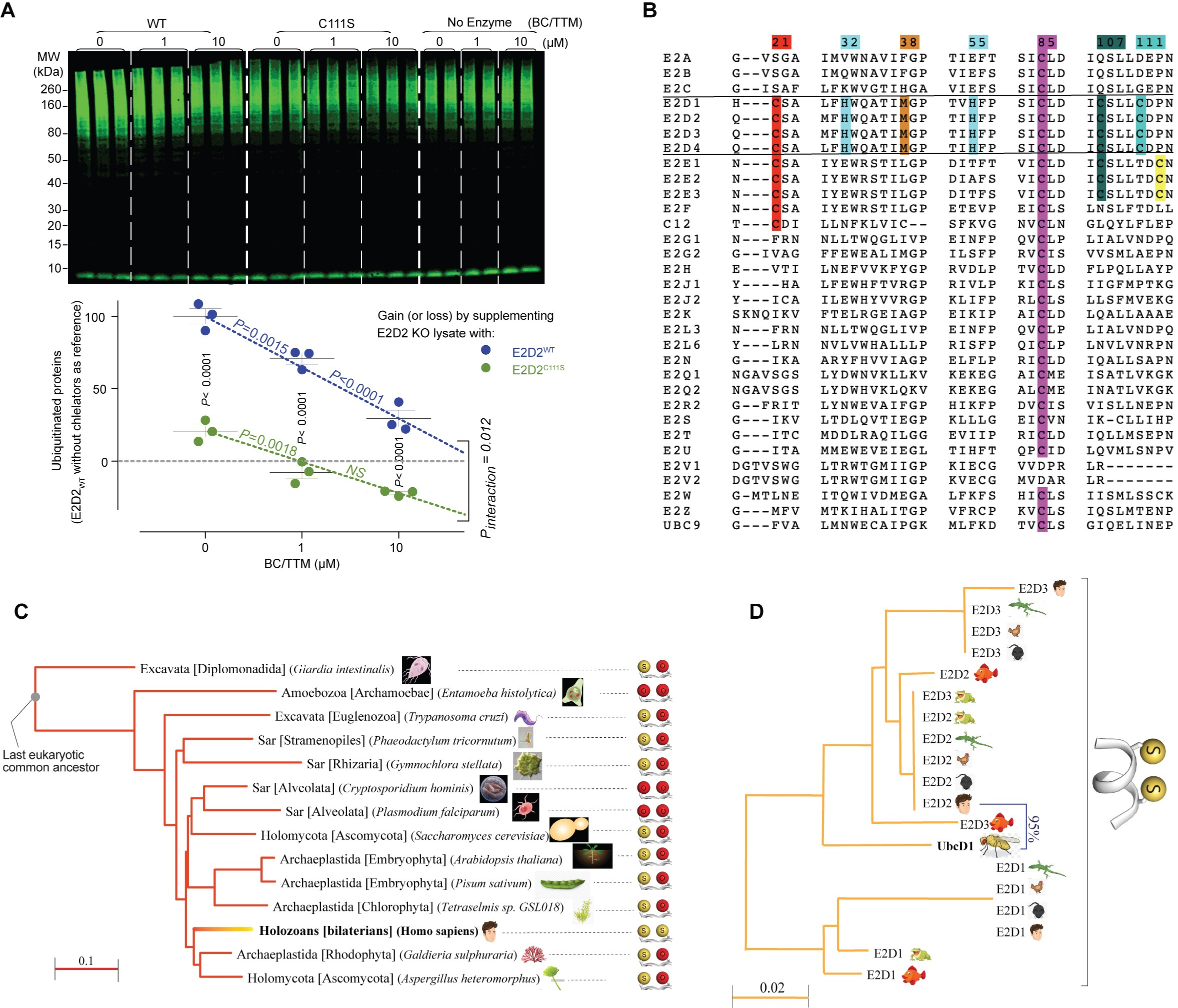
A
WT
C111S
No Enzyme
(BC/TTM)
0 1 10 0 1 10 0 1 10 (µM)
MW (kDa)
260
160
80
50
40
30
20
15
10
Ubiquitinated proteins (E2D2WT without chelators as reference)
100
50
0
P=0.0015
P<0.0001
P=0.0018
NS
P< 0.0001
P< 0.0001
P< 0.0001
P interaction = 0.012
Gain (or loss) by supplementing E2D2 KO lysate with:
E2D2WT
E2D2C111S
BC/TTM (µM)
B
21 32 38 55 85 107 111
E2A G--VSGA IMVWNAVIFGP TIEFT SICLD IQSLLDEPN
E2B G--VSGA IMQWNAVIFGP VIEFS SICLD IQSLLDEPN
E2C G--ISAF LFKWVGTIHGA VIEFS SICLD IQSLLGEPN
E2D1 H---CSA LFHWQATIMGP TVHFP SICLD ICSLLCDPN
E2D2 Q---CSA MFHWQATIMGP TIHFP SICLD ICSLLCDPN
E2D3 Q---CSA MFHWQATIMGP TIHFP SICLD ICSLLCDPN
E2D4 Q---CSA LFHWQATIMGP TIHFP SICLD ICSLLCDPN
E2E1 N---CSA IYEWRSTILGP DITFT VICLD ICSLLTDCN
E2E2 N---CSA IYEWRSTILGP DIAFS VICLD ICSLLTDCN
E2E3 N---CSA IYEWRSTILGP DITFS VICLD ICSLLTDCN
E2F N---CSA IYEWRSTILGP ETEVP EICLS LNSLFTDLL
C12 T---CDI LLNFKLVIC-- SFKVG NVCLN GLQYLFLEP
E2G1 N---FRN NLLTWQGLIVP EINFP QVCLP LIALVNDPQ
E2G2 G--IVAG FFEWEALIMGP ILSFP RVCIS VVSMLAEPN
E2H E---VTI LNEFVVKFYGP RVDLP TVCLD FLPQLLAYP
E2J1 Y----HA LFEWHFTVRGP RIVLP KICLS IIGFMPTKG
E2J2 Y---ICA ILEWHYVVRGP KLIFP RLCLS LLSFMVEKG
E2K SKNQIKV FTELRGEIAGP EIKIP AICLD LQALLAAAE
E2L3 N---FRN NLLTWQGLIVP EINFP QVCLP LIALVNDPQ
E2L6 Y---LRN NVLVWHALLLP RISFP QICLP LNVLVNRPN
E2N G---IKA ARYFHVVIAGP ELFLP RICLD IQALLSAPN
E2Q1 NGAVSGS LYDWNVKLLKV KEKEG AICME ISATLVKGK
E2Q2 NGAVSGS LYDWHVKLQKV KEKEG ALCME INATLVKGK
E2R2 G--FRIT LYNWEVAIFGP HIKFP DVCIS VISLLNEPN
E2S G---IKV LTDLQVTIEGP KLLLG EICVN IK-CLLIHP
E2T G---ITC MDDLRAQILGG EVIIP RICLD IQLLMSEPN
E2U G---ITA MMEWEVEIEGL TIHFT QPCID LQVMLSNPV
E2V1 DGTVSWG LTRWTGMIIGP KIECG VVDPR LR-------
E2V2 DGTVSWG LTRWTGMIIGP KVECG MVDAR LR-------
E2W G-MTLNE ITQWIVDMEGA LFKFS HICLS IISMLSSCK
E2Z G---MFV MTKIHALITGP VFRCP KVCLS IQSLMTENP
UBC9 G---FVA LMNWECAIPGK MLFKD TVCLS GIQELINEP
C
Excavata [Diplomonadida] (Giardia intestinalis)
Amoebozoa [Archamoebae] (Entamoeba histolytica)
Excavata [Euglenozoa] (Trypanosoma cruzi)
Sar [Stramenopiles] (Phaeodactylum tricornutum)
Sar [Rhizaria] (Gymnochlora stellata)
Sar [Alveolata] (Cryptosporidium hominis)
Sar [Alveolata] (Plasmodium falciparum)
Holomycota [Ascomycota] (Saccharomyces cerevisiae)
Archaeplastida [Embryophyta] (Arabidopsis thaliana)
Archaeplastida [Embryophyta] (Pisum sativum)
Archaeplastida [Chlorophyta] (Tetraselmis sp. GSL018)
Holozoans [bilaterians] (Homo sapiens)
Archaeplastida [Rhodophyta] (Galdieria sulphuraria)
Holomycota [Ascomycota] (Aspergillus heteromorphus)
Last eukaryotic common ancestor
0.1
D
E2D3
E2D3
E2D3
E2D3
E2D2
E2D3
E2D2
E2D2
E2D2
E2D2
E2D2
E2D3
UbcD1
95%
E2D1
E2D1
E2D1
E2D1
E2D1
E2D1
0.02

**Figure S7. Potential $Cu^+$ ligands across human E2s and evolutionary trees of E2D orthologues in eukaryotes and in vertebrates, related to Figure 7.**

(A) *In vitro* ubiquitination using cell extracts. Recombinant WT ($E2D2_{WT}$) and C111S-mutated ($E2D2_{C111S}$) E2D2 (1 μM) were incubated with cellular fractions from E2D2 knockout (KO) cells in the absence or presence of strong copper chelators (bathocuproine/ tetrathiomolybdate [BC/TTM] combination) for 1h. The formation of ubiquitinated proteins was detected by Western blot (P4D1 antibody). Activity of $E2D2_{WT}$ was more sensitive to copper chelation than the C111S mutant's. Graph summarizes data (n=3, normalised to $E2D2_{WT}$ without chelators), two-way ANOVA followed by Bonferroni-adjusted pairwise comparisons.

(B) Sequence alignment of potential $Cu^+$ ligands across human E2 conjugating enzymes. The Cys85 is the enzyme active site (magenta). Cys107 and Cys111 are highlighted in green. Other putative $Cu^+$ binding residues are highlighted.

(C) Evolutionary tree of E2D orthologs among eukaryotes. Representative species from major eukaryotic lineages were used for generating a Fast Minimum Evolution tree based on Grishin protein distance (scale, left-lower corner), with human E2D2 as reference. Side chain atoms of the $Cu^+$-binding ligands at position 107 (left) and 111 (right) are shown as spheres, coloured red/yellow for oxygen/sulfur, respectively, at Ser/Cys107 and at Thr/Cys111 (Asp111 in *G. Intestinalis*).

(D) Evolutionary tree of three E2D paralogues among vertebrates. Representative species from five vertebrate classes were used for generating a Fast Minimum Evolution tree based on Grishin protein distance (scale, left-lower corner), with their *D. melanogaster* orthologue UbcD1 as reference. Side chain atoms of the $Cu^+$-binding ligands at position 107 (left) and 111 (right) are shown as spheres, with all species harbouring cysteines' sulfur atoms at both 107 and 111 positions. Homology between human E2D2 and fly UbcD1 is denoted.

**Table S1. Ubiquitomics – MaxQuant parameters, MaxQuant output, filtering and annotation (Excel Table), related to Figures 3 and S3.**

Sheet 1- MaxQuant parameters. Parameters of the MaxQuant analysis are provided. Abbreviations: Acetyl (K), acetylation on lysine residue; GlyGly (K), ubiquitylation remnant on lysine residue after trypsin digestion; ppm, parts per million; Da, Daltons; PEP, Posterior Error Probability; Min, Minimum; Lfq, Label free quantitation.

Sheet 2- MaxQuant output. Protein Groups table, listing 1,563 MaxQuant outputs based on parameters in Table S1.

Sheet 3- Filtering. Following filtering to remove contaminants, reverse hits, proteins only identified by site and proteins with < 2 unique peptides, 912 unique proteins were considered for further analysis.

Sheets 4-6- Annotation and ID conversion. Following conversion of Uniprot mouse identifiers to their Ensembl protein IDs of their human orthologs (sheet 4), matching with original protein identifiers (sheet 5) and removing UBE2 enzymes, 882 Ub-associated proteins, considered as putative substrates for ubiquitination, were used all subsequent analysis (sheet 6, used as input for next stage). Light green-coloured cells represent protein IDs and amino acid sequences retrieved from STRING database, which correspond to protein IDs from MaxQuant output (pink-coloured cells).

**Table S2. Ubiquitomics – Quantifying relative abundance using Perseus (Excel Table), related to Figures 3 and S3.**

Sheet 1: Input, which includes protein identifiers (both Ensembl and Uniprot), protein characteristics (weight, cysteine, and methionine content) and LFQ intensities.

Sheet 2: Output. Data was transformed, scaled and normalized, followed by quantitative statistical analysis. Yellow columns: LFQ intensities for three experimental conditions (3 replicates per condition) following log-transformation and missing values imputation (the latter visualized in fig. S6B). Light blue columns: statistical analysis for differences in relative abundance for each individual protein, including -log p value and corresponding q-value for one-way ANOVA comparing the 3 experimental conditions and student's t-test difference between copper enriched (both concentrations grouped together, 6 columns in total) vs control conditions (3 columns). Orange columns: dichotomous categorization of proteins, based on whether i) ANOVA was significant (i.e. protein's abundance is significantly altered across media conditions, n=66, Visualised in Fig 2A); ii) ANOVA was significant AND *t*-test difference > 0 (protein is significantly more abundant in Cu-enriched media, n=48, green font) and iii) ANOVA was significant AND *t*-test difference < 0 (protein is significantly less abundant in Cu-enriched media, n=18, red font). Light green columns; protein IDs, names, length, mass and frequency of cysteine and methionine residues (log2 % out of total number of amino acids).

**Table S3. Ubiquitomics – Enriched ontology terms (Excel Table), related to Figures 3 and S3.**

Sheet 1: Perseus input, which includes same 882 Ub-associated proteins as Sheet 1 of Data File2. Protein identifiers (both Ensembl and Uniprot), protein characteristics (weight, cysteine and methionine content) and LFQ intensities are provided.

Sheet 2: Perseus output/Metascape input, which includes Ub-associated proteins over-abundant in $Cu^+$-supplemented (column A, $CuCl_2$ 10 μM, n=50; column B, $CuCl_2$ 25 μM, n=76) conditions relative to control, and those over-abundant in control conditions relative to either of the two $Cu^+$-supplemented conditions (column C, n=37 after accounting for duplicates), based on two independent-sample t-tests ($S_0$=2, FDR=0.05), visualised in volcano plots (Figure 3B).

Sheet 3: Metascape output-gene annotation, depicting annotation of genes according to list in which they appear (visualization in Circos plot, Figure S3F, is limited to proteins over-abundant in $Cu^+$-supplemented conditions, as there is no overlap with proteins over-abundant in control conditions).

Sheet 4: Metascape output-pathway enrichment based on all genes of input, depicting enriched pathways (pathways for which genes coding for proteins significantly altered by Cu treatment are significantly over-represented compared to genes coding for all 882 Ub-associated proteins captured in MS), alongside enrichment significance (per all genes of input rather than per list, raw and FDR-adjusted) and actual genes annotated to each pathway. Based on similarity, pathways were classified into three clusters.

Sheet 5: Metascape output-pathway enrichment based on individual input lists, depicting enriched pathways (pathways for which genes coding for proteins significantly altered by Cu treatment are significantly over-represented compared to genes coding for all 882 Ub-associated proteins captured in MS), alongside enrichment significance (based on each gene list separately and also based on all genes of input, followed by raw and FDR-adjusted) and actual genes annotated to each pathway (hits). Pathways are presented according to their cluster classification (sheet 4). Several highly significant pathways from each of two clusters that were enriched in genes coding for ubiquitin-associated proteins that were over-abundant in $Cu^{+}$-supplemented media and were selected by Metascape for network presentation (using Cytoscape), are highlighted in colours matching their cluster, as visualised in Figure 3C.

**Table S4. Ubiquitomics – The interactome of $Cu^{+}$-enhanced Ubiquitin-associated proteins (Excel Table), related to Figures 3 and S3-S4.**

Sheet 1: Protein-protein interaction data. Data were curated from STRING 10.5 Human Database, with the 48 Cu-enhanced protein identified above as the input and a combined score cutoff of 0.7 (high confidence). A total of 216 protein-protein interactions (PPIs) among 29 proteins were retrieved. For each PPI, interaction scores are provided in separate columns according to the following types of associations: experimentally-determined-interactions (such as co-purification and co-crystallization experiments), co-expression (proteins whose genes are observed to be correlated in expression, across a large number of experiments), phylogeny, homology, and database-annotated (using known metabolic pathways, protein complexes, signal transduction pathways, etc. from curated databases) and finally the combined interaction score. Unsupervised textmining (automated text-mining of the scientific literature) was excluded from the combined score calculation to reduce bias potentially arising from disproportional citations of some proteins.

Sheet 2: Network analysis. Using the Extract Connected Components option in the Network Analyzer app, a fully connected subnetwork of 27 proteins (nodes) and 215 PPIs (edges) was retrieved. The table lists graph-based node parameters calculated using Network Analyzer app across this subnetwork, as visualised in Figures 3D and S4A.

**Table S5. Ubiquitomics – Enrichment of $Cu^{+}$-enhanced proteins among predicted substrates of selected E2 clades (Excel Table), related to Figures 4 and S3-S4.**

*Sheets 1-3: Enrichment among predicted substrates of the E2D clade. Visualised in Fig. 2C.*

Sheet 1: With 886 proteins as input (882 potential substrates and four E2D enzymes), 17,340 PPIs with a combined score of 0.4 or above were identified using STRING 10.5 Human Database. For each PPI, interaction scores are provided in separate columns according to the following types of associations: experimentally-determined-interactions (such as co-purification and co-crystallization experiments), co-expression (proteins whose genes are observed to be correlated in expression, across a large number of experiments), phylogeny, homology, neighborhood and database-annotated (using known metabolic pathways, protein complexes, signal transduction pathways, etc. from curated databases), and finally the combined interaction score. Unsupervised textmining (automated text-mining of the scientific literature) was excluded from the combined score calculation in order to reduce bias potentially arising from disproportional citations of some proteins.

Sheet 2: Using Cytoscape, 70 of 882 Ub-associated proteins were identified as interacting with at least one member of the E2D clade (predicted E2D substrates). Of those, 24 (34%) were also among the 48 protein that were enhanced in copper-supplemented media.

Sheet 3: Using Cytoscape, 812 of 882 Ub-associated proteins were identified as not interacting with any member of the E2D clade (predicted E2D non-substrates). Of those, 24 (3%) were among the 48 protein that were enhanced in copper-supplemented media.

*Sheets 4-7: Enrichment among predicted substrates of the E2A/B clade and of both clades. Visualised in Fig S4G.*

Sheet 4: With 886 proteins as input (882 potential substrates and the E2A and E2B enzymes), 17,258 PPIs with a combined score of 0.4 or above were identified using STRING 10.5 Human Database1. For each PPI, interaction scores are provided in separate columns according to the types of associations outlined above. Unsupervised textmining (automated text-mining of the scientific literature) was excluded from the combined score calculation in order to reduce bias potentially arising from disproportional citations of some proteins.

Sheet 5: Using Cytoscape, 47 of 882 Ub-associated proteins were identified as interacting with E2A and/or E2B (predicted E2A/B substrates). Of those, 6 (13%) were also among the 48 protein that were enhanced in copper-supplemented media.

Sheet 6: Using Cytoscape, 835 of 882 Ub-associated proteins were identified as interacting with neither E2A nor E2B (predicted E2A/B non-substrates). Of those, 42 (5%) were among the 48 protein that were enhanced in copper-supplemented media.

Sheet 7: 32 proteins were predicted to be substrates for both an E2D and an E2A/B enzyme.

*Sheet 8: Effects of mass and redox-sensitive amino acids and of being an E2D or E2A/B substrate on $Cu^{+}$-enhanced ubiquitination. Visualised in Fig. 2D.* Multiple logistic regression model, output from SPSS with log2-mass, log2-cysteine/methionine content and being an E2D or E2A/B substrate as independent predictors and enhancement in Cu-supplemented media as the dichotomous outcome. n=882.

| Protein (P) | Total [Cu] (mM)[a] | Starting [GSH] (mM) | Remaining [GSH] (mM)[b] | Species Proportion (%)[c] | | | Remaining soluble P (%)[e] | [Cu] (μM) | | Free [GSH] (mM)[i] | Free $[Cu^+]$ (fM)[j] | Affinity (fM)[k] | |
|---|---|---|---|---|---|---|---|---|---|---|---|---|---|
| | | | | apoP | CuP[d] | $Cu_2P$[d] | | in P[g] | in GSH[g] | | | $K_{D1}$ | $K_{D2}$ |
| $E2D2_{C21/85S}$ | 0 | 1.0 | 1.0 | 100 | 0 | 0 | 100 | 0 | 0 | 1.0 | 0 | - | - |
| $E2D2_{C21/85S}$ | 0.10 | 1.0 | 0.91 | 74 | 24 | 2 | 97 | 56 | 44 | 0.84 | 0.15 | [l] | 0.45 |
| $E2D2_{C21/85S}$ | 0.20 | 1.0 | 0.82 | 59 | 32 | 9 | 99 | 101 | 99 | 0.67 | 0.25 | [l] | 0.46 |
| $E2D2_{C21/85S}$ | 0.30 | 1.5 | 1.23 | 62 | 27 | 11 | 104 | 96 | 204 | 0.92 | 0.19 | 0.50 | 0.44 |
| $E2D2_{C21/85S}$ | 0.40 | 1.5 | 1.14 | 50 | 33 | 17 | 99 | 135 | 265 | 0.74 | 0.28 | 0.54 | 0.42 |
| $E2D2_{C21/85S}$ | 0.50 | 1.5 | 1.05 | 37 | 35 | 28 | 101 | 182 | 318 | 0.57 | 0.43 | 0.53 | 0.46 |
| $E2D2_{C111S}$ | 0 | 1.0 | 1.0 | 100 | 0 | 0 | 100 | 0 | 0 | 1.0 | 0 | - | - |
| $E2D2_{C111S}$ | 0.20 | 1.0 | 0.82 | 77 | 23 | - | ~90[f] | <46[h] | >154 | <0.59 | >0.34 | - | - |
| $E2D2_{C111S}$ | 0.50 | 1.0 | 0.65 | 79 | 21 | - | ~60[f] | <42[h] | >458 | saturated | >1 | - | - |

**Table S6. NMR - Determination of $Cu^+$-binding affinity by protein speciation analysis of NMR shifts, related to Figures 6 and S6.**

Proteins (200 μM) were in 100 mM KPi buffer (pH 7.4) containing $NH_2OH$ (2.0 mM) with added GSH (Starting [GSH]).

[a] Added as $CuSO_4$ in pre-mixing with GSH and $NH_2OH$ (Ellman assay confirmed that ~90% $Cu^{2+}$ was reduced by GSH and ~10% by $NH_2OH$).

[b] Estimated remaining GSH based on ~90% added $Cu^{2+}$ being reduced by GSH[a]; Remaining GSH = Starting GSH – (oxidised GSH) – ($Cu^+$-complexed GSH). These estimates are made to ensure that there is abundant GSH remaining during acquisition of NMR data.

[c] Detected by NMR spectroscopy. For $E2D2_{C21/85S}$ the proportion of each protein species was inferred by summing the NH resonance peak intensities of all protein species for the well-resolved $Cu^+$-responsive resonances of N77 and L86 (shown in Fig. 5C) under the given condition, and then calculating the proportional distribution of each species averaged between N77 and L86. For $E2D2_{C111S}$, the estimation was based on L86 only, since N77 resonances did not alter in the presence of $Cu^+$.

[d] Proportions (based on the summed peak intensities at N77 and L86) of the apo-E2D2$_{C21/85S}$ (apoP) and the two resonance shifts induced in N77 and L86 by titrating in $Cu^+$ (from 0 to 2.5:1 $Cu^+$:protein). Two additional peaks for these residues appeared in response to increasing $Cu^+$. We assigned the first peak to appear as "CuP" and the second peak as "$Cu_2P$". The $Cu^+$-concentration-dependent increases in the proportions of the species are consistent with sequential binding of two $Cu^+$ ions to the same $Cu^+$-binding site, with comparable affinities.

[e] Estimated by summing the different NH resonance peak intensities of N77 and L86 in the presence of $Cu^+$ and then dividing by the peak intensity of the reference apoP in the absence of $Cu^+$. If the protein precipitates, these resonances decrease in intensity.

[f] Since the NMR resonances of the E2D2$_{C111S}$ samples with addition of $Cu^+$ lost intensity due to Cu-induced protein precipitation, the histogram of Fig. 5F was plotted with the proportions of the NMR-detected intensity peaks of the protein species (apoP, CuP) normalized to that of apoP with no added $Cu^+$.

[g] [Cu] in P = [CuP] + 2[$Cu_2P$]; [Cu] in GSH = total [Cu] – [Cu] in P.

[h] Estimated from the resolved $Cu^+$-responsive residue L86 only. Since the $Cu^+$-interaction appeared to be adventitious, the estimated value was a maximal limit.

[i] Free GSH = Remaining [GSH] – 1.5*[Cu] in GSH, since 1.5 GSH is required for each Cu in the $Cu_4(S\text{-}Cys)_6$ cluster.

[j] Calculated based on the equation 2 in Methods Section.

[k] Calculated based on $K_{D1} = [Cu^+]*[CuP]/[Cu_2P]$; $K_{D2} = [Cu^+]*[apoP]/[CuP]$.

[l] Cu occupancy is too low to allow a reliable estimation.

**Table S7. Evolution – Evolutionary tree of E2D orthologues among eukaryotes and among holozoans and of E2D paralogues among vertebrates (Excel Table), related to Figures 7 and S7.**

*Sheets 1-4: Evolutionary tree of E2D orthologues among eukaryotes, visualised in Figure S7C.*

Sheet 1: FASTA sequences of human E2D2 and 13 representative species from major eukaryotic lineages.

Sheet 2: With human E2D2 as reference, the summary table depicts for each accession (i.e., orthologue) the following:

- Scientific and common names of the species studies
- Taxonomy ID
- Max Score - the highest alignment score of a set of aligned segments from the same database sequence. The score is calculated from the sum of the match rewards and the mismatch, gap open and extend penalties independently for each segment. This normally gives the same sorting order as the E Value.
- Query Coverage - the percent of the query length that is included in the aligned segments).
- E(xpect) Value - the number of alignments expected by chance with a particular score or better. The expect value is the default sorting metric and normally gives the same sorting order as Max Score.
- % identity
- Protein length

Sheet 3: Data used to generate phylogenetic tree, presented in Abstract Syntax Notation (ASN) format.

Sheet 4: Sequence alignment map. Red squares depict deviations from reference (human E2D2) sequence. C107 and C111, corresponding to red and yellow spheres visualised in Figure S7C, are highlighted.

*Sheets 5-8: Evolutionary tree of E2D orthologs among holozoans, visualised in Figure 7A.*

Sheet 5: FASTA sequences of human E2D2 and 11 representative species from major holozoan (unicellular and multicellular) lineages, with an additional representative of holomycota.

Sheet 6: With human E2D2 as reference, the summary table depicts for each accession (i.e., orthologue) the information as in Sheet 2.

Sheet 7: Data used to generate phylogenetic tree, presented in Abstract Syntax Notation (ASN) format.

Sheet 8: Sequence alignment map. Red squares depict deviations from reference (human E2D2) sequence. C107 and C111, corresponding to red and yellow spheres visualised in Figure 6A, are highlighted.

*Sheets 9-12: Evolutionary tree of E2D paralogues among vertebrates, visualised in Figure S7D.*

Sheet 9: FASTA sequences of vertebrate E2D1-3 paralogues drawn from representative species from each of five major vertebrate lineages, alongside fly orthologue UbcD1.

Sheet 10: With *D. melanogaster* orthologue UbcD1 as reference, the summary table depicts for each accession the information as in Sheet 2.

Sheet 11: Data used to generate phylogenetic tree, presented in Abstract Syntax Notation (ASN) format.

Sheet 12: Sequence alignment map. Red squares depict deviations from reference (UbcD1) sequence. C107 and C111, corresponding to yellow spheres visualised in Figure S7D, are highlighted.
